# Previously hidden intraspecies dynamics underlie the apparent stability of two important skin microbiome species

**DOI:** 10.1101/2024.01.10.575018

**Authors:** Jacob S. Baker, Evan Qu, Christopher P. Mancuso, A. Delphine Tripp, Arolyn Conwill, Tami D. Lieberman

**Affiliations:** Institute for Medical Engineering and Sciences, Massachusetts Institute of Technology; Cambridge, MA 02139, USA; Department of Civil and Environmental Engineering, Massachusetts Institute of Technology; Cambridge, MA 02139, USA; Department of Systems Biology, Harvard University; Cambridge, MA 02138, USA; Broad Institute of MIT and Harvard; Cambridge, MA 02139, USA; Ragon Institute of MGH, MIT, and Harvard; Cambridge, MA 02139, USA

## Abstract

Adult human facial skin microbiomes are remarkably similar at the species-level, dominated by *Cutibacterium acnes* and *Staphylococcus epidermidis,* yet each person harbors a unique community of strains. Understanding how person-specific communities assemble is critical for designing microbiome-based therapies. Here, using 4,055 isolate genomes and 360 metagenomes, we reconstruct on-person evolutionary history to reveal on and between-person strain dynamics. We find that multiple cells are typically involved in transmission, indicating ample opportunity for migration. Despite this accessibility, family members share only some of their strains. *S. epidermidis* communities are dynamic, with each strain persisting for an average of only 2 years. *C. acnes* strains are more stable and have a higher colonization rate during the transition to an adult facial skin microbiome, suggesting this window could facilitate engraftment of therapeutic strains. These previously undetectable dynamics may influence the design of microbiome therapeutics and motivate the study of their effects on hosts.

## Introduction

Although human facial skin is constantly exposed to the diverse microorganisms of our surrounding environments, only select species and strains are able to colonize and persist^1–5^. Notably, two globally prevalent skin dwelling species, *Cutibacterium acnes* and *Staphylococcus epidermidis,* predominate in nearly every adult facial skin microbiome (FSM) sampled to date– together representing about ∼80% of the average individual’s relative abundance^4,6^. The universality of this composition across individuals and its simplicity relative to gut communities makes mechanistic study of this site’s ecology more easily attainable.

Despite commonality at the species level and the potential for transmission across individuals, each person’s facial skin microbiome is composed of a unique community of coexisting strains distinct from those found on other individuals^5,7,8^. This strain-level individuality indicates the presence of unknown ecological barriers that hinder the migration of new strains, as evidenced by the difficulty of achieving durable engraftment of topically applied *C. acnes*^9^. Determining the neutral^10^ or selective factors^11^ that create these unknown ecological barriers is of critical importance for developing durable probiotics, predicting the consequences of antibiotic interventions, and contextualizing the differences between healthy and diseased microbiomes^10,12^.

Measuring the dynamics of strain communities can identify the nature of these ecological barriers. For example, neutral (non-selective) priority effects should permit colonization only during early life or specific periods of disturbance^13^. On the other hand, selective ecological barriers to transmission can maintain person-specificity even when there is strain-level dynamism throughout life. Observations of strain transmission between individual people and the on-person dynamics of individual strains can differentiate neutral from selective ecological forces^14^.

When intraspecies diversity is high, as it is on the skin^15^, it can be difficult to decipher strain dynamics from metagenomics. In particular, closely-related strains can be indistinguishable with metagenomes, obscuring their distinct dynamics on and across people. Here, we use thousands of cultures from individual bacterial cells to obtain whole-genomes and overcome these limitations. This resolution enables us to cluster intraspecies populations into extremely closely-related clades– groups of isolates separated by so few mutations that they could have originated from diversification on a single person, which we refer to as lineages^16,17^. Cells from a lineage are typically separated by less than 100 point mutations across their genomes, reflecting their recent origin on an individual person, and are related by direct or indirect transmission^8,18–20^.

From a cohort of families we describe the dynamics of the FSM during the transition to adulthood and beyond, from the species to lineage-level. While birth and infancy have received extensive attention as critical periods for the assembly of human microbiomes^11,15,21–23^, the role of the period between infancy and adulthood in facilitating lineage colonization is underexplored. There is a well-documented transformation of the skin during this time, during which lipid secretion surges^24^ and bacterial density in the FSM increases more than 10,000-fold per cm^2^ and remains elevated throughout adulthood^25–28^. While this increase is largely driven by the anaerobic *C. acnes*, aerobes also increase 100 fold from early childhood to adulthood^28^. Using a combination of culture-based genomics and culture-independent metagenomics, we ask whether this transition is associated with the alleviation of priority effects and enhancement in new strain colonizations, and whether this change alters the stability of already colonizing lineages (Figure 1).

**Figure 1.**
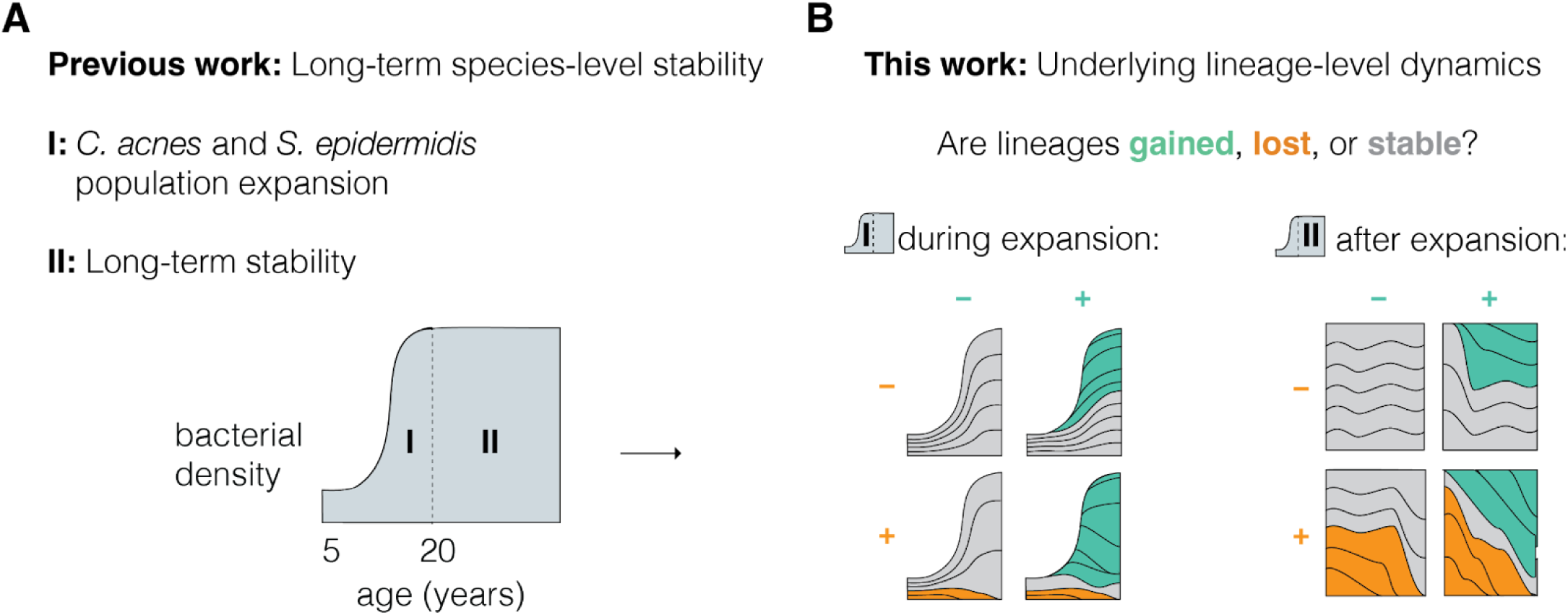
Unknown within-species facial skin microbiome dynamics at the lineage level. **(A)** Previous has shown that human development drives dramatic expansions of both *C. acnes* and *S. epidermidis* in l Skin Microbiomes (FSMs) during the transition to adulthood^28^, including a 10,000-fold increase in bundance of *C. acnes* and 100-fold increase in the abundance of *S. epidermidis* per cm^2^. The population of both species then remains high throughout adulthood. (**B**) It is known that many strains of both es coexist on the skin of each person^4^ In this work, we characterize the underlying turnover and migration dynamics of these populations by describing their membership at the lineage-level– the genomic ution needed to implicate direct sharing. The area between each set of lines represents the abundance of age.

We amassed a unique collection comprising 2,025 *S. epidermidis* and 2,030 *C. acnes* isolates from 57 individuals, supplemented with associated metagenomes. This extraordinary resolution and depth enables analyses traversing genomic levels of resolution from species down to individual genotypes. We reveal commonalities across species, including incomplete lineage sharing within families and primarily neutral on-person evolution. We also reveal species-specific dynamics with implications for the design of probiotic therapy: *S. epidermidis* lineages have high turnover throughout life, suggesting a selective barrier to transmission and a limit to the durability of natural probiotic strains. In contrast, *C. acnes* lineages have low turnover and have highest engraftment potential during the transition to adult-like FSM, consistent with alleviation of neutral priority effects during expansion.

### Species-level formation of the adult-type FSM

We collected facial skin microbiome samples from children and their parents at a single K-8 school across a 1.4 year period (Figure 2A). Individuals were sampled opportunistically on 5 different dates, and not every family member was sampled at the same time. In total, we sampled 30 children (aged 5 to 15) and 27 parents (aged 34 to 61). Basic health and demographic information was acquired via a questionnaire (Methods); given the modest size of our cohort, this information was primarily used for hypothesis generation.

**Figure 2.**
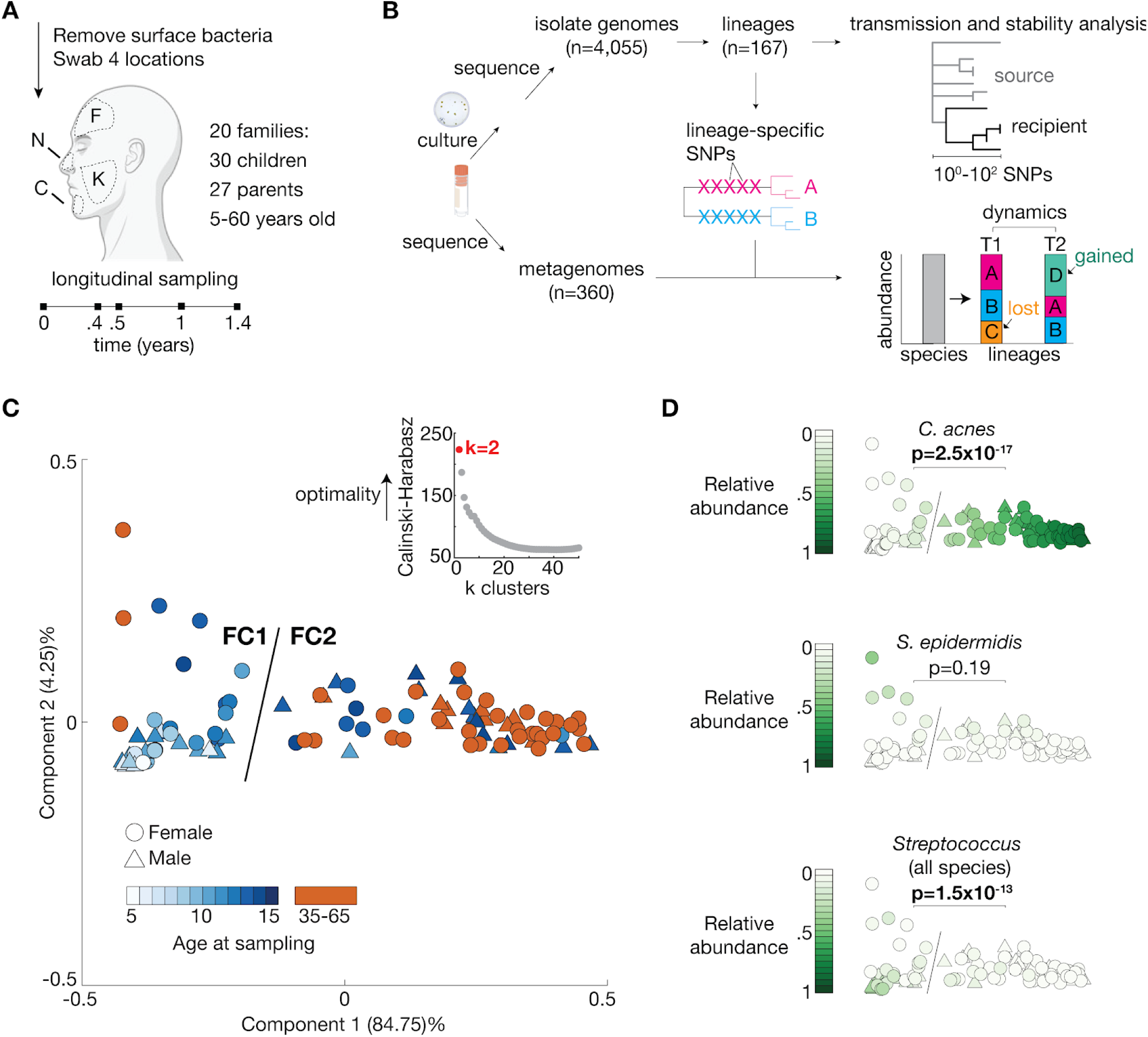
Metagenomic sequencing distinguishes between children who have and have not yet undergone the transition to a low-diversity adult-type FSM. **(A)** We collected samples of the FSM from four facial skin sites from 57 people aged 5-60 years from 20 distinct nuclear families. About half of these individuals were sampled once, and others were sampled at additional time points based on availability (Supplementary Table 1, Figure S2C). **(B)** From 8 of these families, we obtained 4,055 genomes (top) via a culture based approach and clustered them into 89 distinct *C. acnes* and 78 distinct *S. epidermidis* lineages. Concurrently (bottom) we sequenced metagenomic samples from all subjects and combined these reads with information obtained from isolate-defined lineages (lineage-specific SNPs) to measure lineage abundances and dynamics over time with higher sensitivity. Metagenomics also enabled analysis of species and coarse within-species diversity from subjects from which we did not perform culturing, as well as analysis from time points without cultured isolates. As we observe only 5 modest differences between facial sites (Figure S1), consistent with prior results ^16^, data from different facial sites were concatenated. We used metagenomics to detect transmission and turnover events and intra-lineage phylogenies (top right) to infer dynamics in further detail. **(C)** Hierarchical clustering of metagenomic data from 101 samples robustly separates two distinct types of species-level communities which we call ‘Facial Cutotypes’ (FCs): a low-diversity adult-type (FC2, n=61) and a subadult-type (FC1, n=40), consistent with expectations ^26,28^. Each dot represents one subject at one time point. These data are most optimally partitioned into two clusters (inset, Calinski-Harabasz score; Figure S2A). Points are colored by subject age, showing FC2 children are generally older than FC1 children. **(D)** As expected ^26,28^, the relative abundance of *C. acnes* is significantly higher in FC2 samples (‘adult-like’ FSM). The relative abundances of *S. epidermidis* is not significantly different between FCs, and the relative abundance of *Streptococcus* is significantly higher in FC1 samples. P-values show results of two-sided ranksum tests and are significant

Four facial locations were sampled from each subject using a pre-wet swab (forehead, nose, cheek, and chin). To limit collection of surface contaminants, each subject’s face was washed using a gentle cleansing cloth, rinsed with a saline wipe, and air dried. This process enabled repopulation of the surface from sebaceous secretions during the drying period^1,29^. From each sample, we profiled the entire community using metagenomic sequencing (mean filtered non-human species-assigned reads/sample = 3.1×10^6^, range 1.9×10^5^-22.2×10^6^, Methods), enabling study of both species-level and within-species diversity (Figure 2B). We found only minor compositional differences amongst facial sites (Figure S1), consistent with previous results^4,16^; we therefore grouped metagenomic reads across sites for each time point (herein called ‘samples’) to increase sensitivity to lowly abundant intraspecies groups.

Consistent with the expectation of dramatic changes to the skin during development^26,27^, younger children and adults have distinct species-level FSM communities. Hierarchical clustering of species-level communities revealed two clusters, herein called Facial Cutotypes (FCs) (Figures 2C and S2A-B, Methods). FC1 communities are found primarily on samples from younger children (median =11.4 years). Samples from nearly all parents (one exception, Figure S3A), and most older children, have FC2 communities (median age among samples from children =14.1 years). We classified children as FC2 if at least 1 of their samples is classified as FC2 (Figure S2C) and FC1 otherwise. While FC2 children are significantly older than FC1 children, (P=8.2×10^-4^, Figure S2D) there is not a clear boundary, consistent with individual differences in progression through Tanner stages^26^. It is possible that sampling more individuals might have revealed a continuum rather than a dichotomy; regardless, the clear separation between FC1 and FC2 found here enables us to clearly delineate which children have not yet begun developing an adult FSM from metagenomics alone.

The adult-type FC2 community is distinguished by *C. acnes* dominance and a corresponding loss of species-level alpha diversity (Figure S3B), as expected from previous results^28^. FC1 communities are dominated by various *Streptococcus* species. While *Staphylococcus epidermidis* is found at similar relative abundance in FC1 and FC2 communities (Figure 2D,6.4% vs 5.3%, P=0.19), the absolute abundance of *S. epidermidis* is significantly higher in parents than children (Figure S3F, P<0.05), highlighting shifts in microbial density during the transition to adulthood.

### Apparently neutral lineage-level coexistence on individuals

We next characterized intraspecies diversity and dynamics at the lineage level, the level required to understand migration dynamics. As determining lineage membership requires whole-genome resolution, we employed a high-throughput, culture-based approach. We focused on 8 family units for which we had samples from at least 3 subjects, each including at least one parent and one child. From each time point and individual, we obtained whole genomes from 1-133 *C. acnes* (median 51) and 1-186 *S. epidermidis* (median 51) single-colony isolates, derived using mildly selective growth conditions (Methods). As the number of cultured isolates varied across individuals, we used thresholds for inclusion of subjects in each analyses, listed throughout the text, figure legends, and in the Methods. The total of 4,055 isolates were clustered into 167 lineages, each with at least 3 isolates, using an approach that guarantees isolates within a lineage are more closely related to one another than any are to isolates from other lineages (Methods). While we observed evidence of gene content variation among isolates from the same lineage (Supplementary Table 4), we focus here on single-nucleotide polymorphisms for transmission-related clustering because of their clock-like accumulation over time. Isolates from the same lineage are separated by ≤ 62 and ≤ 90 SNPs across the whole genome for *C. acnes* and *S. epidermidis*, respectively (Figure 3A), reflecting a common ancestor within the lifespan of an individual person and therefore relationship through recent transmission^8^.

**Figure 3.**
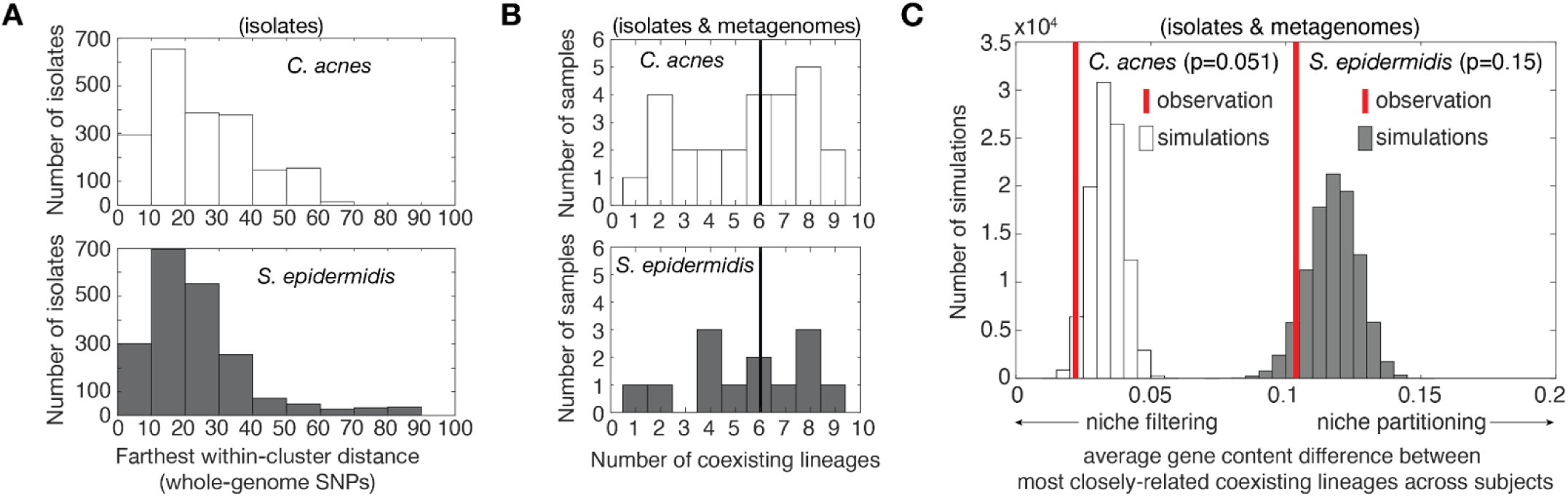
Closely-related lineages coexist on individuals. **(A)** Lineages are so closely related that the maximum within-lineage distance for either species is only 90 SNPs across the genome (90 SNPs for 2,025 *S. epidermidis* isolates, 62 SNPs for 2,030 *C. acnes* isolates) indicating a most recent common ancestor within the lifespan of an individual person. **(B)** For both species, we almost always observe multiple co-existing lineages at any given time point. For each sample, all lineages detected in either isolates or metagenomics are included (see Figure S4A-B, Methods). Only samples with both >25 isolates and >70% lineage-level metagenomics classified are included (n=26 for *C. acnes,* 13 for *S. epidermidis*). Vertical bars represent the median. **(C)** The average gene content difference between the most closely-related coexisting lineages on a subject are not more different than expected by chance for either species, as would be expected under metabolic niche partitioning. Coexisting lineages are the same as (B), and their gene content difference is measured from lineage co-assemblies (Methods). P-values represent twice the proportion of Monte Carlo simulations (1×10^5^ for each species) which are less than the observed value.

To better detect lowly abundant lineages missed by culturing but shared between subjects, we also assessed lineage composition using metagenomics (Methods). This also enabled us to query intraspecies diversity from subjects from whom no isolates were obtained, at coarser intraspecies resolutions. Metagenomic-inferred lineage abundances are significantly correlated with isolate-inferred abundances in subjects with sufficient data for both (Fig. 3B, Figure S4; P=5.1×10^-22^ and R^2^=0.55 for *C. acnes*; P=3.8×10^-18^ and R^2^=0.82 for *S. epidermidis*). Across all subjects, we find that multi-lineage colonization is common for both species, consistent with previous results^16,30^, with up to 9 lineages of a species coexisting on a person simultaneously.

To test if co-existing lineages of the same species fill distinct functional niches, we examined gene content and phylogenetic differences among co-existing lineages. If lineages coexist because they occupy distinct functions, we would expect to see overdispersion– a greater dissimilarity between pairs of lineages on a person than expected from a random sampling of lineages. For both species, we find no signal for overdispersion for either gene content or phylogenetic distance (Figure 3C and Figure S5), indicating that closely-related lineages are able to co-exist. While these results cannot rule out the possibility of niche partitioning driven by gene content below the limit of detection, these results trend closer to underdispersion than overdispersion and are consistent with a neutral model of lineage coexistence^31^ for *C. acnes* and *S. epidermidis* on facial skin.

### Family members share some, but not all of their lineages

Lineages are extensively shared within, but not between families, as expected from studies in the gut and oral microbiome^20^. Using metagenomic data to detect potential low-level sharing of lineages, we find 44% and 46% of the detected *C. acnes* and *S. epidermidis* lineages were shared among two or more people in a family (Methods). In contrast, only 2 *C. acnes* lineages are shared between non-family members at the school in metagenomics samples (and 0 for *S. epidermidis*). In both of these cases of between-family sharing, the lineages are found at low (<5%) abundance on one of the two individuals implicated, potentially indicating experimental or computational error. An additional 4 *C. acnes and 8 S. epidermidis* lineages had any isolates from more than one family; however, these sharing events were not able to be corroborated in metagenomes. While each of these instances involves students from the same school and it is plausible that they could indicate transient transmission, experimental error cannot be ruled out due to the small numbers of isolates involved (Methods, Figure S6). Together, these results indicate that lineage sharing is more likely between those in close contact.

It is notable that family members retain unique communities despite this potential for exposure-dependent sharing (Figure 4A, S7). Notably, the majority of lineages on parents are not shared between both parents (23/35 of *C. acnes* and 23/24 of *S. epidermidis* lineages). Subject-specific lineages can often be found at high abundance (Figure S8B), suggesting this is not just an artifact of detection limits or limited potential for exposure. This incomplete sharing suggests the presence of transmission barriers on adults that are either neutral (e.g. priority effects^32^) or selective (e.g. person-specific selection), with higher barriers for *S. epidermidis*.

**Figure 4.**
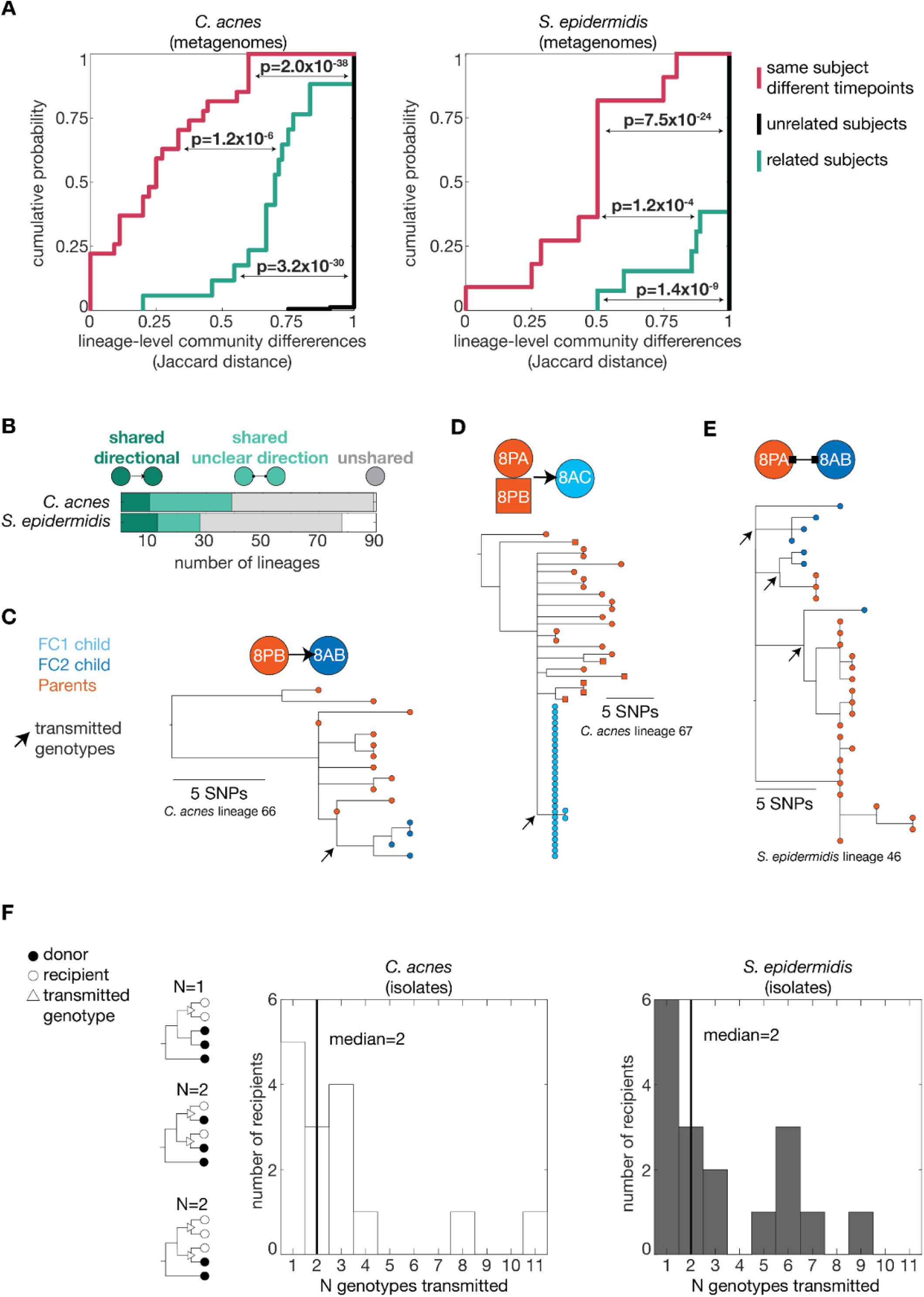
Not all lineages are shared between family members despite low physical barriers to transmission. **(A)** For both species, family members share some, but not all, of their lineages. Lineage membership was determined using metagenomics samples with >70% lineage-level assignment. Jaccard distance was chosen because this metric is not sensitive to differences in abundance, and values were calculated using all lineages found across samples for each subject. Distribution of within-family (green, n=17 for *C. acnes,* 13 for *S. epidermidis*) Jaccard distances is significantly lower than different time points from the same subject (magenta, n=27 for *C. acnes,* 11 for *S. epidermidis*). Sharing between non-family members from the same school (gray, n=157 for *C. acnes,* 92 for *S. epidermidis*) is exceedingly rare. **(B)** When lineages (89 for *C. acnes*, 78 for *S. epidermidis*) are shared between people (shades of green), the directionality is not always clear, but tree topologies indicate that multiple cell transmissions are common. **(C-E)** Three example phylogenies from the same family are shown, with the minimum inferred transmitted genotypes to the focal recipient indicated by arrows. Phylogeny **(C)** shows a clear transmission from 8PB to their child 8AB. Phylogeny **(D)** shows a single-cell bottleneck to child 8AC, but which parent this came from is unclear. Phylogeny **(E)** shows multiple transmission between 8PA and 8AB, the directionality of which is unclear. (**F)** Across all lineages and recipients, the minimum number of transmitted genotypes required to explain the diversity of lineages is usually greater than 1 (n=15 for *C. acnes*, 17 for *S. epidermidis*, median 2 for both species, vertical bars), suggesting the migration of multiple cells at one or multiple time points. Lineages shared by three or more individuals are counted once for each recipient. P-values represent the result of two-tailed ranksum tests and are all significant at a false discovery rate of 5% using the Benjamin-Hochberg procedure.

### Lineage sharing is mediated by multi-cell transmission

To understand the directionality and number of cells involved in successful transmissions, we turned to isolate-based lineage phylogenies (Figure 4B-F). Directionality, as evidenced by an individual’s isolates having a younger most recent common ancestor (MRCA) than that of another individual’s isolates, was inferrable for only a minority of lineages with sufficient isolates (Figure 4B,S9A, Methods). A parent was the donor in the majority of inferred directional transmissions (Figure 4C-D, S9B-C, 9/10 for *C. acnes* and 9/13 for *S. epidermidis*). Complex topologies from which directionality could not be inferred emerged from either the involvement of more than two individuals, the transmission of multiple genotypes, or both (Figure 4E, S9D).

Multi-cell transmission is common for both species when lineages are shared between family members. We estimate a median of at least two transmitted cells per recipient per lineage for both species (up to 11 transmitted genotypes, Fig 4F, Methods). These estimates are conservative, as multi-cell transmission of isogenic genotypes cannot be detected and the number of transmitted genotypes increases with the number of recovered colonies for *C. acnes* (Fig S10A). While our approach cannot distinguish if cells are transmitted simultaneously or at distinct time points, these results show that single-cell bottlenecks are rare during between-person transmission for both species.

The predominance of multi-cell transmission in *C. acnes* is consistent with our previous work indicating that each sebaceous follicle is independently colonized, creating an archipelago-like community structure which maintains coexisting genotypes by preventing direct competition^16^. It is unclear if there is an anatomical basis for the coexistence of multiple transmitted *S. epidermidis* genotypes, given the unknown fine-scale spatial structure of *S. epidermidis* populations on human skin^29,30^. *C. acnes* and *S. epidermidis* lineages have significantly different phylogenetic topologies, with *C. acnes* having a more ‘comb-like’ structure with a higher proportion of unshared mutations than *S. epidermidis* (Figure S9E), which could indicate that populations of *S. epidermidis* are more efficiently mixed. Regardless, the absence of single-celled bottlenecks for either species suggests the possibility of a spatially structured environment that facilitates stable coexistence of multiple transmitted genotypes.

### *S. epidermidis* lineages are continuously acquired and increasingly stable over time

We next turned to understand the dynamics of lineages on individual people, first by comparing sequential samplings. For *S. epidermidis,* we observe many turnover events of lineages within the short time windows assessed here (max interval length = 1.4 years). A significantly higher proportion of lineages and total abundance of lineage-level communities are lost across time points in *S. epidermidis* when compared to the stable *C. acnes.* (Figure 5A-B). This dynamism can also be seen as rapid changes at the community level, though this does not extend to complete turnover within any time window for any given subject (Figure 5C, S11). While *S. epidermidis* has been previously reported to be stable at this timescale^5^, a reanalysis of results from that study provides further support for a model of continual *S. epidermidis* turnover, with similar rates of strain loss over time as obtained here (Figure S12). We observed similar fluctuations in both parents and children (Figure S11 and S13), suggesting that *S. epidermidis* lineages are continually acquired and lost throughout life stages.

**Figure 5:**
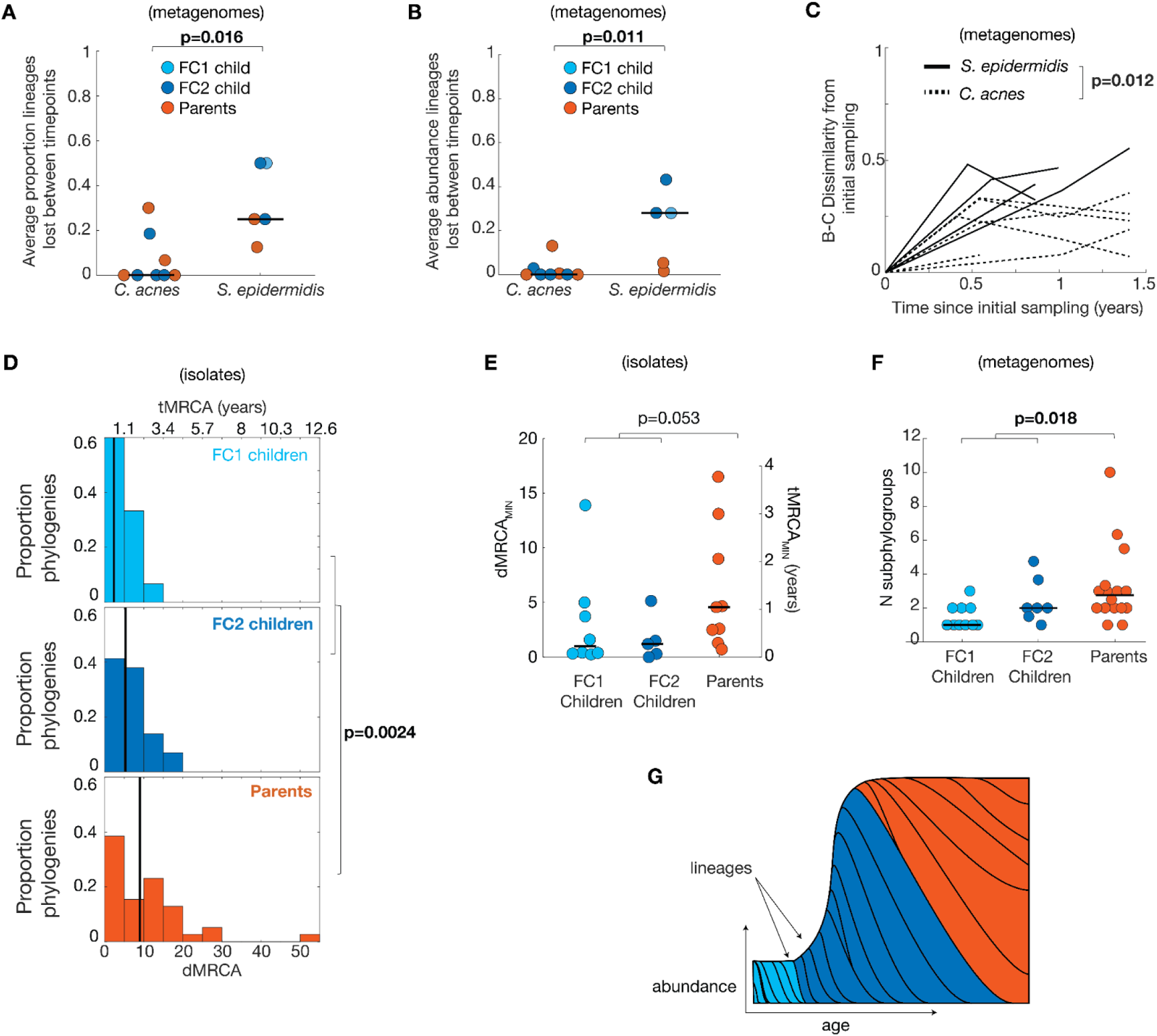
Lineages of *S. epidermidis* are gained and lost throughout life. We examined lineage-level stability in subjects with sufficient metagenomic coverage across multiple time points. (**A**) *S. epidermidis* is much less stable than *C. acnes* at the lineage level. (**B**) As lowly-abundant lineages near the detection limit might create a false appearance of turnover, we assessed the proportion of abundance lost between time points; total abundance of lineages lost between time points is substantial for *S. epidermidis* and negligible for *C. acnes.* The average values across all time point intervals for each subject is shown (n=8 subjects for *C. acnes,* 5 for *S. epidermidis*). **(C)** During our observation window of 1.4 years, individuals’ *S. epidermidis* lineage-level communities turn over faster than those of *C. acnes*, as assessed by Bray-Curtis dissimilarity, a metric that accounts for abundance. Subjects with >70% assignment of metagenomics at the lineage level at least two time points and across at least 6 months are shown (n=7 subjects for *C. acnes,* 4 for *S. epidermidis,* see Figure S11 for analysis of more subjects at higher taxonomic levels and Figure S12 for similar results from the reanalysis of ^5^. **(D)** Instability is also supported by analysis of inferred colonization times as assessed by mutational distance to the most recent common ancestor on each person (dMRCA), which we convert to time to the most recent common ancestor (tMRCA) using a molecular clock (Figure S14). Lineages were analyzed for subject dMRCA if they had at least 5 isolates derived from that subject. The median age of *S. epidermidis* lineages on individuals is ∼1.2 years (n=83, max 12.2 years). Lineages on parents are significantly older than those on children, while still relatively young compared to adult life spans (see also Fig. S15). These age estimates are not an artifact of sampling depth, as lineages with many isolates often have low dMRCAs (Figure S17B-C). (**E**) The most recently colonized lineage per-subject (tMRCA MIN) is similar across groups (n=8 FC1 Children, 5 FC2 children, 9 Parents), suggesting similar rates of new colonizations. **(F)** Consistent with higher residence time and similar colonization rates, intraspecies richness is higher on parents than children. Intraspecies richness is assessed by the number of coexisting sub-phylotypes of *S. epidermidis*, in order to include subjects without lineage data, (n=10 FC1 Children, 7 FC2 children, 16 Parents; The average number of sub-phylotypes across time points are shown for each subject. Time point samples were only analyzed if they had > 70% assignment at sub-phylotype level. **(G)** Our observations suggest a model in which lineages of *S. epidermidis* are continuously gained and lost, but persist for longer on adults, who have higher carrying capacity of *S. epidermidis* cells ^28^. The area between each set of lines represents the abundance of a lineage, and each lineage is colored by the time window in which it was acquired. All P-values show results of two-sided ranksum tests and are bolded if significant at a false discovery rate of 5% using the Benjamin-Hochberg procedure.

To confirm our inferences of *S. epidermidis* lineage-level instability using an orthogonal method, we analyzed the age of the MRCAs (most recent common ancestors) of each lineage on each person using within-lineage phylogenies of isolate genomes. We first measured the rate at which *S. epidermidis* accumulates mutations, obtaining a value of 4.5 mutations/genome/year (Methods, Figure S14), in line with literature estimates^17,33–35^. We then used this rate to convert each lineage’s dMRCA (distance to MRCA on a subject) to its tMRCA (time since MRCA).

As expected from the instability seen in longitudinal data, all values of tMRCA were significantly smaller than their subjects’ ages, with a maximum value of tMRCA of 12.2 years and a median tMRCA of 1.2 years (Figure 5D, S15A). While low tMRCAs could alternatively reflect a recent selective sweep or neutral bottleneck that purged diversity^17^, we do not find evidence of frequent adaptive evolution, as indicated by parallel evolution, ratios of nonsynonymous to synonymous mutations, or sweeps between time points (Figure S16). Low values of tMRCA are found even in lineages with many isolates (Figure S17B). Together, these results indicate that *S. epidermidis* lineages generally persist on individuals for only a matter of years, despite apparent stability at the species level.

Inferred tMRCAs on parents were significantly longer than those on children (Figure 5D, median of 2.1 for parents vs. 0.52 and 1.2 years for FC1 and FC2 children, respectively). We note that this is not a trivial result from colonization at birth, as tMRCAs on children are still younger than their age. Thus, despite the capacity for constant turnover, each *S. epidermidis* lineage colonization is generally more stable on parents. To understand how the rate of migration of new lineages varies across ages, we investigated the lowest tMRCA across lineages on a given subject (tMRCA_min_), a value that reflects the age of the most recently acquired lineage on a subject. We find no significant difference in tMRCA_min_ across groups (Figure 5E), consistent with a model in which *S. epidermidis* lineages arrive on subjects of all ages at similar rates, with enhanced stability of individual lineages on parents.

These two results from tMRCA analyses– enhanced stability of lineages on parents and no difference in the acquisition of new lineages– together predict that more lineages coexist on adults than children at any given time. Indeed, we find a significant enrichment of intraspecies richness on parents (Figure 5F; Figure S18). These results are consistent with a model in which the increased *S. epidermidis* carrying capacity (absolute abundance) on adults relative to children^28^ (Fig. S3D) reduces drift and therefore enhances stability. Together, our results from independent approaches –longitudinal studies with metagenomics and phylogenetic inferences from isolates–indicate that *S. epidermidis* lineages are acquired and lost from at least childhood through adulthood, with longer persistence of individual lineages on adults (Figure 5G). This lineage-level dynamism implies that life-long priority effects cannot explain observations of incomplete sharing between family members (Figure 4A).

### *C. acnes* is readily acquired during population expansion

In contrast to *S. epidermidis*, *C. acnes* lineages were remarkably stable across individuals during the time intervals studied (Figure 5C, P=0.0061, Methods), consistent with prior observations from metagenomics^5^ (Figure S12). Lineages with turnover had lower confidence at detected time points than stable lineages (Figure S19), suggesting these lineages may appear gained or lost due to fluctuations near the limit of detection. *C. acnes* communities remained more stable than *S. epidermidis* communities when examining all subjects (including those without isolates) using the broader taxonomic level of phylotypes (Figure S11 and S13). This observation, despite the added statistical power from the larger number of phylotypes in *C. acnes* (9 vs 4, Figure S20), furthers evidence for the durability of *C. acnes* colonizations over time.

Despite this stability, *C. acnes’* lineage dMRCAs did not significantly correlate with host age or FC type (Figure 6A), and we were unable to identify a significant molecular clock signal to infer tMRCAs. The absence of a correlation between dMRCA and age could theoretically emerge from on-person selective sweeps, which purge accumulated diversity. However, we observe no evidence of selection in *C. acnes* (Figure S16), consistent with prior results from our group^16^. The consistency of dMRCAs across the lifespan may be due to recently co-transmitted *C. acnes* genotypes causing lineages to appear as old on recipients as their sources (Figure 6B); in cases of multi-cell transmission, the MRCA of a recipient’s lineage diversity could have emerged on the source individual, potentially even before a subject was born. Accordingly, we find deeply branching clades coexisting on young individuals (Figure 6B, inset) and no significant difference in dMRCA_max_ between children and parents (Figure 6B). Multi-clade transmission may also play a role in preventing a significant molecular clock signal for *C. acnes* here (Figure S14) and in previous data^16^ for converting values of dMRCA to tMRCA.

**Figure 6.**
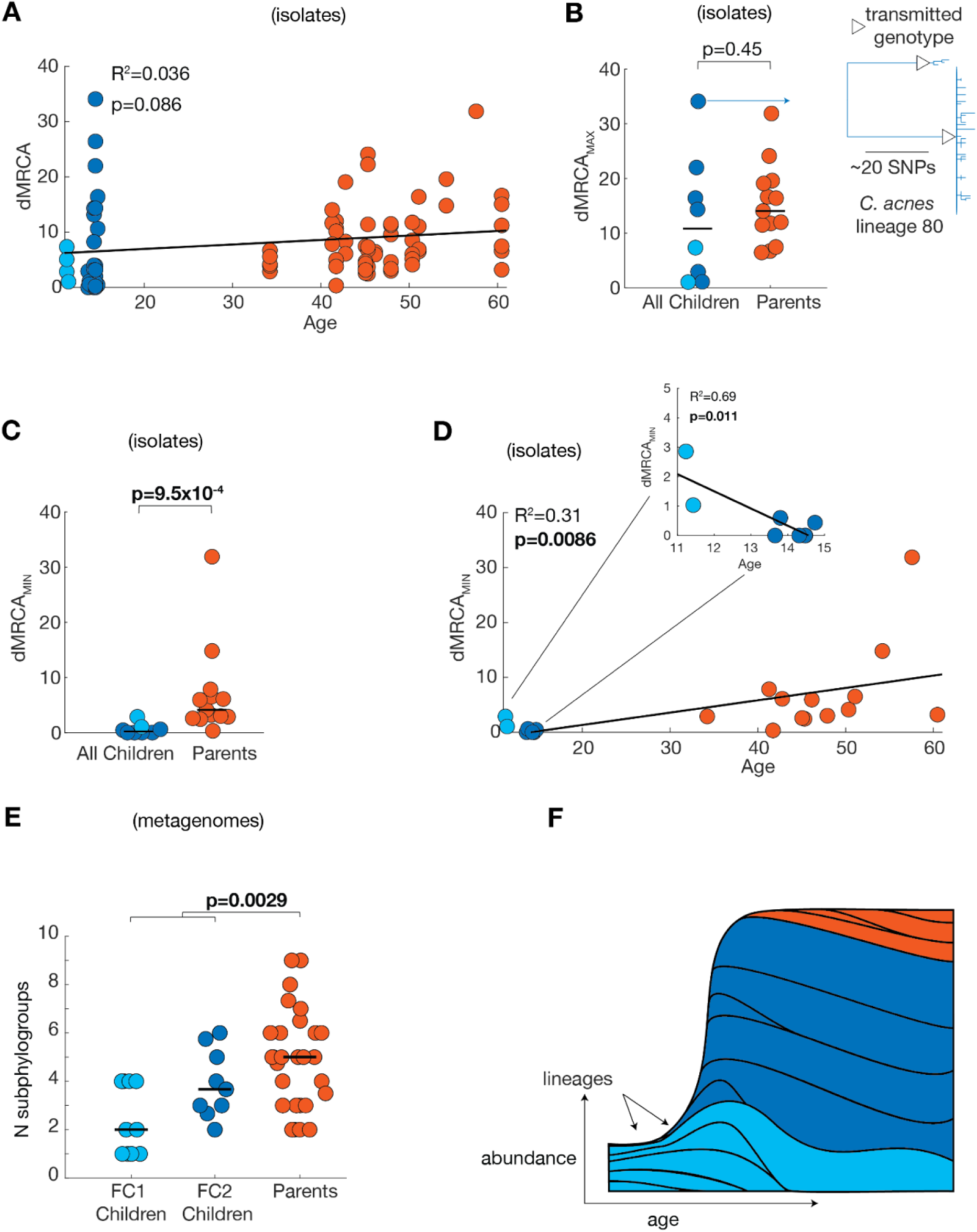
Rapid acquisition of *C. acnes* lineages during cutotype transition suggests population expansion temporarily alleviates ecological barriers to colonization. **(A)** The dMRCAs of *C. acnes* lineages are not significantly correlated with subject classes or ages (n=83), unlike for *S. epidermidis* (Fig. 5D, S15A). Lineages were analyzed for subject dMRCA analyses if they had at least 5 isolates derived from that subject. **(B)** Moreover, the lineages with the longest dMRCA per subject (dMRCA _MAX_) are not significantly different between parents and children (n=2 FC1 Children, 6 FC2 Children, 13 Parents). This lack of signal is consistent with the observation of multi-cell transmission of lineages (Figure 4F); Inset shows the most extreme lineage phylogeny of isolates from a child. **(C)** Regardless, we find an age-based signal when considering the dMRCA of the most recently colonized lineage in each subject, dMRCA _MIN_. Parents’ most recently colonized lineages have larger values of dMRCA _MIN_ than those in all children, indicating less frequent colonizations over time. (**D)** Consistently, there is a significant correlation between dMRCA MIN with age, with a notable exception among children (inset): older children have more recently colonized lineages than younger ones, indicating a window period of rapid acquisition. **(F)** Supporting an accumulation of new lineages of *C. acnes* over time, in our larger cohort without lineage-level data, we find a larger number of coexisting sub-phylotype in adults (n=8 FC1 Children, 10 FC2 Children, 26 Parents; The average number of sub-phylotypes across time points are shown for each subject, and time point samples are only considered if they have >70% assigned at the sub-phylotype level. **(G)** Together, these results are consistent with a model in which lineages are acquired most readily during the transition to an adult FSM, alongside population expansio n. The area between each set of lines represents the abundance of a lineage, and each lineage is colored by the time window in which it was acquired. All correlation coefficients and P-values are for linear (Pearson) correlations or two-sided ranksum tests and are bolded if significant at a false discovery rate of 5% using the Benjamini-Hochberg procedure

We therefore turned to dMRCA_min_ to assess the age of the most recently acquired simple lineage acquisition per subject. Across all subjects, children had significantly lower values of dMRCA_min_ than parents (p=9.5×10^-4^, Figure 6C), consistent with more recent acquisitions. Moreover, *C. acnes* dMRCA_min_ was significantly correlated with age (Figure 6D; R^2^=0.31, p<0.01). When considering only child subjects, dMRCA_min_ was lower on FC2 subjects and negatively correlated with age (p=0.011; Figure 6D inset). Together, these trends support a model in which *C. acnes* lineages are acquired at a higher rate during the transition to FC2, concurrent with the increase in the size of the *C. acnes* population.

This influx of new lineages during the transition to an FC2 community could result in either replacement or addition to preexisting lineages. Consistent with a model of addition, and with an increased carrying capacity, the number of sub-phylotypes is higher on parents than on children (Figure 6E). While we were unable to obtain statistically significant assessments of *C. acnes* within-species diversity between FC1 and FC2 children due to the low abundance of *C. acnes* on FC1 children (Figure S18, S2E), we observe trends in agreement with a model of *C. acnes* lineage addition during the transition to the FC2 cutotype (Figure 6F).

Altogether, these results demonstrate distinct on-person dynamics for *C. acnes* and *S. epidermidis* lineages. The rate of *S. epidermidis* lineage turnover is higher throughout the lifespan. While we cannot determine the basis of this difference, it is interesting to note that *S. epidermidis* lineages have a larger and more dynamic accessory genome^5,30^ (Figure S21) and may therefore experience more between-lineage competition. Another notable difference is that only *C. acnes* lineages appear to be most rapidly gained specifically during expansion and the transition to an adult-like facial skin microbiome. The more dramatic changes to *C. acnes* population size during this timespan (10^5^-fold vs. 10^2^-fold increase)^28^ may be responsible for this difference.

## Discussion

Here, we reveal previously hidden lineage-level assembly dynamics and report distinct models of community assembly for the two most dominant bacteria of the sebaceous skin microbiome, with implications for therapeutic design.

We find an enhanced rate of *C. acnes* lineage colonization during the transition to an adult facial skin type (Fig 6), consistent with a model of neutral ecology and diminished drift barriers due to a dramatic increase in population size. This result suggests that probiotic *C. acnes* strains^9,36^ may more readily engraft long-term if applied during this life stage.

In contrast, *S. epidermidis* lineages exhibit similar rates of colonization throughout the lifespan, and have high turnover (Figure 5). The lack of homogenization among parents for *S. epidermidis,* despite this high invasion potential, suggests the presence of selective engraftment barriers. These results suggest that natural probiotic strains of *S. epidermidis* will persist for a few years at most (Fig 5D).

Despite these differences, we report commonalities across species. Lineage-sharing is common between parents and children, rare between unrelated members at the same school, and characterized by multi-cell transmission (Figure 4F). Despite ample opportunity, family members do not share all of their lineages for either species (Fig 4A). While previous works have revealed person-specificity of the skin microbiome at the strain level^4,30^, this is the first to report that skin microbiome person-specificity extends even within families. Lastly, neutral forces dominate on-person SNP evolution for both species (Fig S16). Future work will be needed to understand which of these commonalities also extend other colonizers of human skin and other human microbiomes. Incomplete strain sharing between family members has also been shown in the gut^37,38^ and oral microbiomes^20,39^, mirroring these results. In contrast, strong on-person adaptive evolution has been shown in the gut microbiome, though we find no evidence for these two skin species (Fig S16). The generality of multi-cell transmission cannot yet be assessed, as this is the first work conducted at the resolution required to address clonality of lineage transmission.

For *C. acnes*, the finding of incomplete sharing can potentially be rationalized in terms of its biology. *C. acnes* growth occurs primarily within sebaceous follicles (pores), and the population within each pore is dominated by the descendants of a neutral single cell bottleneck event^16^. Pores which briefly become accessible to colonization are more likely to be seeded by lineages that are abundant on that person, rather than lowly-abundant migrating lineages^31^. In our model, colonization by an entirely new lineage of *C. acnes* is rare, and incomplete sharing is a neutral outcome of the limited rate at which new strains engraft (which increases during the transition to adulthood). Whether or not similar local priority effects can be invoked for *S. epidermidis* populations is unclear; however, the potential for rapid turnover seen here (Fig. 5) suggests that any such priority effects are short lived. While transmission has been studied across diverse body sites for *S. epidermidis*^30^, showing rapid lineage transmission across the body, the fine-scaled spatial structure and local niche remain unknown.

The primacy of adolescence for *C. acnes* lineage acquisition can similarly be explained by known forces. During adolescence, increased sebum production^24^ supports a larger *C. acnes* population on the surface^28^, with a 10^5^-fold increase in measured colony-forming units per area^28^. This increased carrying capacity on the surface could increase the likelihood that a newly opened pore is colonized by a migrating *C. acnes* lineage– and thereby alleviating priority effects. In addition, other factors associated with increased carrying capacity, such as a higher fraction of pores accessible to colonization, could increase the likelihood of lineage establishment^40–42^. Regardless of the precise mechanism, the importance of this transitional period for *C. acnes* strain acquisition highlights the critical role of life stages beyond birth and infancy for community assembly.

Adolescence may be a weaker force for *S. epidermidis* colonization than for *C. acnes* because the change in absolute abundance during this time window is relatively marginal (10^2^-fold vs. 10^5^-fold, respectively)^28^, but other factors may be at play. More work will be needed to understand forces that shape *S. epidermidis* person-specificity at the lineage level despite the potential for continual turnover. In particular, we do not explore here the rule of interbacterial interactions through bacteriocins or small molecules^43–46^, nor the role of host genetics and immunity– factors which are likely to affect both transmission and turnover. As these traits are difficult to completely predict with genomic data alone (e.g. though some antimicrobials are known^44,47^, there are likely more^48,49^), future work with deep phenotyping may be required.

Both of these species have been considered as potential probiotics^9,50^, and we anticipate that the dynamics described here will be helpful in the design of therapies involving these species. Notably, the lower relative abundance of *C. acnes* on skin with acne vulgaris^51,52^, psoriasis^53,54^, and active atopic dermatitis flares^55^ relative to healthy skin raises the possibility that this species or some strains thereof^52,56,57^ may provide a benefit to the host^58^. If some or all *C. acnes* are beneficial for treating disease or providing colonization resistance to infection^44^, the openness of the skin microbiome to *C. acnes* colonization during adolescence suggests that probiotics applied during this time window may be more likely to result in long-lasting efficacy. Finally, it is tempting to contemplate whether recently acquired lineages of either species interact with the host immune system differently, and we anticipate that future work will explore whether the lineage acquisitions and turnovers described here are consequential for acne vulgaris or other diseases.

## Acknowledgments

We thank the participants of the study and Courtney Dickinson, Stephanie Friedhoff, Michael Hirsch, Sarah Zuckermann, and Joshua Schuler for assistance with study recruitment. We thank Tucker Lynn for experimental assistance and the BioMicroCenter at MIT for assistance with DNA sequencing. We thank Otto Cordero, Greg Fournier, and members of the Lieberman Lab for feedback on the manuscript.

This work was funded by a pilot grant from the MIT Center for Microbiome Informatics and Therapeutics, by a Smith Family Foundation Award for Excellence in Biomedical Research, and NIH grant 1DP2GM140922 (all to TDL).

## Author contributions

Conceptualization: JSB, TDL; Methodology: JSB, EQ, CPM, ADT, AC, TDL; Investigation: JSB, EQ, CPM, ADT, AC, TDL; Funding acquisition: TDL; Writing – original draft: JSB, TDL; Writing – review & editing: JSB, EQ, CPM, ADT, AC, TDL.

## Competing interests

Authors declare that they have no competing interests.

## Data and materials availability

Sequencing data is available on the NCBI Sequence Read Archive under Bioproject #PRJNA1052084. All code needed to reproduce the analysis in this work can be found at https://github.com/jsbaker760/highres_dynamics.

## Materials and Methods

### Sample collection

We opportunistically sampled subjects at a single K-8 School at five events over the course of 1.4 years. Consent was obtained from each individual under MIT IRB 1802230066. For children, both parental consent and child assent were obtained. In total, this work includes samples from 57 individuals belonging to 23 nuclear families, in addition to two researchers who were collecting samples. Before sampling, subjects filled out a questionnaire with basic health and demographic information, including the time since their most recent antibiotic use. This questionnaire was used primarily to identify potential outliers, and details are withheld to avoid deidentification, with the exception of time since antibiotic usage (Supplementary Table 1). Only one subject was excluded from analyses based on metadata; subject 8PB indicated they were currently taking topical antibiotics at timepoint 1 and we therefore included isolates from this subject in transmission analyses but excluded this sample from all metagenomic analyses.

Each time a subject was sampled, swabs were collected from four locations (Forehead, Nose, Cheek, and Chin) on the left-hand side of the face. Each set of four sampling sites is referred to as a “kit” in the methods. To collect skin-dwelling bacteria while reducing the collection of incidental surface-dwelling bacteria, debris, and cosmetics, we first firmly wiped all four skin sites with a cleansing wipe (Cetaphil, DPCI: 037-12-0886), ensuring that a different side of the wipe was used for each site. We then used saline cotton wipes (Winner #504021), using the same technique, to remove any residue from the cleansing wipe. After a 1-minute drying period, samples were taken from the forehead, nose, cheek and chin, in that order. During the drying period, examiners recorded whether the subject had facial hair and noted the presence of acne lesions and categorized the subject’s skin at the time of sampling as dry, oily, or combination. For each sample, a swab (Puritan, REF#25-1506 1PF 100) was first dipped into sterile PBS with .05% w/v Tween80, and then applied with light pressure with upward and downward strokes while moving swabs laterally and rotating the swab. The tip of each swab was immediately broken off into 1.5mL cryotubes containing 1000µL of storage solution (20:80 glycerol:PBS with 0.05% w/v Cysteine (as a reducing agent) which had been vacuum filtered at 0.22um and reduced in an anaerobic chamber). Upon completion of a sampling kit, all samples were vortexed for 60s at maximum speed and placed on dry ice. Samples were transferred to long-term storage in a -80°C freezer the same day they were collected.

### Isolating colonies

We isolated colonies on both LB agar plates cultured in anaerobic conditions (to enrich for *Cutibacterium* species) and TSB agar plates cultured in aerobic media (to enrich for *Staphylococcus* species).

For each sampling kit cultured, tubes were thawed and resuspended by vortexing for 60s at max speed. A serial dilution was prepared using 100µL of this solution and plated onto petri dishes with disposable spreaders. After growth for 3 days aerobically (TSB plates) or up to 7 days anaerobically (LB plates), colonies originating from each tube were selected from the most dilute plate or plates possible to limit competition. When few isolates could be obtained for the initial round of culturing, particularly common in younger subjects with a lower abundance of bacteria, additional serial dilutions were made using lower dilutions.

Each colony was picked into a 1000µL of liquid medium (either LB or TSB) and grown for 3 days aerobically or up to 7 days anaerobically at 37°C. Isolates were not frozen prior to outgrowth. Colonies were picked either using a disposable loop or a PIXL colony picking robot (Singer Instruments, PIX-001). When picking by hand, we randomly selected colonies which were far enough apart to avoid cross-contamination. When using the PIXL to pick colonies, we implemented size, circularity, and radius filters to randomly pick colonies that were sufficiently far apart while limiting cross-contamination. These cultures were used for both DNA extraction and storage. For each round of culturing, we used two types of negative controls to ensure the absence of contaminating bacteria. First, an entire unused sampling tube from the sampling event poured onto a single petri dish to control for batch contamination. Second, a plate with an aliquot of the PBS used during serial dilutions, after each sample had been plated, to control for reagent contamination during plating. The first type of negative controls never showed growth indicative of bacterial contamination in the collection tubes. When contaminants were identified in the second type of negative controls due to reagent contamination, all plates were discarded and the experiment was repeated.

No restreaking of isolates was performed in order to limit within-lab evolution. Isolates and downstream steps were processed in 96-well format and thus had some potential for cross-contamination. Methods for removing isolates with between-species contamination can be found in <u>Filtering of isolates and calculating pairwise distances</u>. Methods for removing isolates with suspected within-species contamination can be found in <u>Clustering isolates into lineages.</u>

### DNA preparation for isolate genomes

Cells were spun down at 4200g for 10 minutes and the supernatant was removed. Pellets were resuspended in 100µL of PBS and lysozyme (Sigma #L6876) was added to reach a final concentration of 1000U/mL. Plates were sealed and vortexed until the suspension was homogenous (at least 2 minutes), then incubated at 37°C overnight. To each well, 2µL of 20ug/µL proteinase K (NEB, #P8107S) and 2µL of 0.8% SDS solution were added, and plates were then incubated at 55°C for an additional 3 hours, after which the cell suspensions had no apparent turbidity. DNA was purified from these cell lysates using the standard protocol in the DNA extraction kits (Zymo Research #D3010), except that DNA was resuspended in sterile water in the final step. DNA was quantified using SYBR Safe (Thermo, #S33102) and normalized the concentration of each sample to between 1 and 5 ng/µL.To prepare libraries for Illumina sequencing, the tagmentation-based plexWell DNA library prep kits (Seqwell, #PW096) was used according to the manufacturer’s protocol.

### Metagenomic samples

Lysates were prepared as above, starting with 100µL aliquots of the primary samples. All reagents were sterilized, vacuum filtered, and autoclaved prior to use. DNA was purified using Purelink gDNA extraction kits (Thermo, #K182104A) as they improved yield in these low-biomass samples. The only modification to the Purelink protocol was elution of DNA into 10µL of sterile water three consecutive times, incubating the plate for 2 minutes at 37°C each time. DNA libraries were prepared using HackFlex^59^ with the following modifications: (1) the tagmentation was scaled to start with 10µL of DNA; (2) the tagmentation-stop step was skipped; (3) we used standard Illumina barcodes for plates MG1-3 and UDI primers (20091660) for plates MG4-7 (Supplemental Table 3); (4) we used KAPA HiFi master mix (Roche, 07958935001) and 19 cycles of PCR.

### DNA Sequencing and quality filtering

All samples were sequenced using the Illumina platform, on a variety of instruments and either 75bp or 150bp reads. Some libraries were re-sequenced to achieve higher depth. Isolates that met all below filters had a median depth of (8.6×10^5^ reads). Metagenomic samples that met all below filters had a median depth of 3.1×10^6^ reads. Details for each sample are found in Supplementary Tables 3 and 5. Raw reads were filtered and trimmed as previously described^16^.

### Species and genus-level microbiome composition from metagenomics

All reads from metagenomics samples were de-duplicated with the function dedupe.sh (-Xmx100g) from BBMAP (v39)^60^. Kraken2^61^ (--minimum-hits-group 3 –confidence .1) was used (v2.1.7, PlusPF db built 01/12/24) to classify reads and Bracken (v2.9)^62^ was used to estimate abundances at the species and genus levels. Immediately after classification, reads labeled as *Homo sapiens* were discarded. Reads labeled as *Aeromonas sp.* were also removed because they were suspected to emerge from contamination due to their high abundance in negative control samples. Abundances of taxa beginning with “Human” or “*Toxoplasma*” and all taxa which were found at <0.5% across all samples were removed from every sample and abundances were re-normalized. Samples with <100,000 reads assigned at the species-level from Bracken, across sequencing runs, were removed from downstream analysis.

Metagenomic abundances from tubes in the same kit were considered together since we found minimal compositional differences (Figure S1). Bracken abundances were averaged and re-normalized, resulting in 101 kit-level samples, each reflecting a single time point. Species and genus-level microbiome data are available in Supplementary Table 3.

The pairwise species-level community distances between samples were calculated using Euclidean distance. Hierarchical clustering revealed two clusters (Figure 2, Figure S2). The exact same set of clusters were found when using Ward linkage or complete linkage, and multiple evaluation methods supported clustering the data into two clusters (Figure S2). Cluster FC2 contains samples from older children (median 14.1 years old at time of sampling) and all parents except 11PA (Fig 2C, S3A). Cluster FC1 contained samples from younger children (median 11.4 years old at time of sampling) and parent 11PA.

Child subjects were categorized as an “FC2 Child” if any of their time point samples were clustered into FC2. Subjects were only categorized as an “FC1 child” if all samples from that subject were FC1 (Fig. S2C). Subject 11PA was included as a parent for downstream analyses despite their aberrant composition. To obtain per-subject average abundances, Bracken abundances from kits were averaged and re-normalized as above.

### Filtering of isolates and calculating pairwise distances

Given that isolates were grown in permissive culturing conditions, not checked for species identity, or re-streaked before sequencing, we first determined which of the 6,421 sequenced isolates were *C. acnes*, *S. epidermidis*, other species, or might have significant levels of contamination from additional species by examining genome content. We reasoned that high-quality samples of target species would contain the core genes for that species, but not core genes for related species. Genomes were assembled for each isolate using Spades (v.3.15.3, --isolate -k 21,33,55,77,99,127 --phred-offset 33)^63^ and then annotated with prokka (v 1.14.6, --compliant --force --mincontiglen 500)^64^. We did not filter low-coverage contigs from these assemblies in order to be more sensitive to low-level contamination. CDHIT (v 4.8.1, -p .9)^65^ was then used to cluster the homologous protein-coding genes of these assemblies in addition to 79 publicly available reference genomes from NCBI (Supplementary Table 2). Core genes for each of these reference species were defined as the CDHIT gene clusters which were present in 100% of reference genomes for that species, but not found in any reference genomes for other species (1,042 gene clusters for *S. epidermidis* and 1,592 for *C. acnes*). These gene clusters were used only for initial filtering– see <u>Identifying genes within and across lineage assemblies</u> for methods used to identify homologous genes from lineage co-assemblies, which were used for all presented analyses in this work involving gene content. Samples with >87% of *C. acnes* core genes and <10% of the core genomes of *S. epidermidis* were included as *C. acnes*. For *S. epidermidis*, samples with >90% of the *S. epidermidis* core gene clusters and <1% of the core genes of *S. capitis* and *C. acnes* and at least 2× mean coverage from spades were included. This resulted in 2,300 *C. acnes* and 2,172 *S. epidermidis* isolates for further analysis.

We aligned reads to reference genomes to compute pairwise distances between samples and identify samples that might have two strains of the same species. First, the sequencing reads of each *C. acnes* isolate was aligned to the Pacnes_C1 reference genome, and the sequencing reads of each *S. epidermidis* sample was aligned to SepidermidisATCC12228 using bowtie2 (v2.2.6 --very-sensitive --threads 8 -X 2000 --no-mixed --dovetail -x). Samtools mpileup (v.1.5; -q30 -x -s -O -d3000) was used to identify candidate SNPs for each representative isolate. For every potential position in each sample, a nucleotide call was assigned to that position based on the major allele nucleotide if the major allele frequency was above 0.79, the FQ score produced by samtools was less than -45, and the coverage was above 3. Otherwise the position was marked as ambiguous “N”.We then discarded genomic positions across samples if the median coverage across isolates was less than 3, or 15% or fewer of isolates had a non-ambiguous allele at that position. Pairwise distance was calculated as the sum of positions where both samples had different, non-ambiguous, nucleotide calls. At this stage, 11 of the remaining 2300 *C. acnes* isolates were removed because >50% of positions were ambiguous, suggesting the presence of two distant lineages. Isolates with more closely related lineages in a single isolate were removed during clustering (see next section).

### Clustering isolates into lineages

To cluster genomes which passed the above filters into closely-related clades consistent with on-person evolution or recent transmission, we ran clustering algorithms across a wide range of parameters and chose the clustering instance with the most isolates in self-consistent clusters as the best clustering instance. Clusters are considered consistent if and only if all isolates in each cluster are more closely related to each other than to any isolate outside of their clusters^66^. The consistency criteria was inspired by the nature of our data set and previous observations^16,20^; given that samples were taken from unrelated two-generation families, clusters of related strains with intermediate distances are not expected. This consistency criteria also removed samples artificially close to two clusters because of contamination (see next paragraph). A parameter scanning process was required given the absence of solved methods for clustering generally^67^ and for this use case. Where multiple parameters gave an identical result, only one set of parameters was chosen and listed below. Initial parameter ranges were chosen based on the fact that the typical molecular clock range bacteria *in vivo* of about 1 mutation/core-genome/year ^8^ and that we expect isolates from a family to have a single-celled ancestor within the past 50-100 years. Complete lists of parameters attempted are available in the github repository.

Sequencing reads for some isolates showed evidence of cross-contamination by isolates from closely related strains that were not removed by the major allele frequency filter. In such mixed isolates, the number of calls masked by our major allele frequency filter can be relatively small relative to the size of the genome, but still significant enough to artificially decrease inter-isolate distances. The inclusion of such mixtures can thus agglomerate unrelated clusters. To remove these isolates and overcome this challenge for clustering, we: (1) used a network property called the local clustering coefficient (LCC)^68^, which represents how many of one’s neighbors are neighbors with one another to pre-filter samples; and (2) required self-consistency (as described above) for all of our clusters We scanned multiple parameters for LCC cutoffs for sample inclusion for each instance of clustering. In the final best clustering parameter sets, *C. acnes* isolates with an LCC above 0.79 at a 200 SNP cutoff and *S. epidermidis* isolates with a LCC above 0.98 at a 700 SNP *s* were included. This process removed 78 *C. acnes* and 69 *S. epidermidis* isolates.

Clustering was performed with a two step process: initial cluster generation and greedy addition. Initial cluster generation was performed over a range of parameters for both DBSCAN ^69^ and ANCHOR, a custom algorithm. ANCHOR was designed to produce self-consistent clusters while dealing with noisy distance metrics. ANCHOR is initialized by defining anchor points, each of which represents a cluster to which isolates will later be added if they are within a given distance (C_1_). Anchor points are first distributed such that they are at least 2C_1_+1 away from each other as follows: First, a seed isolate was chosen as the first anchor, which was kept the same for each clustering instance. The seed isolate was the isolate which was the farthest away from its nearest isolate. After the addition of each anchor, the isolate furthest away from all anchors is considered as a candidate anchor for addition. Candidate anchors were added if they were at least C_1_+1 away from every already chosen anchor. When no further isolates can be added, the initialization step completes. We then added any isolates which could be unambiguously assigned to these clusters (< C_1_ from only one cluster).

The effects of the aforementioned noise, and as well as potential processes of accelerated mutation like hypermutation, mean that isolates may be >C_1_ from an anchor even if they are of high quality. We therefore implemented a greedy addition process to identify and include these isolates for all instances of clustering (across algorithms, species, and parameters), that resulted in self consistent clusters with a high number of clustered isolates (at least 1800 for *C. acnes* and 1600 for *S. epidermidis*) and a number of lineages consistent with our early clustering attempts (96-150 for *C. acnes* and 100-200 for *S. epidermidis*). First, all unclustered isolates are sorted in ascending order by their distance to the nearest cluster. Then, for each of these, we added it to its nearest cluster only if clusters would remain self consistent and this isolate is not >75 SNPs (a threshold chosen based on early optimal values of C_1_) away from any isolate in its cluster. This process terminates when no isolate can be added without violating either of these properties

We then chose the self-consistent clustering instance which contained the highest number of clustered isolates in clusters of any size. For *S. epidermidis*, DBSCAN (eps=88, minpts=3) gave the optimal result, with 2025 isolates clustered into 78 lineages with at least three isolates, 19 two isolate lineages, and 40 singletons. For *C. acnes,* and ANCHOR (C1=39) gave the optimal result, with 2164 isolates clustered in 98 lineages with at least three isolates, 13 two-isolate lineages and 21 singletons. Clusters with fewer than three isolates were excluded from all downstream analyses as they could not confidently be distinguished from transient colonization and cannot yield informative phylogenies, though clusters with two isolates were used to build the PHLAME classifier. To confirm that we had not overclustered isolates and artificially broken up related isolates into distinct lineages, we confirmed that nearby clusters were not systematically from the same family (Figure S22).

### Detecting lineages and phylotypes from metagenomics

To identify lineages from metagenomics data, we first built a database for each species by aligning all 2164 *C. acnes* and 2025 *S. epidermidis* isolate genomes which passed quality filtering and clustering to the Pacnes_C1 and SepidermidisATCC12228 reference genomes, respectively, using bowtie2 (v.2.2.6; -X 2000 --no-mixed --dovetail). Duplicate reads were filtered out using samtools markdup (v.1.15; -d 100 --mode s). Samtools mpileup (v.1.5; -q30 -x -s -O -d3000) was used to identify candidate polymorphisms. PHLAME (v.0.1)^70^ was used subsequently to build a database of lineage-specific alleles (common to all isolates of a lineage, but not found in any other lineage) within core genome regions (found in >90% of isolates). Alleles were additionally included if <10% of the isolates in a lineage were missing an alignment position entirely, but the remaining isolates had a unanimous allele at that position.

For each sample, the number of reads covering each informative position (containing a lineage-specific SNP) both with and without the allele were extracted and used to determine the presence and frequency of lineages in each metagenomic sample. PHLAME avoids false positive detections of lineages that occur when only a subset of any set of lineage-specific alleles are covered by reads, which may instead represent the presence of an unknown but closely-related lineage in a sample. PHLAME models the number of reads covering a set of lineage-specific SNPs as coming from a zero-inflated negative binomial (ZINB) distribution with rate parameter λ, overdispersion parameter α, and zero-inflation parameter π. The π parameter represents the proportion of lineage-specific SNPs that are systematically missing from a sample. Gibbs sampling, implemented using JAGS v.4.3.0, was used to obtain posterior distributions over these 3 parameters by sampling their full conditional distributions according to the following hierarchical formulation of the ZINB distribution, where *x*_*i*_ represents the set of allele specific counts:

### Likelihoods

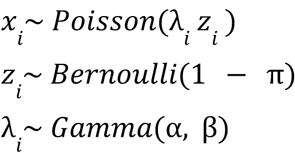

### Priors

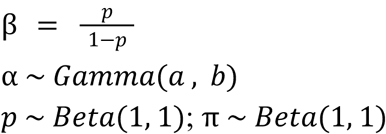

Parameters for a prior for the overdispersion parameter α were obtained empirically for each sample using the total number of reads at SNP-containing positions *y_i_*, where *s_yi_* ^2^ is the sample variance of *y_i_*.

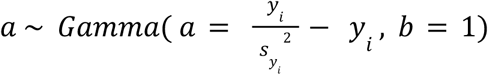

*i*This modeling was performed if the total number of reads supporting lineage-specific alleles in the sample was >10, otherwise the frequency of that lineage was set to 0. Each chain was run for 100,000 iterations with the first 10% discarded as burn-in. In order for a lineage to be detected, 50% of the posterior density over π was required to be below 0.35, and the lower bound of the 95% highest posterior density (HPD) interval of π was required to be <0.1. To increase sensitivity (and therefore limit false observations of gains and losses), if we additionally counted lineage detections where >50% of the posterior density of π was below <0.55 if that lineage was detected using the stricter parameters on the same subject at another time point.

The relative abundance of each lineage was calculated as the maximum likelihood estimate for λ for the allele-specific reads divided by the maximum likelihood estimate for λ for all reads at the same positions. Because each lineage in a sample is modeled independently without reference to others, the sum frequency of lineages in a sample can sum to <1 or rarely, >1. If the sum frequency of calls was >1, we re-normalized frequencies down to 1 (the max sum lineage frequency before this step was 1.05).

We also grouped lineages into major phylogenetic clades, deemed phylotypes and sub-phylotypes, to search for patterns in intraspecies diversity in subjects without isolates (which therefore are not represented in the lineage database) and because some metagenomic samples were not sequenced to a high enough depth to achieve complete lineage-level resolution. To ensure a wide collection of phylogenetic diversity for *C. acnes*, we also included four publicly available isolates from *C. acnes* SLST L (ribotype III) into the reference database at the phylotype level (NCBI strain names: Cacnes_PMH7, Cacnes_PMH5, Cacnes_JCM_18919, Cacnes_JCM_18909). Because the raw reads for these genomes were not publicly available, we first simulated reads from each genome using wgsim (v.0.3.1-r13; -e 0.0 -d 500 -N 500000 -1 150 -2 150 -r 0.0 -R 0.0 -X 0.0) before inputting into the PHLAME pipeline. *C. acnes* phylotypes were defined following an existing single locus strain-typing (SLST) scheme^71^. Four *S. epidermidis* phylotypes were determined manually by cutting at the longest branch lengths of the inter-lineage phylogeny (which matches other *S. epidermidis* phylogenies; Figure S20). We assigned at the sub-phylotype level for both species by picking the set of monophyletic clades that led to the highest average percent assigned across concatenated metagenomics samples. Phylotypes and sub-phylotypes were detected in a similar way to lineages, using instead a database of specific SNPs common to all isolates in a phylotype but not found in any other phylotypes.

### Filtering metagenomic samples

Lineages were defined as “potentially contaminating lineages” if they were found in any sample from any handler (researcher who collected or processed primary samples) by PHLAME at >3% abundance. All tubes in which the total abundance of potentially contaminating lineages was >5% were excluded from all analyses (13 tubes).

Three types of lineages were used to build the PHLAME classifier but were excluded from metagenomic-determined relative abundance after classification: (1) Lineages collected from two handlers (9 *C. acnes* lineages, 0 *S. epidermidis* lineages), which we used to identify and remove samples with potential traces of laboratory contamination (see <u>Filtering metagenomic</u> <u>samples above);</u> (2) lineages for which there were only 2 corresponding isolates; (3) two lineages which appeared to be contaminants from mock community samples processed on the same plates. *C. acnes* lineages 43 and 116 and *S. epidermidis* lineages 6 and 45 were used in high-biomass mock communities (isolated from 1AA, 4AA, and 5PA) on plates MG1, MG2, and MG3 (Sup. Figure S23) and identified in many samples. We therefore conservatively removed their abundances from all samples. In total, the abundance data of 87 *C. acnes* and 76 *S. epidermidis* lineages with sufficient corresponding isolate data and no evidence of contamination were used.

### Lineage diversity and sharing across subjects

The combined set of lineage calls from isolates and metagenomics was used to determine how many lineages are on a person (Figure 3B-C) and how many are shared with other subjects (Figure 4A). Only subject/time point samples for which >70% of the metagenomic abundance could be assigned were used. Only lineages with an inferred abundance of >=1% in were included as present to limit spurious calls from computational or experimental errors. All lineages found across time points are included– a lineage is ‘shared’ between two subjects even if it is found in a single time point for each.

### Within lineage SNPs

Reference genomes were assembled for each lineage to identify mutations in genes present in the flexible gene content of lineages but absent in the reference genomes Pacnes_C1 or SepidermidisATCC12228. First, the same number of reads from each sample (equal to the number of reads in the sample with the least reads) were concatenated into individual files for each lineage. Then, Spades was used (-m 500 -k 21,33,55,77 --phred-offset 33 --careful) to assemble these reads into contigs. We then manually applied a minimum-coverage filter for contigs in each assembly individually in order to reduce the number of small and spurious contigs, which was usually at half the coverage of the longest contig. These cutoff values are found in Supplementary Table 5.

In order to identify outgroups for each clade, the average pairwise distance between lineages was obtained using the same approach as used for clustering. For each lineage, the 5 nearest lineages were identified, and the highest coverage sample from each of these lineages was used as the outgroup. Reads from each sample (all isolates from the lineage and all 5 outgroup samples) were realigned to the lineage’s co-assembly for downstream SNP identification and phylogeny construction using bowtie, as above.

To identify *de novo* SNPs in each lineage and build accurate phylogenies, calls were first strictly filtered using the same general process as above to identify positions where most samples have high-quality calls. To initially filter low-quality calls in the alignments to lineage co-assemblies, we considered calls with a FQ score <30, major allele frequency < 0.75 or coverage <4 as “N”. Next, we excluded positions >40% of samples have an N in that position (Figure S23). To build phylogenies, nucleotide calls at accepted positions were less strictly filtered to retain information at these highly informative positions. Major-allele nucleotides were used as calls provided they had a major allele frequency of 70% and a coverage of at 5 reads (forward or reverse). In addition to excluding low-quality positions and calls, we also excluded recombinant positions, which do not accumulate according to a molecular clock. We excluded regions where SNPs occurred within a 500bp sliding window with a covariation >0.75. We constructed maximum-parsimony phylogenies for each lineage using DNApars.

To identify ancestral nucleotides at each position (for MRCA and directionality analyses, see below), we first considered positions where at least 2 outgroup samples had the same call and there were no other alleles. At all other positions, the reference genome call was used. Remaining positions with N in the reference genome were replaced by the modal call from the ingroup.

### Identifying genes within and across lineage assemblies

First, clade co-assemblies were annotated using Bakta (v1.6.1)^72^. To identify which of these genes were the same across lineage co-assemblies, Panaroo (v1.5.0, sensitive mode)^73^ was used to cluster protein sequences into homologous clusters. Genes from different lineage genomes were considered the same gene if they were in the same Panaroo cluster.

### Adaptation analysis

To search for significantly mutated genes, we first counted how many times each gene was mutated. We excluded mutations which appeared to be the result of recombination (see: Within-lineage SNPs, Supplementary Table 6). We then computed the probability of observing at least as many mutations as the number observed according to a Poisson distribution as done previously^16^. This calculation takes into account the differences in gene length, codon distribution, and mutational spectrum across lineages by taking the average of each of these factors across lineages with mutation in the gene under consideration. To measure the false-discovery rate, we used the Benjamini-Hochberg procedure where each gene was treated as a separate hypothesis. Calculations of dN/dS for each gene were also computed as previously described in^16^. Results are summarized in Figure S16, and SNPs in individual lineages and in homologous gene clusters across lineages are listed in Supplementary Table 6.

### Subject dMRCAs

To calculate dMRCA_SUBJECT_ for a given lineage (Figs. 5 and 6), we only considered instances where each subject had at least 5 isolates in that lineage. Each subject’s isolates were considered independently; as such, per-subject lineage dMRCAs (dMRCA_SUBJECT_) were calculated as the mean number of *de novo* mutations excluding mutations common to all subjects’ isolates.

### Directionality of transmission

Directionality was only determined for lineages from which: (1) at least 2 subjects had at least 3 isolates each in a lineage; (2) at least one subject had a dMRCA (dMRCA_SUBJECT_) which was less than the dMRCA of the clade (recipient); (3) and at least one subject’s dMRCA_SUBJECT_ was the same as the root of the lineage (source). In some cases, one subject was clearly a recipient but multiple individuals could have been the source. In other cases, multiple subjects were the recipient from a single source. Where we list each instance of inferrable directionality in supplementary material (Figure S9), only cases in which the recipients and sources are from a single age class are shown for simplicity.

### Number of genotype transmissions

Since it was often not possible to determine which of the subjects were the source of the lineage and which was the donor, we conservatively estimated the number of transmitted genotypes with at least 5 isolates from each of at least 2 subjects using a parsimony assumption as previously reported^74^. For each lineage, the smallest number of genotypes required to explain the diversity on all subjects was calculated for each potential source. The instance (source) with the smallest total number of genotype transmission was chosen for each lineage. The number of inferred transmitted genotypes for each lineage are listed with associated metadata in Supplementary Table 4.

### Molecular clock estimation

Molecular clock signals from individual people were calculated for all instances where a subject had at least 5 isolates in a given lineage from at least 2 time points. For each of *C. acnes* lineage, there was no significant correlation of the number of *de novo* mutations in isolates versus time since initial sampling at a Benjamini-Hochberg false discovery rate (FDR) of 1% for *C. acnes*; at an FDR of 5%, the only clock rate was negative. For *S. epidermidis,* 3 lineages had significant clock signals for *S. epidermidis* after FDR correction at an FDR of 5%. The estimated rates for *S. epidermidis* ranged from 4.8 to 13 SNPs/genome/year. Since these estimates varied by a factor of ∼3, we also estimated a molecular clock for *S. epidermidis* across lineages and subjects. For each lineage and for each subject with at least 5 isolates at two time points, we plotted the average root-to-tip distance of isolates from each time point against the time elapsed since the first sampled time point. Together, a significant molecular clock signal of 4.37 mutations/genome/year was estimated, which was slightly slower and therefore more conservative than the slowest of the three individually calculated clock rates (meaning that turnover may be faster than reported here) and is consistent with published rates for this species^17,33–35^. Using the same approach for *C. acnes*, we were unable to obtain a significant molecular clock rate (Figure S14).

## Supplementary figures

**Figure S1.**
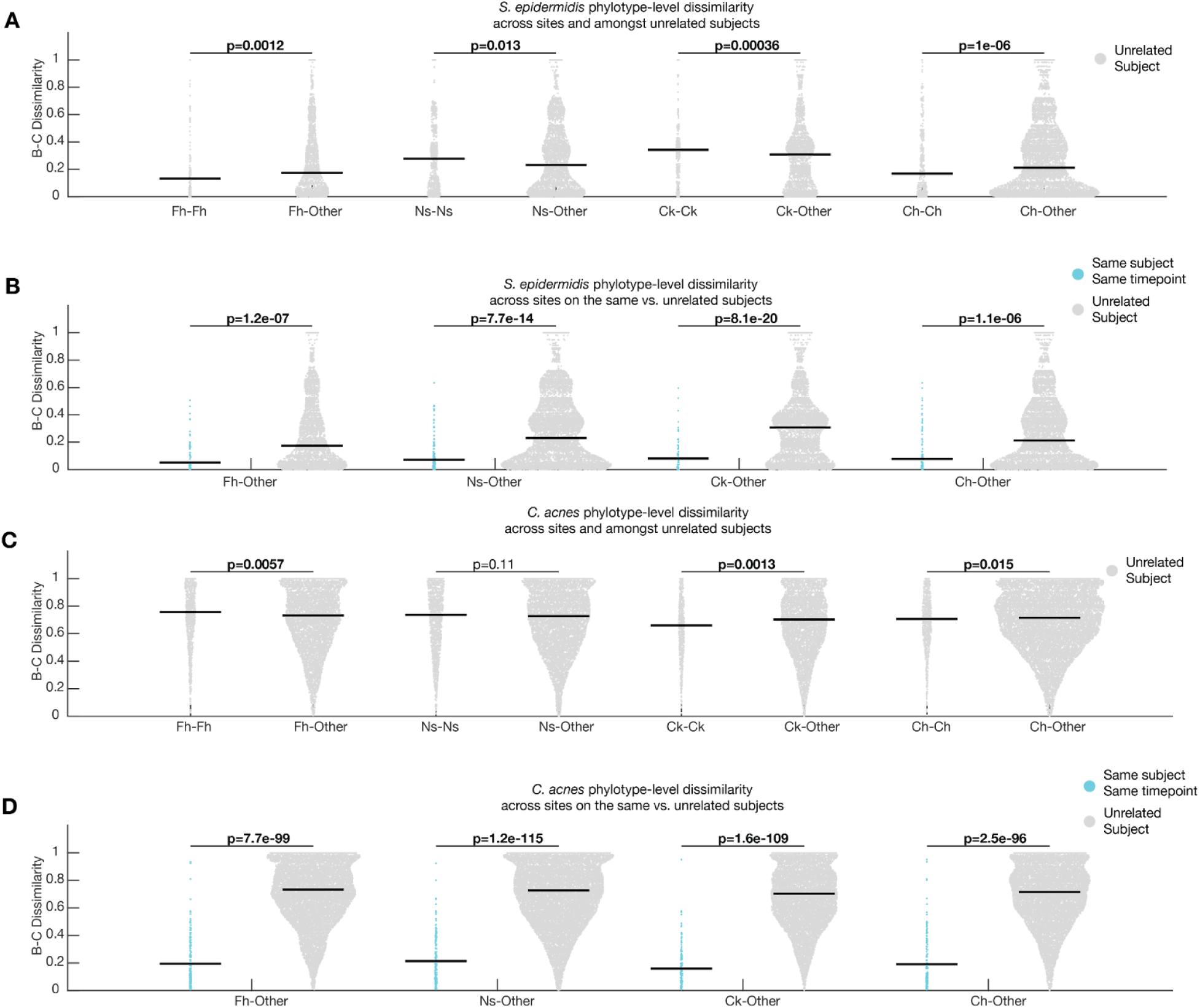
Compositional differences across facial sites at the phylotype level. **(A)** Bray-Curtis dissimilarity of *S. epidermidis* phylotype communities between the same site (ex. Fh-Fh) or different sites (ex. Fh-other) amongst unrelated people using metagenomic-derived phylotype abundances. Results show small but statistically significant site-specificity, consistent with^30^. **(B)** Bray-Curtis dissimilarity of *S. epidermidis* phylotype communities across dissimilar sites in the same subject at the same time point and across unrelated subjects, showing a much larger signal for person-specificity. **(C)** Bray-Curtis dissimilarity of *C. acnes* phylotype communities between the same site (ex. Fh-Fh) or different sites (ex. Fh-other) amongst unrelated people using metagenomic-derived phylotype abundances, showing small but statistically significant site-specificity. **(D)** Bray-Curtis dissimilarity of *C. acnes* phylotype communities across dissimilar sites in the same subject at the same time point and across unrelated subjects, showing strong signals of person-specificity. P-values in all panels come from two-sided Kolmogorov-Smirnov tests. Bolded P-values are significant at a false discovery rate of 5% using the Benjamin-Hochberg procedure

**Figure S2.**
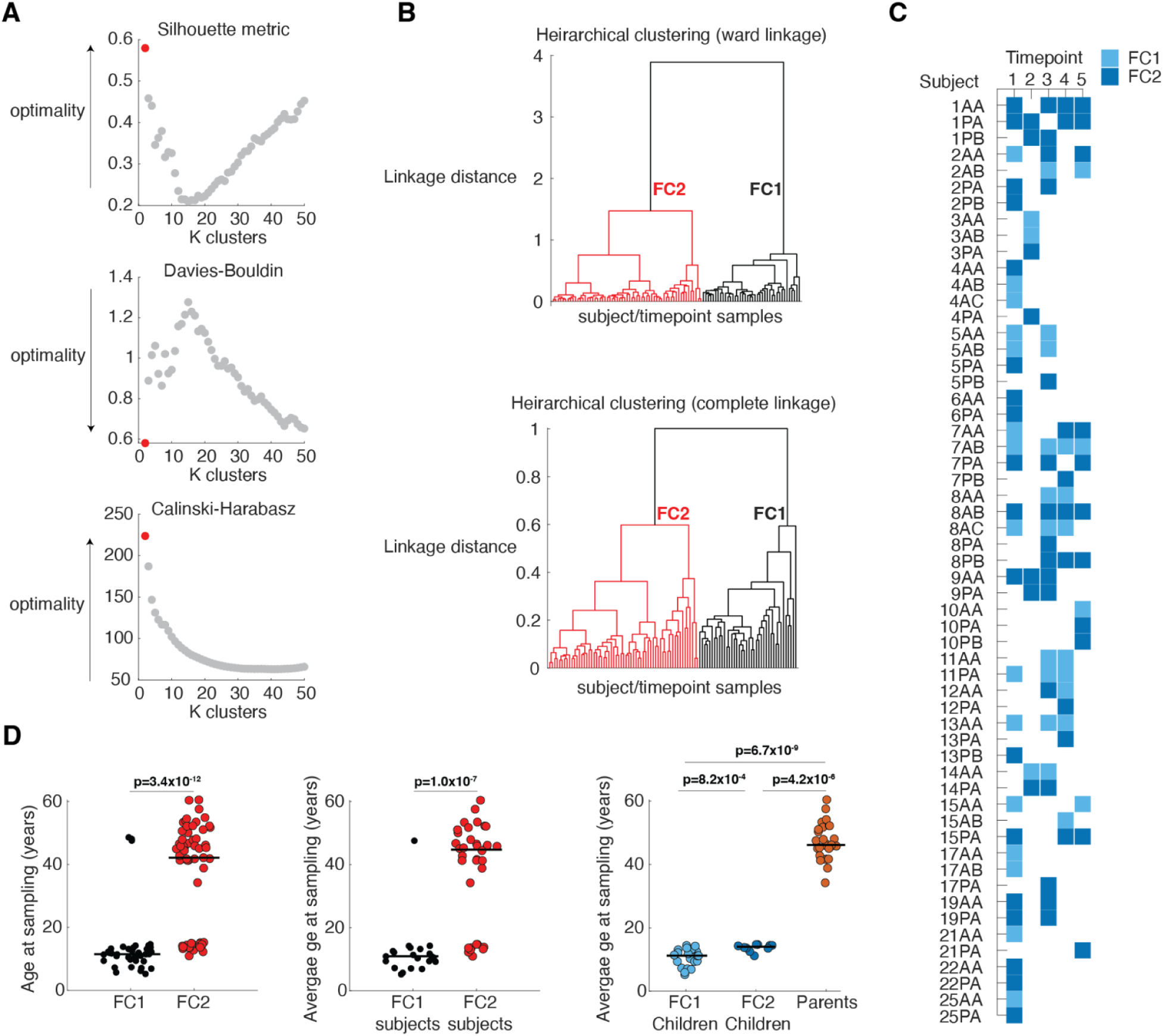
Facial Cutotype clustering and subject classification. **(A)** Three different optimality criteria (Silhouette, Davies-Bouldin, and Calinski-Harabasz) unanimously show that the data are best separated into k=2 clusters. **(B)** Linkage dendrograms show that cluster membership is the same whether using Ward or complete linkage, demonstrating robustness. **(C)** The Facial Cutotype (FC) of each subject/time point sample. The assignment of each sample is either FC1 or FC2. A single sample’s assignment to FC2 (dark blue) triggered classification as an FC2 subject. About half of the subjects were only sampled once. **(D)** Age differences between samples classified as FC1 or FC2 (left, each dot represents one subject at one timepoint), subjects classified as FC1 or FC2 (middle), and between FC1 Children, FC2 Children, and Parents (right). P-values are from two-tailed ranksum tests. Bolded P-values are significant at a false discovery rate of 5% using the Benjamin-Hochberg procedure.

**Figure S3.**
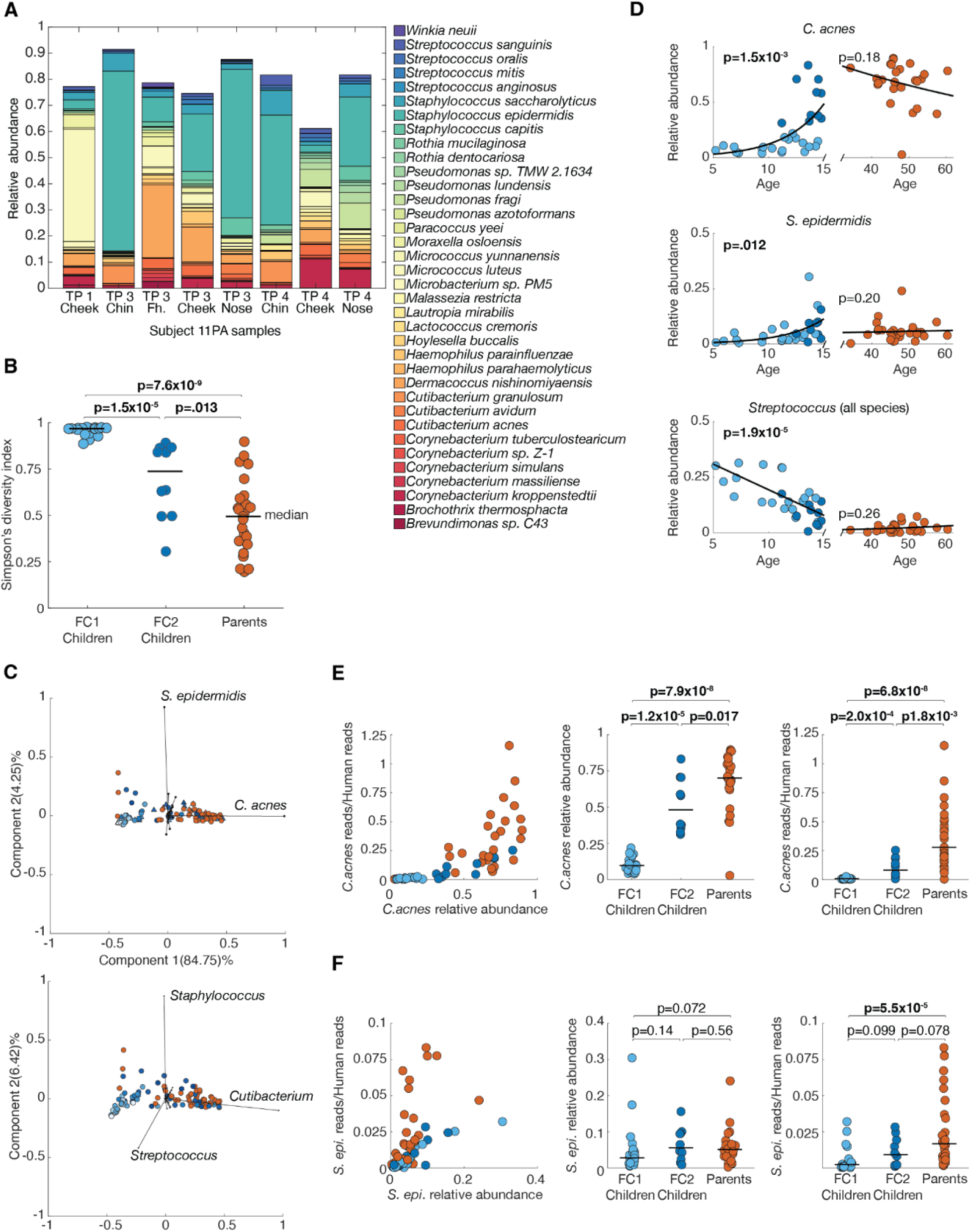
Additional details related to Facial Cutotype clustering. **(A)** Species-level abundance for all taxa found at >1% in subject 11PA, the only adult subject with any FC1-classified samples, showing that they have a consistently unusual microbiome composition across three time points, dominated by *Staphylococcus* and *Micrococcus*. **(B)** Species-level diversity is lost when children transition from FC1 to FC2, and similarly within FC2 between children and parents. **(C)** Principal component analysis on species level data and biplot confirm that 2 species are driving the majority of difference (See Figure 2). **(D)** The relative abundances of *C. acnes*, *S. epidermidis*, and *Streptococcus* are each significantly dynamic in children and do not change significantly in parents. For *C. acnes* **(E)** and *S. epidermidis* **(F)**: the ratio of species reads to human reads from Bracken (left), the relative abundance from the species-level Bracken data across three age groups (middle) and the ratio of species reads to human reads across three age groups (right) colored by subject assignment as FC1 Children, FC2 Children, or Parents. Lines indicate medians. These results indicate that the absolute abundance of both species is higher in parents despite minimal changes in relative abundance for *S. epidermidis.* Trendlines are first-degree exponentials for *C. acnes* and *S. epidermidis* in children, and linear elsewhere. P-values for all trendlines are for linear correlation (Pearson). P-values for comparisons between groups are from two-tailed ranksum tests. Bolded P-values are significant at a false discovery rate of 5% using the Benjamin-Hochberg procedure.

**Figure S4.**
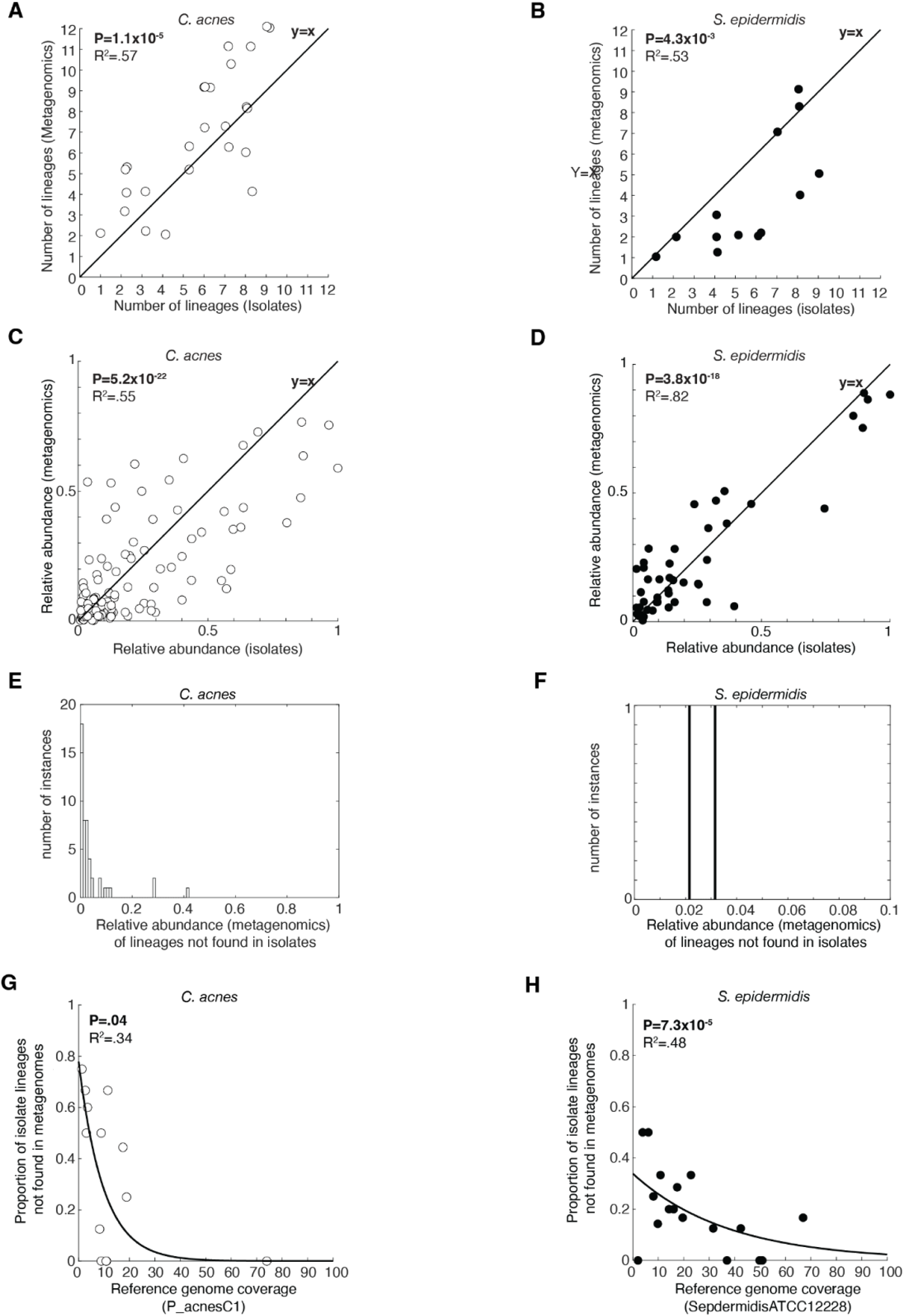
Comparison of lineages detection in metagenomics and isolates. **(A-B)** For samples with >25 isolates and >70% metagenomics assignment, the number of lineages found on subjects calculated from metagenomics or isolates alone is closely correlated. **(C-D)** in these samples, the abundances of lineages which are detected in both isolates and metagenomics show good agreement for both *C. acnes* and *S. epidermidis* **(E-F)** When lineages are found in metagenomics but not isolates, they tend to be low abundance for both *C. acnes* and *S. epidermidis*, **(G-H)** For samples with >25 isolates and >70% metagenomics assignment, the proportion of lineages found in isolates but not in metagenomics decreases with reference genome coverage. Trendlines for (G-H) are first-degree exponentials, and P-values and adjusted R-squared values are from Spearman correlation coefficients. For (A-D), P-values and R-squared values are from Pearson (linear) correlation coefficients.

**Figure S5.**
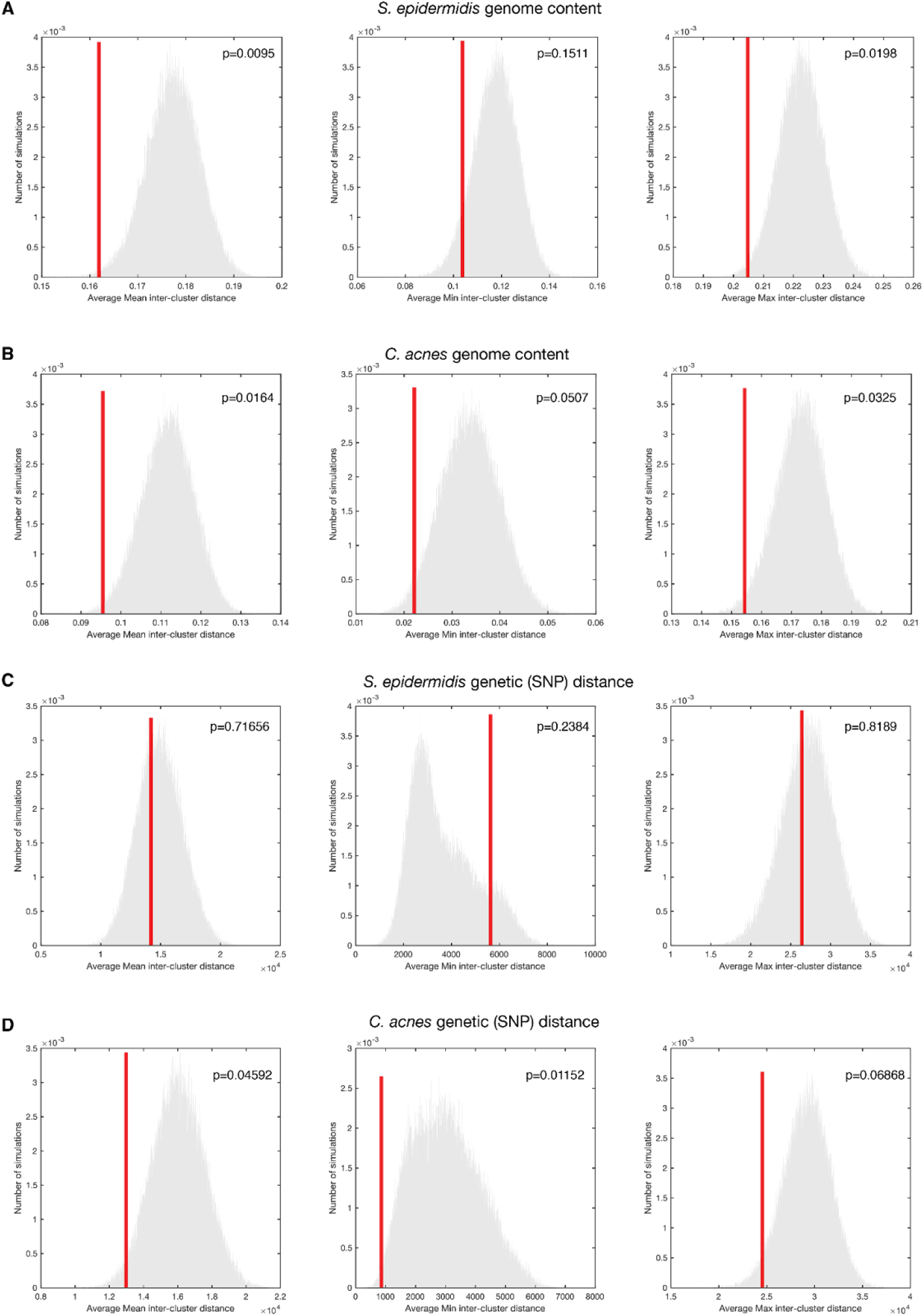
Complete set of simulations comparing the differences amongst coexisting lineages. In each box, gray histograms summarize the results of 10^5^ simulations of shuffling lineage identity across subjects (Methods). For each subject, the minimum (left), average (middle), and maximum (right) inter-cluster gene-content was calculated; The red bar shows the mean observed value across subjects in this study. Rows show gene-content differences between co-colonizing lineages for *S. epidermidis* (**A**) and *C. acnes* (**B)**, and average nucleotide distances for *S. epidermidis* (**C**) and *C. acnes* (**D)**. After Benjamini-Hochberg correction with a false discovery rate of 5%, none remain significant. These results suggest neutrality or modest underdispersion (niche filtering), suggestive of person-specificity, and do not support niche partitioning.

**Figure S6.**
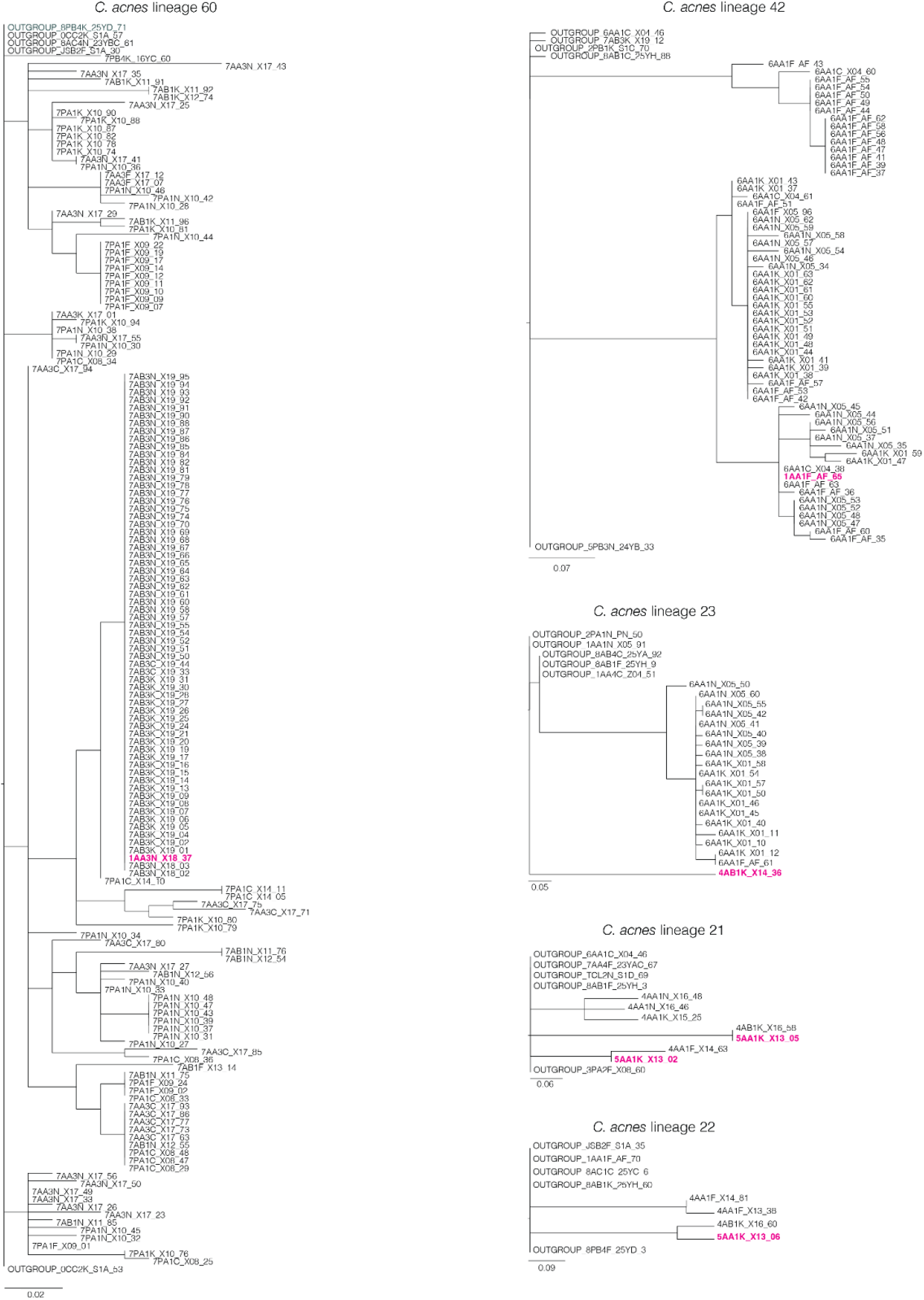
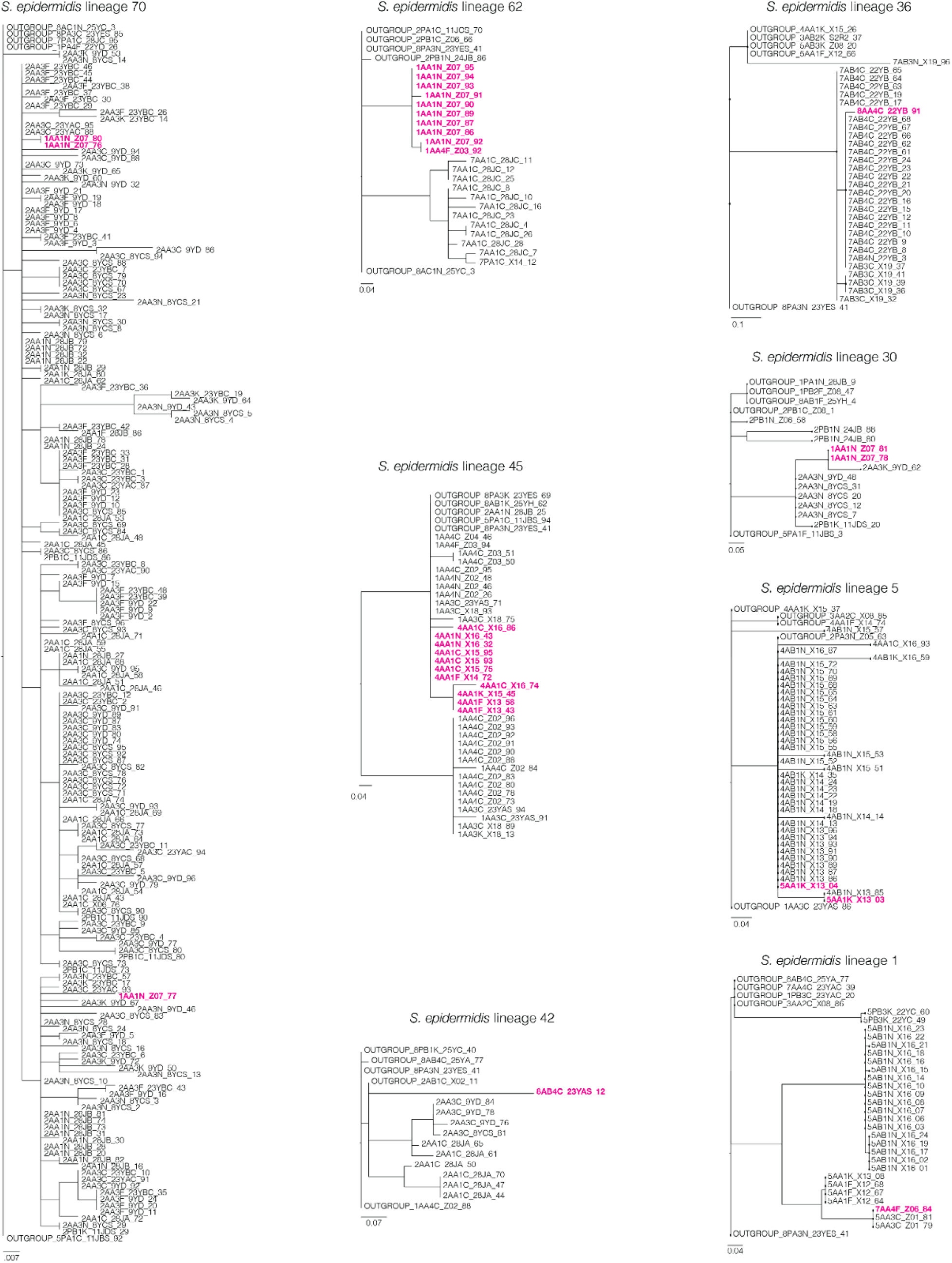
Phylogenies of each lineage with isolates collected from subjects in multiple families. In each of the 5 displayed lineages for *C. acnes* (8 for *S. epidermidis*) we identified small numbers of isolates from subjects outside of the majority family, which are shown in magenta. In each of these instances, the subject from outside the family was a student. Additionally, the phylogenetic topologies of all instances indicate either transmission from another student from the same school or transmission to two or more student subjects from a shared unknown source. While these observations could plausibly indicate the transient sharing of lineages due to exposure, most of these observations involve a small number of isolates and we were not able to corroborate most of them with metagenomics (Figure 4A) due to insufficient metagenomics assignment in the relevant samples (Methods) and could indicate experimental error.

**Figure S7.**
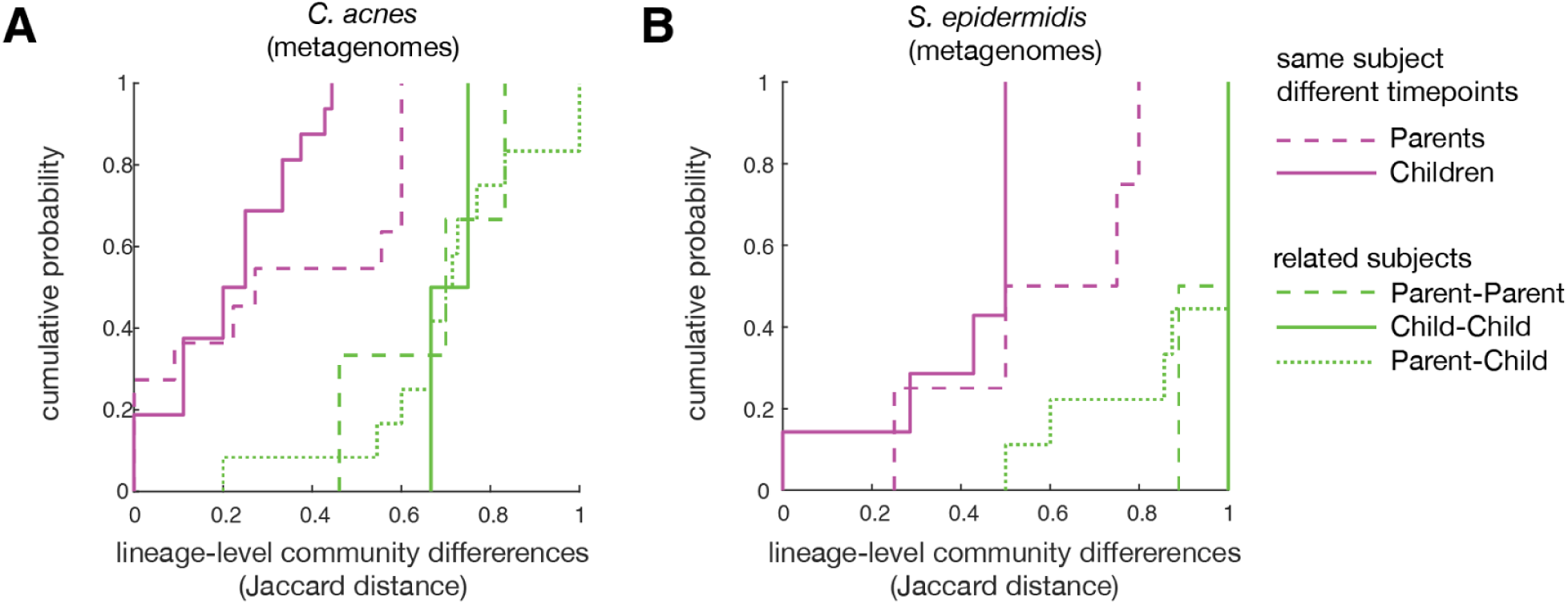
Lineage-level community similarity between related subjects. For *C. acnes* **(A)** and *S. epidermidis* **(B)**, the lineage-level community differences are shown for both the same subject over time (magenta) and between related subjects in different categories (green).

**Figure S8.**
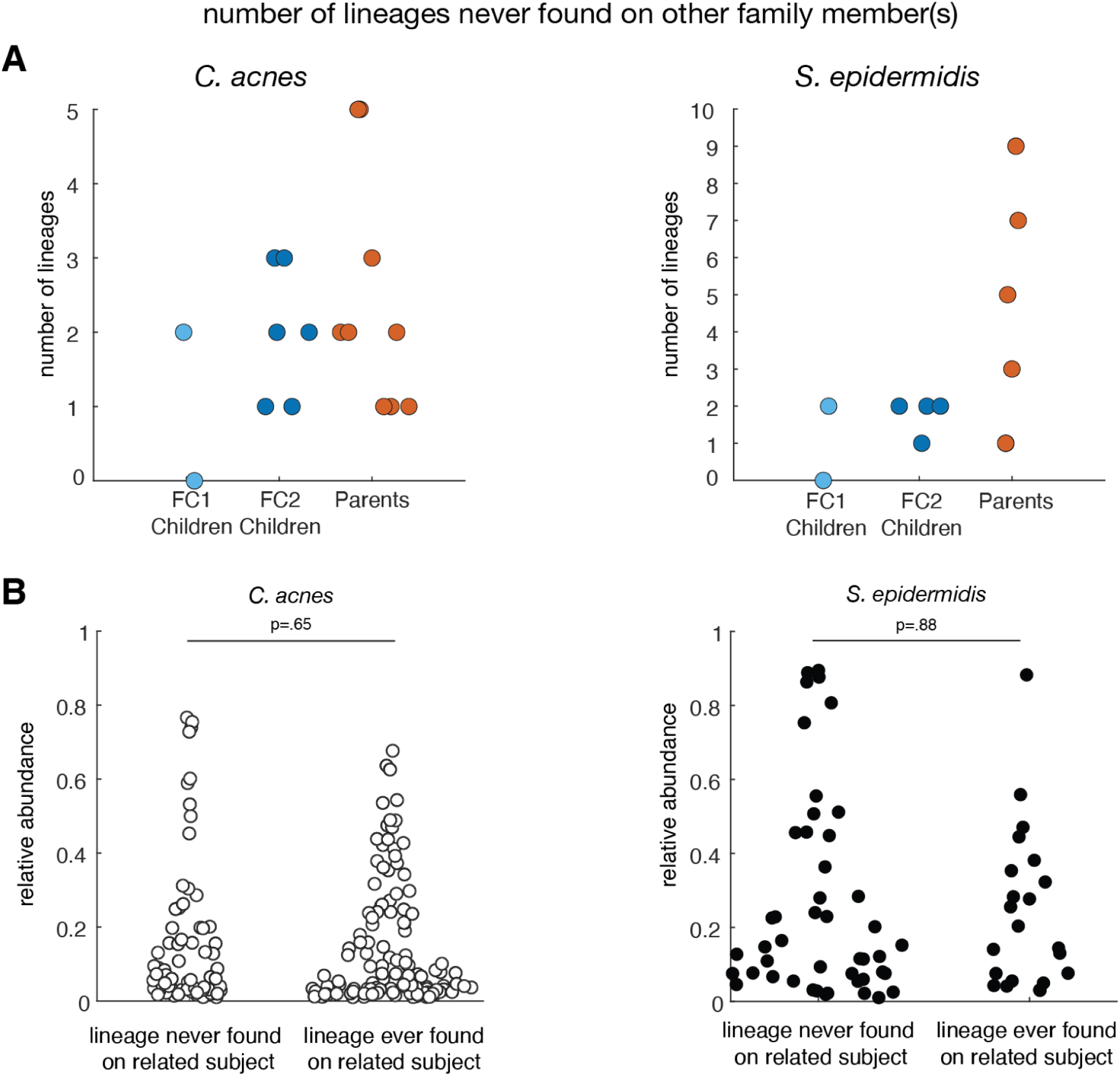
Lineages on individuals which are never found on their family members. **(A)** In both species, we observe lineages from metagenomics on individuals which are not found on their family members, indicating that lineage-level composition does not homogenize despite frequent contact. **(B)** From metagenomics, we do not find significant differences in the relative abundance between shared and unshared lineages. P-values are from two-tailed ranksum tests.

**Figure S9.**
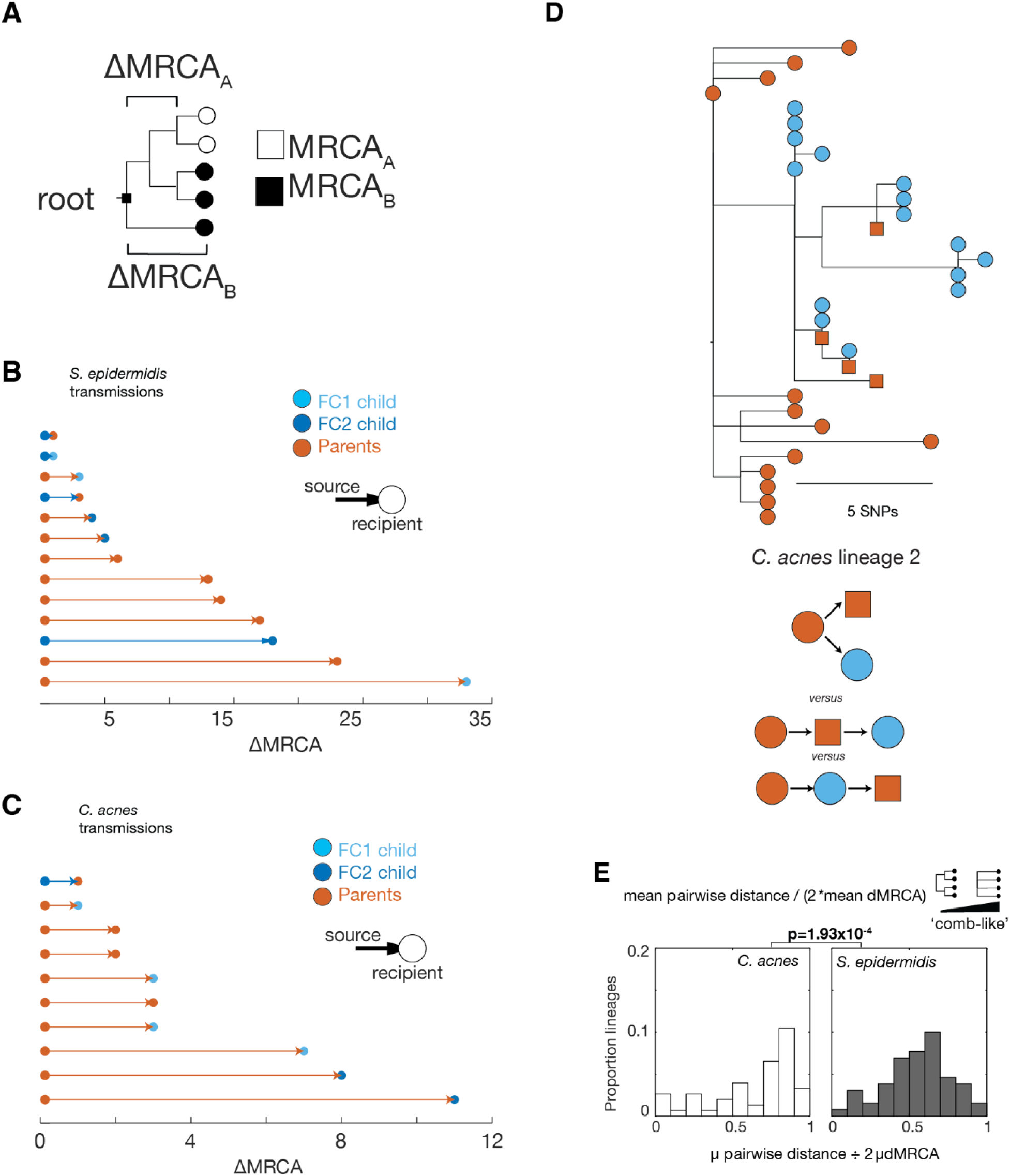
Instances of inferable transmission for both species. **(A)** Schematic representation of how ΔdMRCA, a metric used in inference of transmission, is calculated. Each open circle represents an isolate genome from one individual and each closed circle represents an isolate from another individual. **(B** and **C)** Each instance of *S. epidermidis* and *C. acnes* transmission with an inferable direction (Methods), colored by the subject category of the source (left dot and arrow) and recipient (right hand dot). **(D)** Example phylogeny of a shared lineage of unclear direction demonstrating the complexity of interpreting transmission dynamics, even when one subject (orange circles) is the donor: the source could have transmitted this lineage to both recipients independently (upper); to the subject represented by orange squares who subsequently transmitted it to the subject represented by blue dots (middle); or to the subject represented by blue dots who subsequently transmitted it to the subject represented by orange squares (lower). **(E)** To quantify the comb-like nature of lineage phylogenies, we calculated the ratio of average pairwise distance to average dMCRA for each lineage. This metric is similar to Tajima’s D, but is polarized based on rooting of lineages (to outgroups, Methods). Comb scores are significantly different between phylogenies of *C. acnes* and *S. epidermidis* lineages, with a less comb-like phylogenetic topology for *S. epidermidis*.

**Figure S10.**
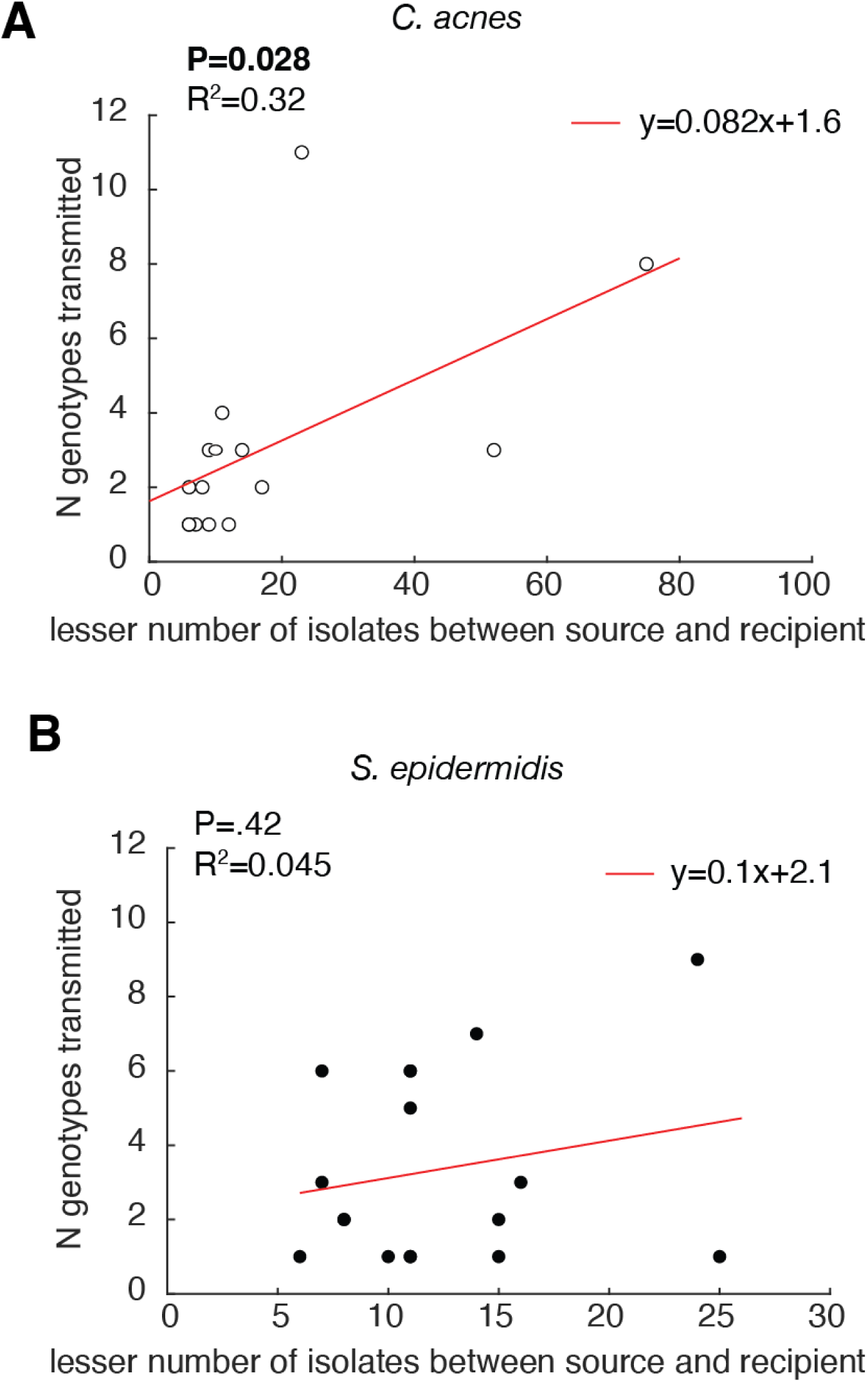
Comparison between the number of inferred transmitted genotypes and the number of isolates collected. **(A)** The number of genotypes transmitted to a subject is correlated with the lesser of the number of isolates from the donor and from the recipient, indicating that collecting more isolates would increase the number of inferred transmission events and that we are under-estimating the number of transmitted genotypes. **(B)** In *S. epidermidis,* the number of genotypes transmitted from a subject is not correlated with the lesser of the number of isolates from the donor and from the recipient, suggesting that we are not under-estimating the number of transmitted genotypes.

**Figure S11.**
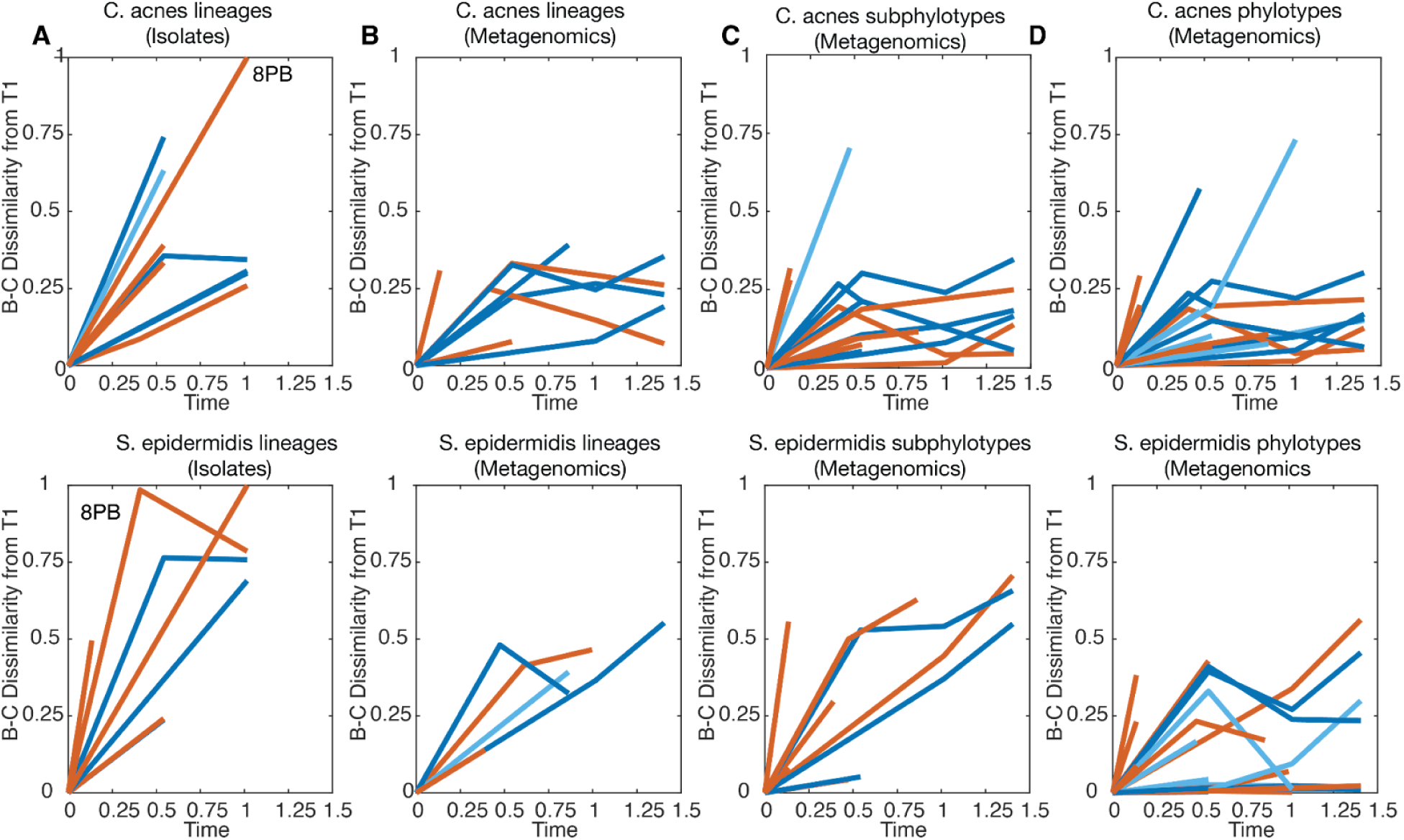
Bray-Curtis dissimilarity over time data for all subjects at different taxonomic levels. For each panel, the data for *C. acnes* are on the upper set of axes and the data for *S. epidermidis* are on the lower set of axes. **(A)** Lineage-level Bray-Curtis dissimilarity over time at the lineage level, with subject 8PB, whose data was excluded from metagenomics due to antibiotics use (methods) highlighted **(B)** as in A but for metagenomics **(C)** As in previous panels, but for the sub-phylotype level, which is only defined for *S. epidermidis* and not *C. acnes.* **(D)** As in previous panels, but for the phylotype-level from metagenomics. In all panels, Time refers to years since initial sampling. Isolate-inferred dynamics are likely to be overestimated due to the noise inherent in sampling low numbers of colonies.

**S12:**
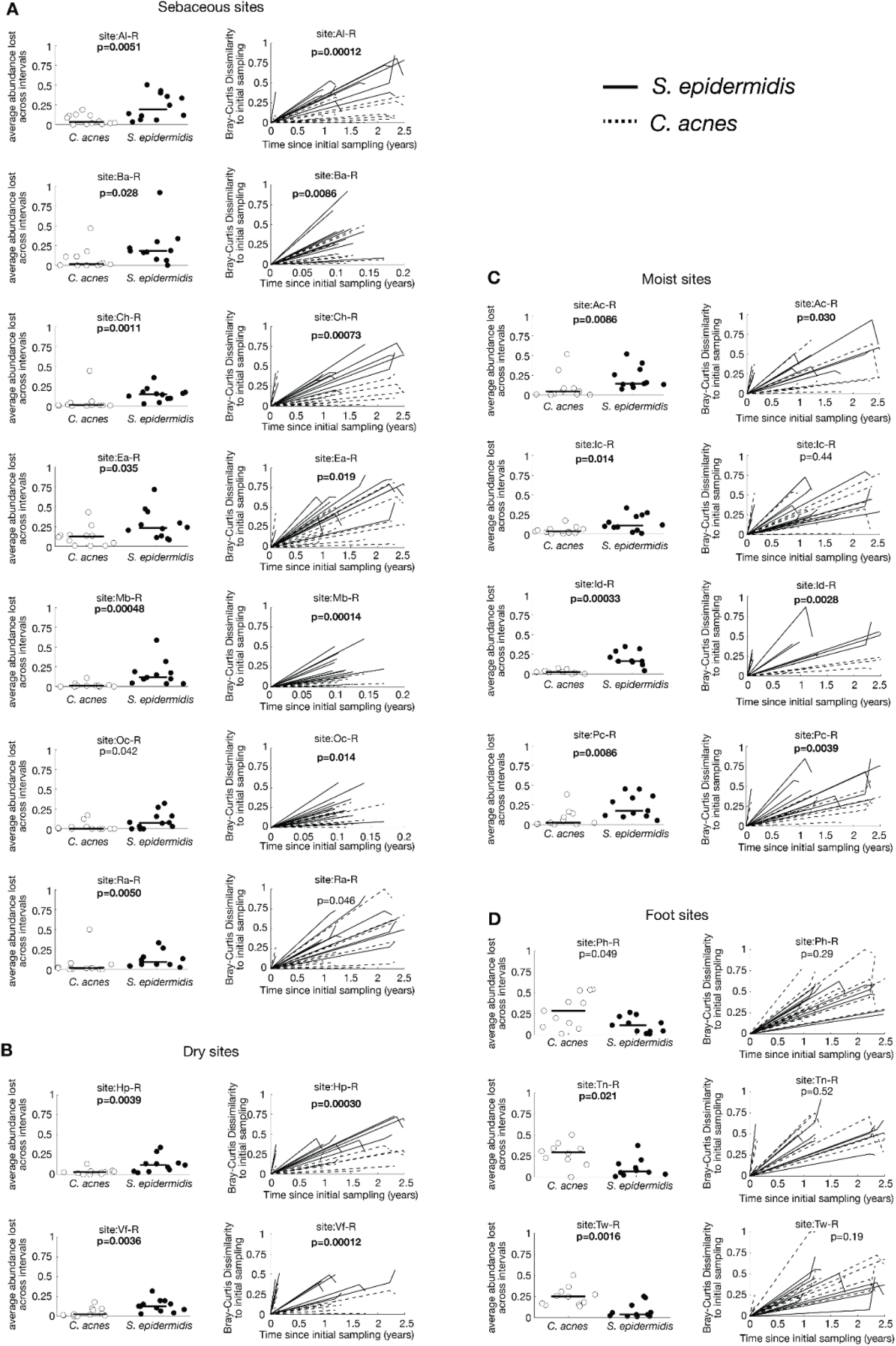
Strain-level turnover of *C. acnes* and *S. epidermidis* across body sites from Oh *et. al.* We measured the average strain-level abundance lost between sampling intervals per subject (as in Figure 5A-B) and the Bray-Curtis dissimilarity from the initial sampling over time (As in Figure 5C) from supplemental data provided by Oh *et al.*^5^. In almost all (A) sebaceous (B) dry and (C) moist skin sites, we found that the abundance of *S. epidermidis* strains lost between timepoints (left) and changes in strain-level community composition over time (right) are significantly higher than *C. acnes*, consistent with our results in Figure 5A-C. In foot sites (D), community-level changes are not significantly different between the species, and turnover of *C. acnes* appears to be higher than *S. epidermidis* in two of the three sites. All P-values are from two-tailed ranksum tests. P-values for Bray-Curtis dissimilarity compare the highest observed value per subject between the two species. Bolded P-values are significant at a false discovery rate of 5% using the Benjamin-Hochberg procedure.(Al-R: right alar crease, Ba-R: right back, Ch-R: right cheek, Ea-R: right external auditory canal, Mb-R: right manubrium, Oc-R: occiput, Ra-R: right retroauricular crease, Hp-R: right hypothenar palm, Vf-R: right volar forearm, Ac-R: right antecubital fossa, Ic-R: right inguinal crease, Id-R: right interdigital web, Pc-R: right popliteal fossa, Ph-R: right plantar heel, Tn-R: right toe nail, Tw-R: right toe web)

**Figure S13.**
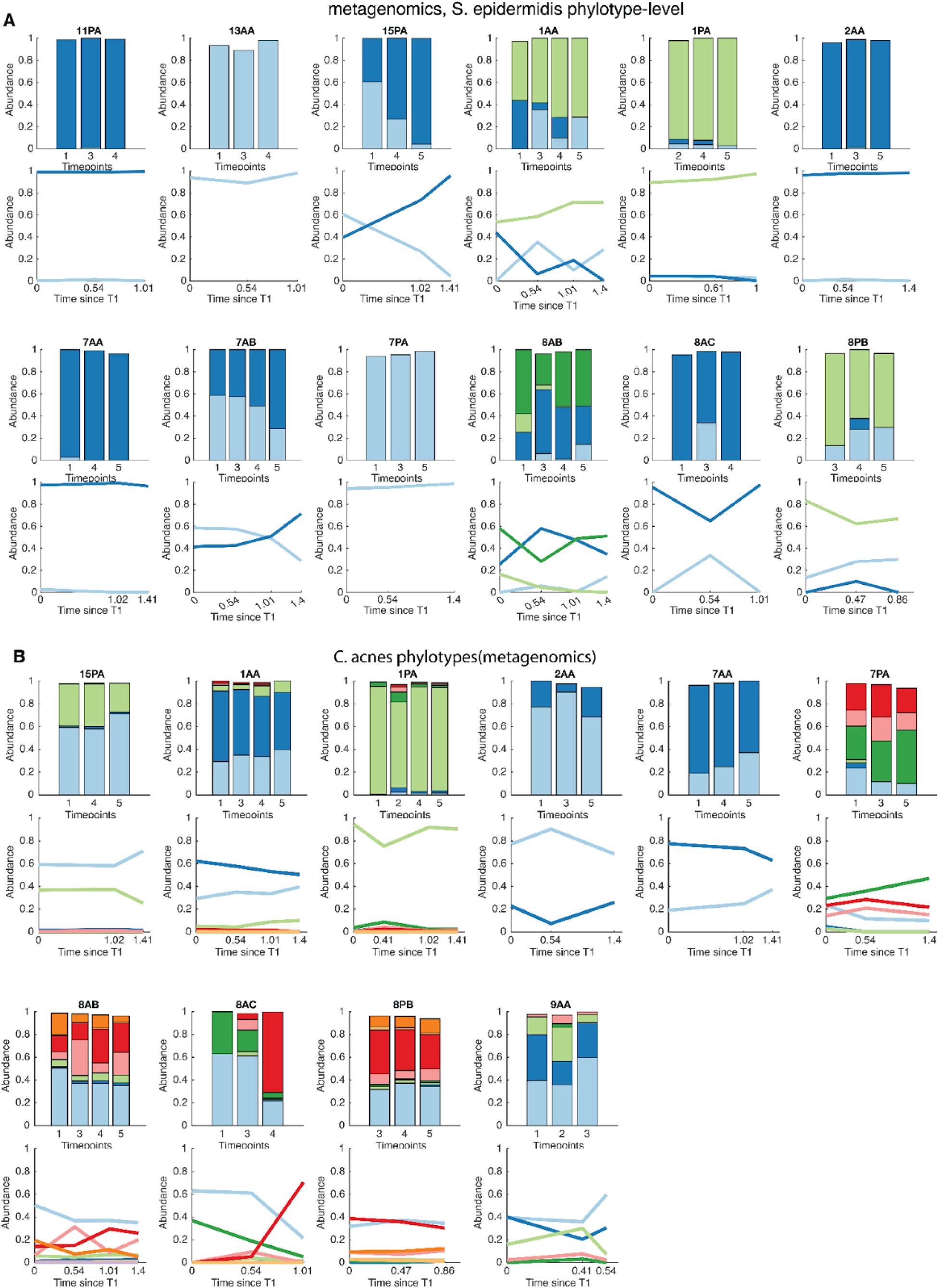
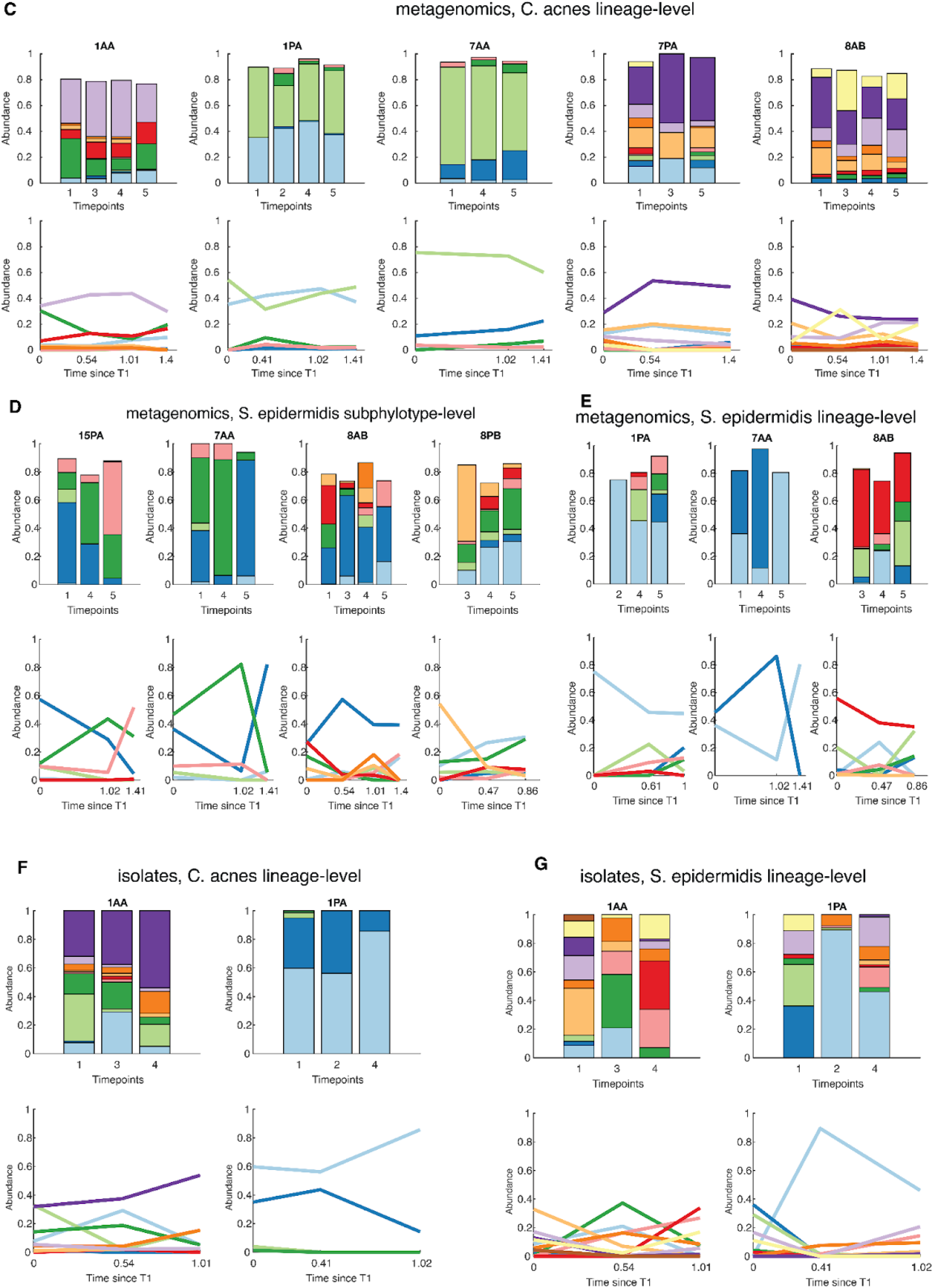
Longitudinal lineage-level dynamics for all subjects with at least 3 time points with sufficient data across different taxonomic levels. For each panel, the colors are arbitrary, and the same color across different subjects does not correspond to the same lineage. Each set of bars corresponds to the axes of line plots below it, which represent the relative abundance of each lineage or phylotype. **(A)** For *S. epidermidis* phylotypes (metagenomics), some clades are found at only one time point. (**B**) *C. acnes* phylotypes (metagenomics) exhibit high stability. **(C-E)** Lineage and sub-phylotype level data from metagenomics. **(F-G)** Lineage level data from isolates. See also Supplementary Tables 3 and 5 for complete lists of abundances.

**Figure S14.**
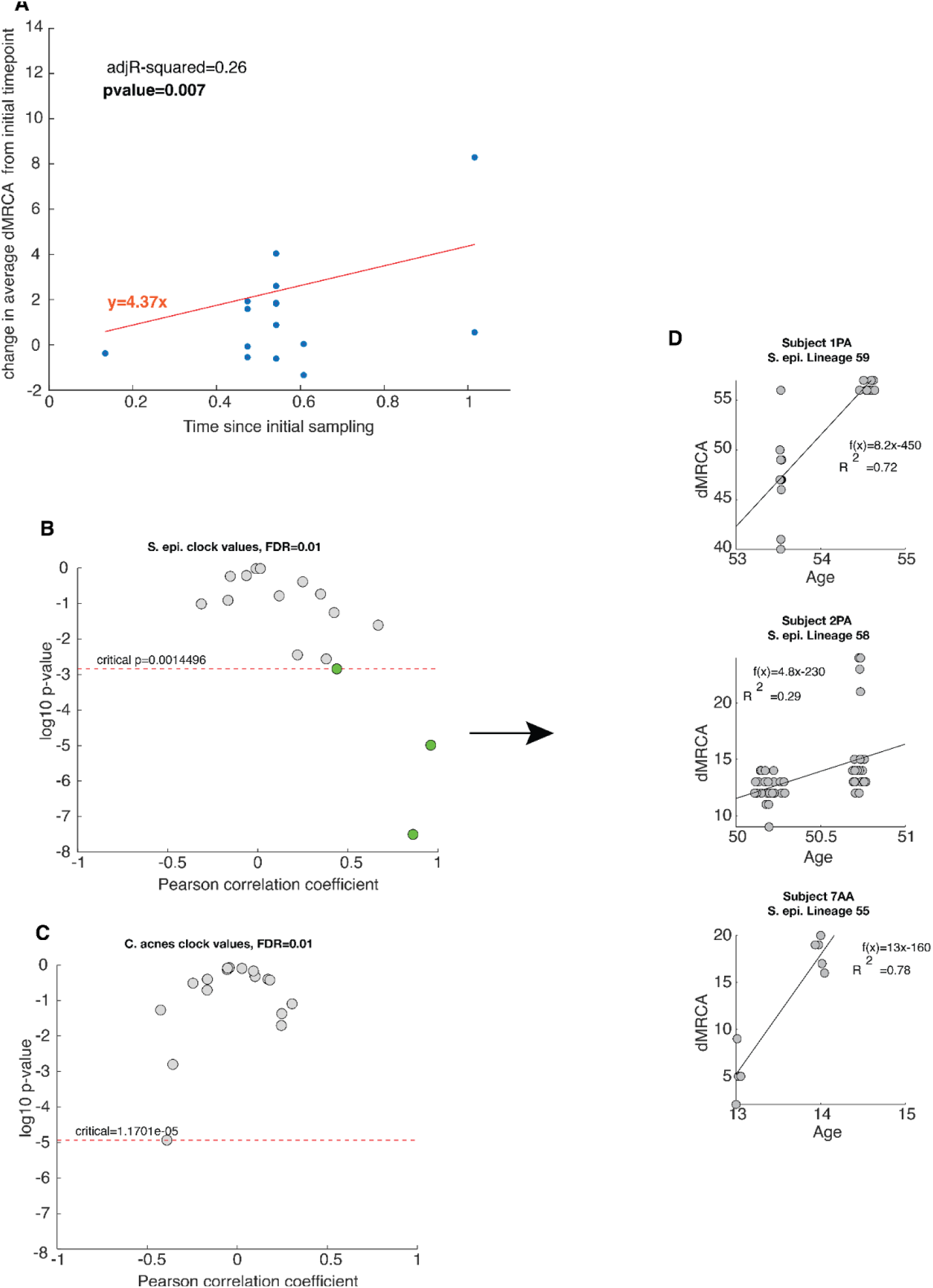
Molecular clock estimation. (A) For each instance where a subject had at least 5 *S. epidermidis* isolates in a given lineage at at least 2 time points, the change in average root-to-tip distance from isolates across time points was calculated and plotted against the interval between time points. The resulting clock value was 4.37 mutations per/genome/year. **(B-C)** When calculating molecular clocks for each lineage using root-to-tip (per isolate) versus time calculations, only three lineages of *S. epidermidis* received significant values. **(D)** Each of three lineages with significant molecular clock signals suggest faster molecular clocks, which would imply even higher rates of *S. epidermidis* dynamics.

**Figure S15.**
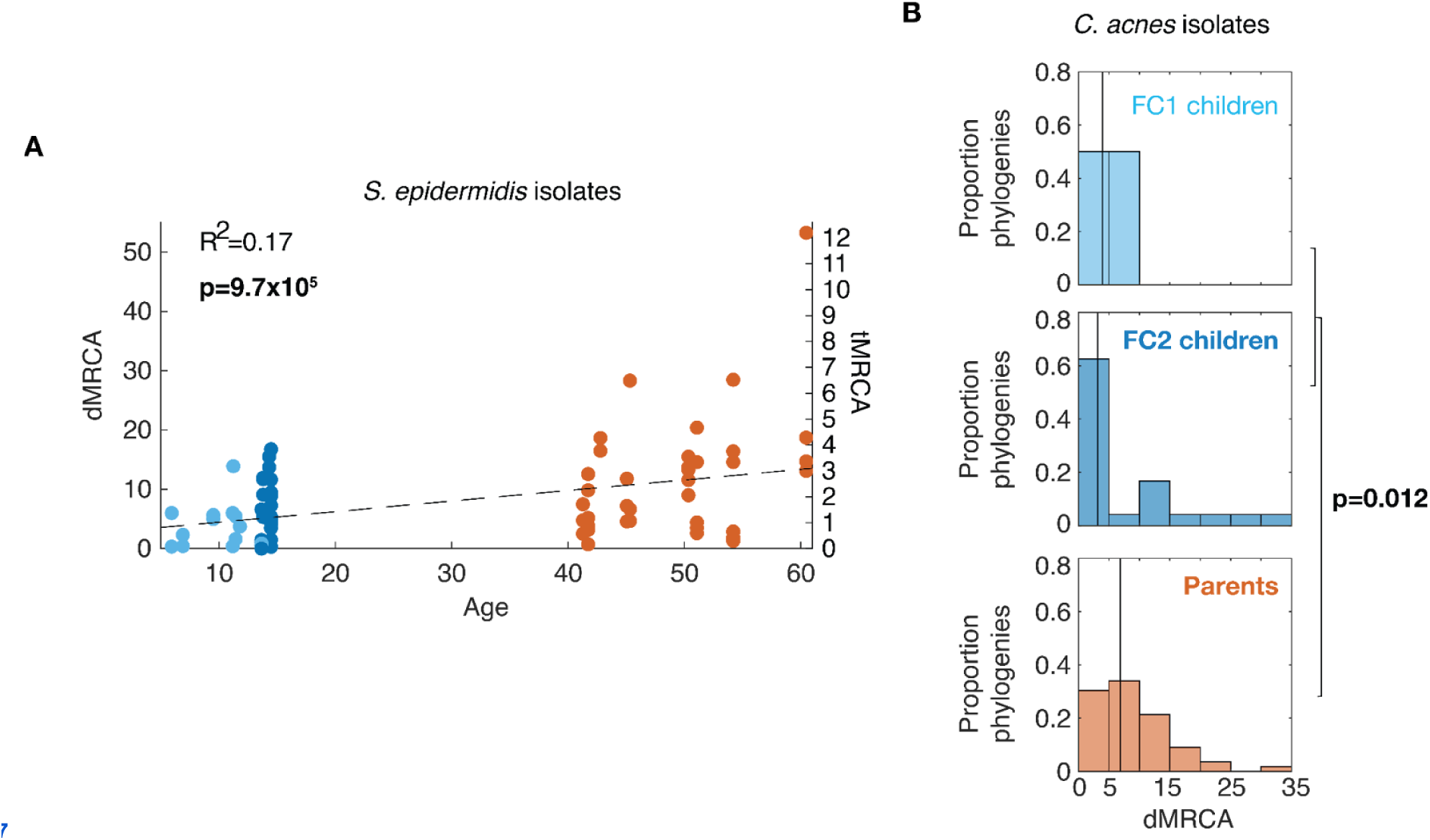
corresponding dMRCA analyses for *C. acnes* and *S. epidermidis*. **(A)** The subject dMRCAs of *S. epidermis* lineages are correlated with subject age (see Figure 6A for analogous analysis for *C. acnes*). **(B)** The subject dMRCAs of *C. acnes* lineages are higher in parents than children (see Figure 5D for analogous analysis for *S. epidermidis*). All correlation coefficients and P-values are for linear (Pearson) correlations or two-sided ranksum tests and are bolded if significant at a false discovery rate of 5% using the Benjamini-Hochberg procedure.

**Figure S16.**
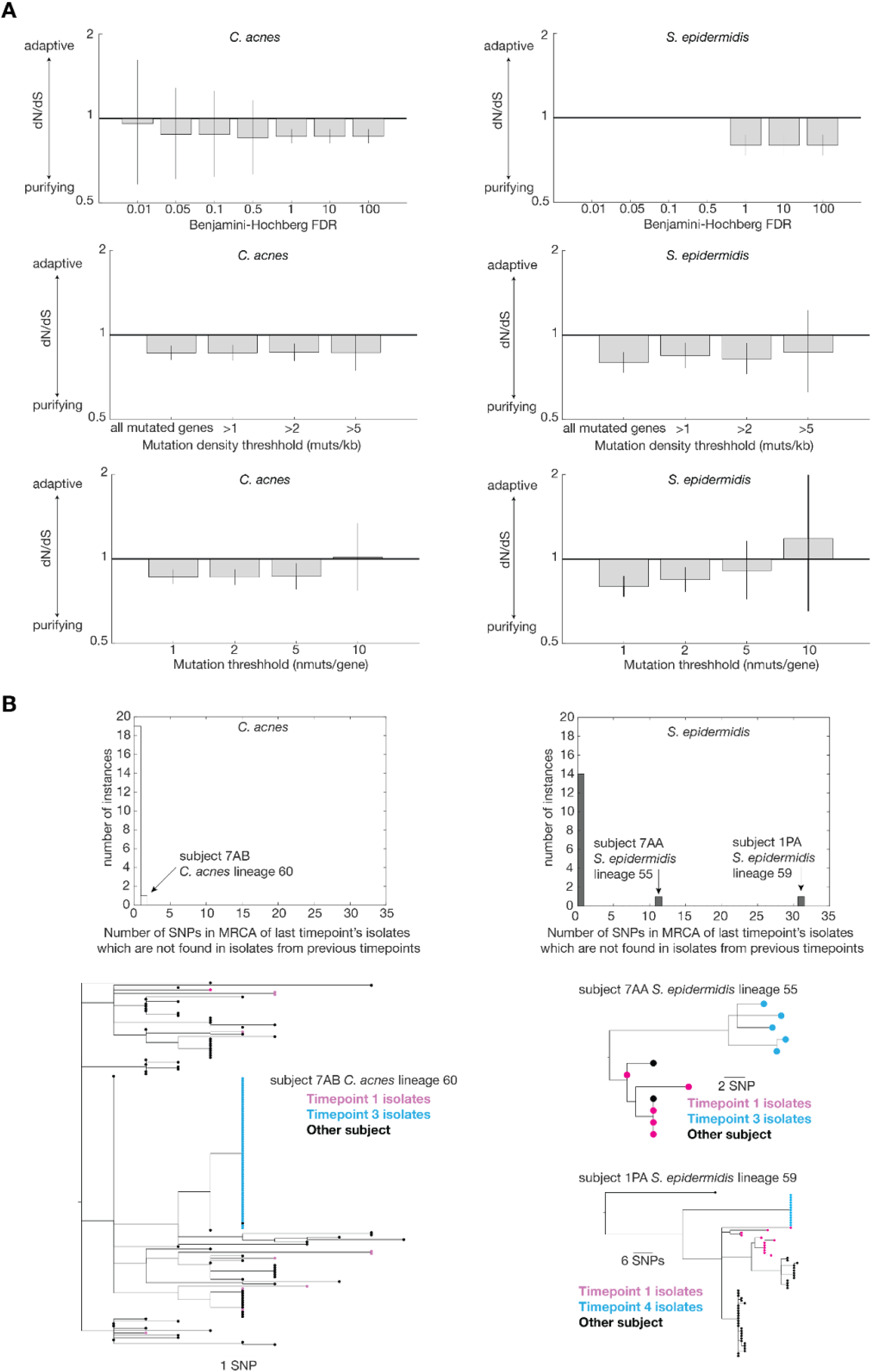
Within-lineage evolutionary dynamics are dominated by neutral forces. We performed tests for enrichment of *de novo* mutations (within-lineage mutations) for both species as in^16^ and detected no significant enrichment. **(A)** Correspondingly, we find no enrichment of nonsynonymous mutations as assessed by dN/dS for either species when considering the significance of the observed number of mutations (top), the density of mutations (middle), or the absolute number of mutations (bottom). **(B)** For instances where a subject had at least 5 isolates in a lineage at two timepoints, we also searched for evidence of sweeps by calculating the number of SNPs which are present in MRCA of all isolates of the latest time point, but are not fixed in the isolates from previous timepoints. Each non-zero instance for both species are shown. In each instance, the observed difference may be consistent with re-colonization of the lineage rather than a sweep, as the MRCA of the isolates observed at the last time point do not originate from within the diversity of the genotypes observed at initial timepoints. The isolates shown in black are from other subjects.

**Figure S17.**
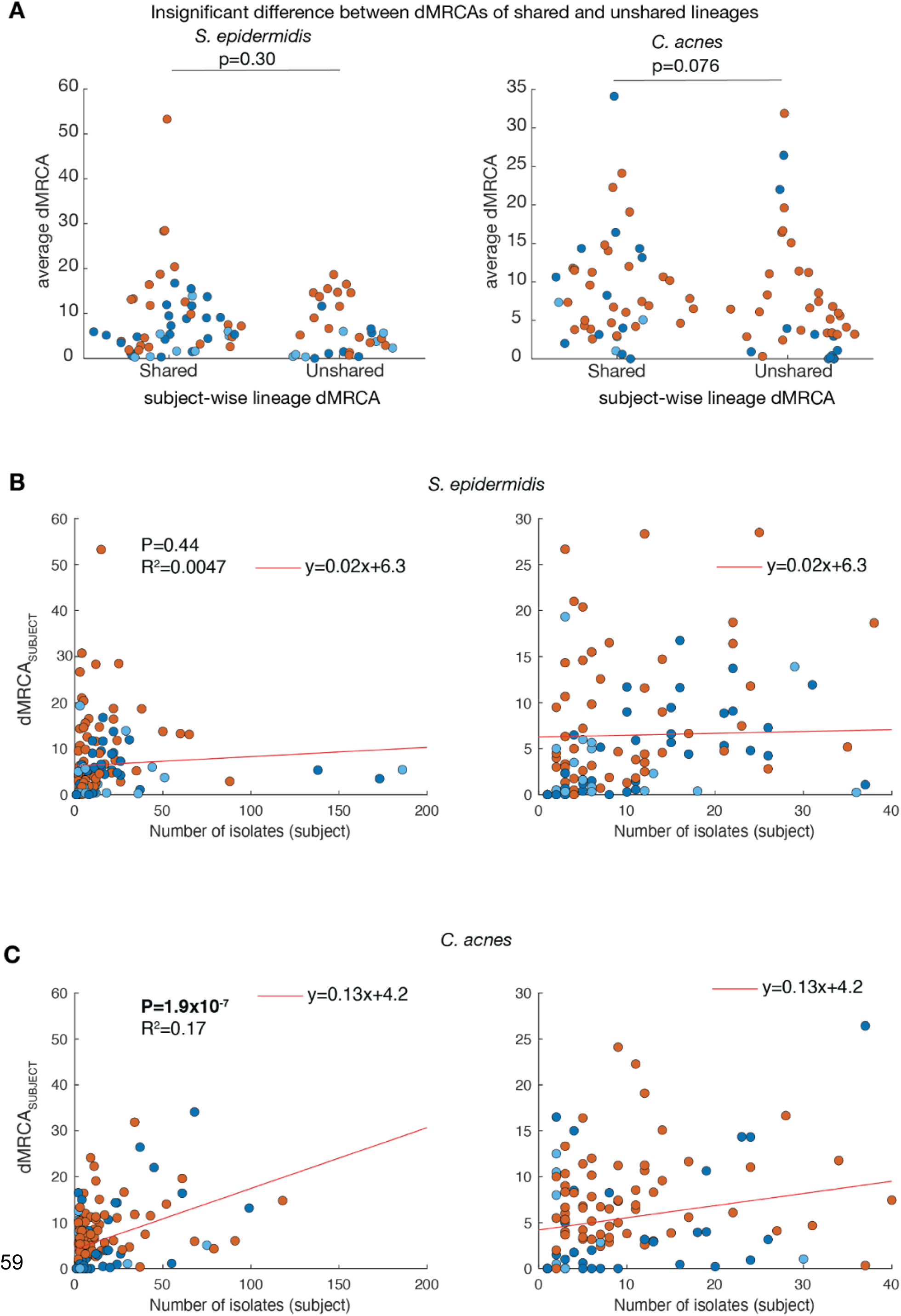
Shared and unshared individual dMRCAs. **(A)** The dMRCA_SUBJECT_ between shared and unshared lineages is not significantly different for either *C. acnes* and *S. epidermidis,* suggesting that lineages are not unshared as a consequence of insufficient time for sharing **(B-C)** For both species, dMRCA_SUBJECT_ is not substantially influenced by the number of collected isolates, indicating that lineages with fewer isolates do not significantly influence inferred dMRCAs.

**Figure S18.**
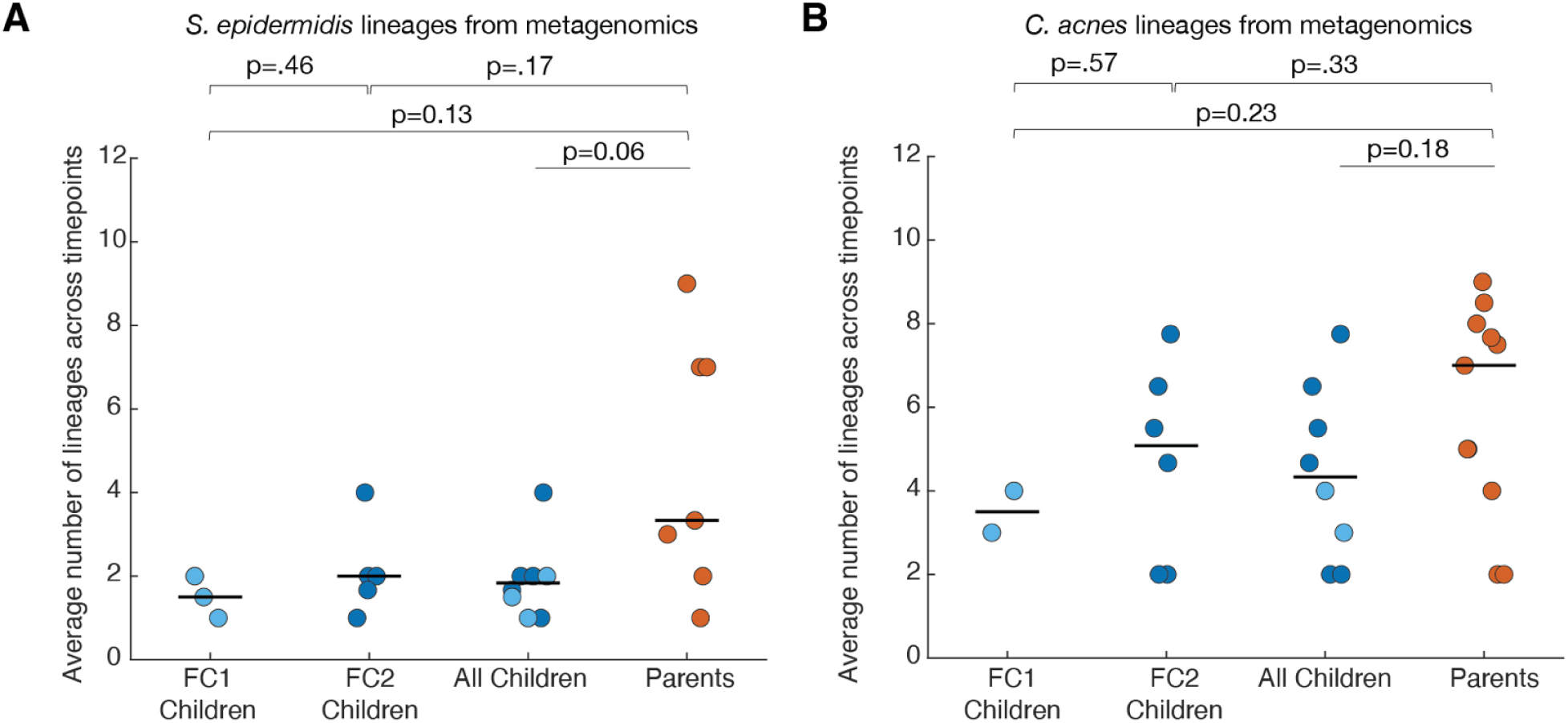
Lineages detected in subjects of different ages from metagenomics. (A-B) More *S. epidermidis* and *C. acnes* lineages, respectively, are found in parents than children, but not significantly. These trends support the analyses at the sub-phylotype level in Figures 5 and 6.

**Figure S19.**
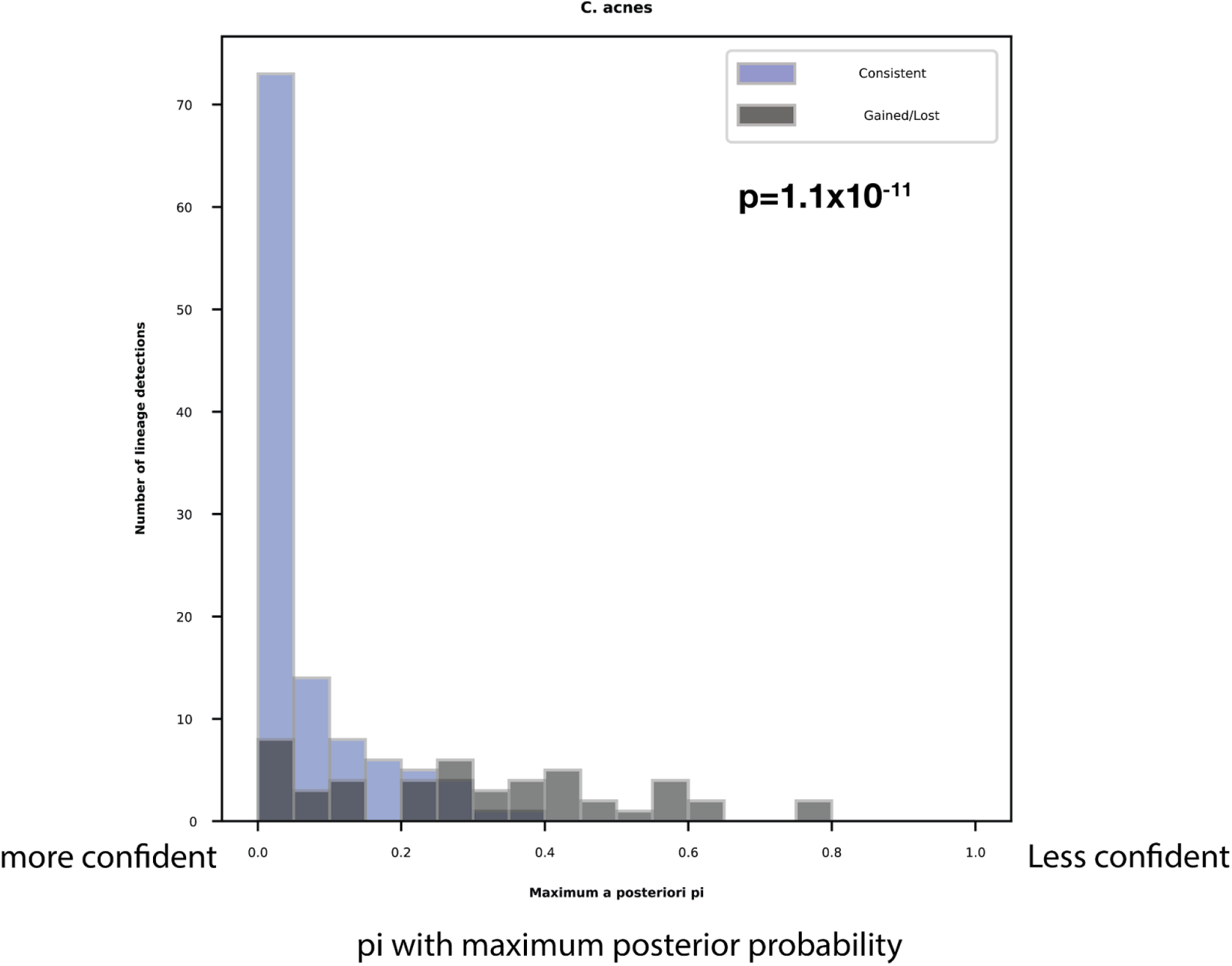
Confidence of stable versus unstable *C. acnes* lineages. Lineages which appear to be gained and lost between time points on the same subjects have a higher pi value (less confident) than lineages which are stable between time points, indicating that apparent gains and losses of low abundance *C. acnes* lineages can be noise.

**Figure S20.**
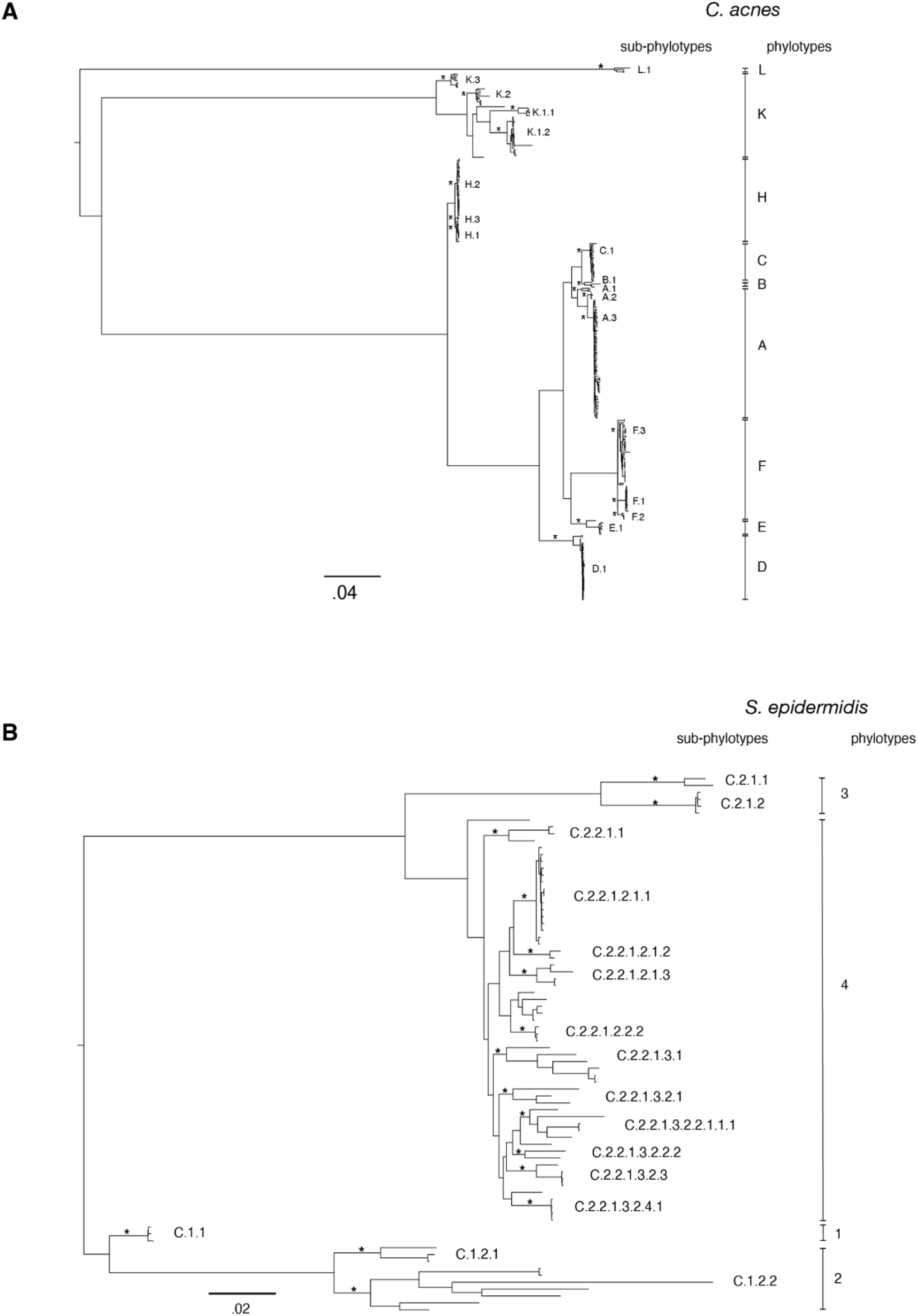
**Species-level phylogenies of *C. acnes* and *S. epidermidis.*** Maximum-likelihood phylogeny of **(A)** *C. acnes* and **(B)** *S. epidermidis* isolates with phylotypes and sub-phylotypes labeled. For both species, the isolate from each lineage with the highest coverage to the reference genome (Pacnes_C1 or SepidermidisATCC12228) was chosen as a representative. To ensure a wide collection of phylogenetic diversity for *C. acnes*, we also included a representative isolate from each of the 53 *C. acnes* lineages defined in Conwill et al^16^, as well as 197 publicly available *C. acnes* isolates from NCBI. Because the raw reads from some publicly available isolates were not available, we first simulated reads from these genomes using wgsim (v.0.3.1-r13; -e 0.0 -d 500 -N 500000 -1 150 -2 150 -r 0.0 -R 0.0 -X 0.0) before aligning all isolates to their respective reference genome (bowtie2 v.2.2.6; -X 2000 --no-mixed --dovetail). After alignment, samtools mpileup (v.1.5; -q30 -x -s -O -d3000) was used to identify candidate SNPs for each representative isolate. To filter out low-quality polymorphisms and positions across lineages, we first marked allele calls as ambiguous (“N”) in a sample if the major allele frequency was below 0.85, the FQ score produced by samtools was above -30, the coverage across either the forward or reverse reads was below 2, or if greater than 50% of reads at a position supported an indel. We then masked positions across samples if the median coverage across isolates was less than 3, or if more than 5% of isolates had an ambiguous allele at that position. We finally removed positions for which the correlation of non-reference alleles within 300bp of each other across isolates was above 0.75, which suggests that these alleles were acquired through recombination or other horizontal inheritance events. Positions which passed these filters and still contained at least one polymorphism across isolates were used to generate a maximum-likelihood phylogenetic tree under a GTR model of nucleotide substitution using RaXML (v. 8.2.12; -p 060782 -m GTRCAT).

**Figure S21.**
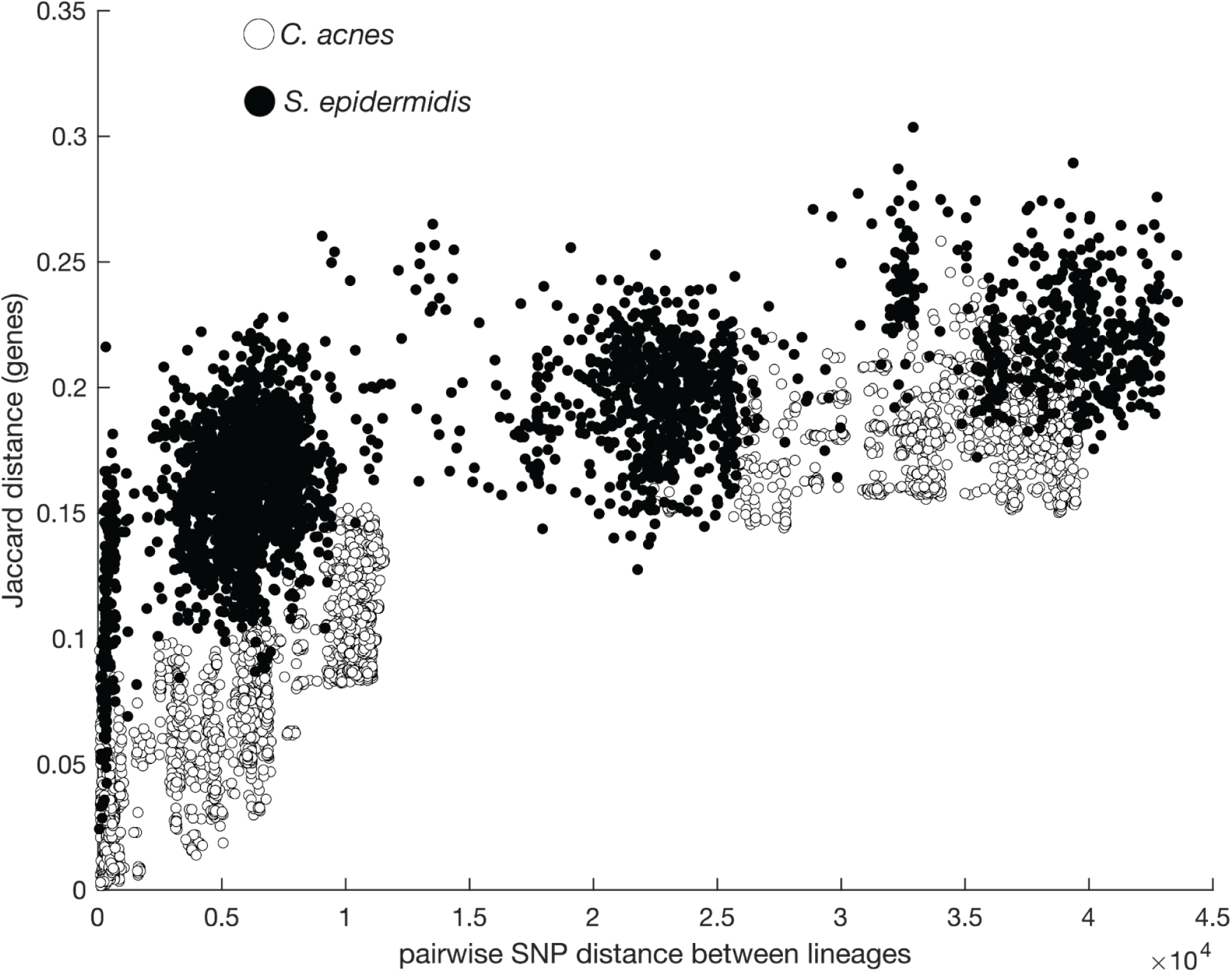
*S. epidermidis* has a more dynamic accessory genome than *C. acnes*. At low phylogenetic distances (x-axis), *S. epidermidis* lineages are more distinct at the gene-content level(y-axis) than *C. acnes*, highlighting higher rates of HGT or gene loss in *S. epidermidis*.

**Figure S22.**
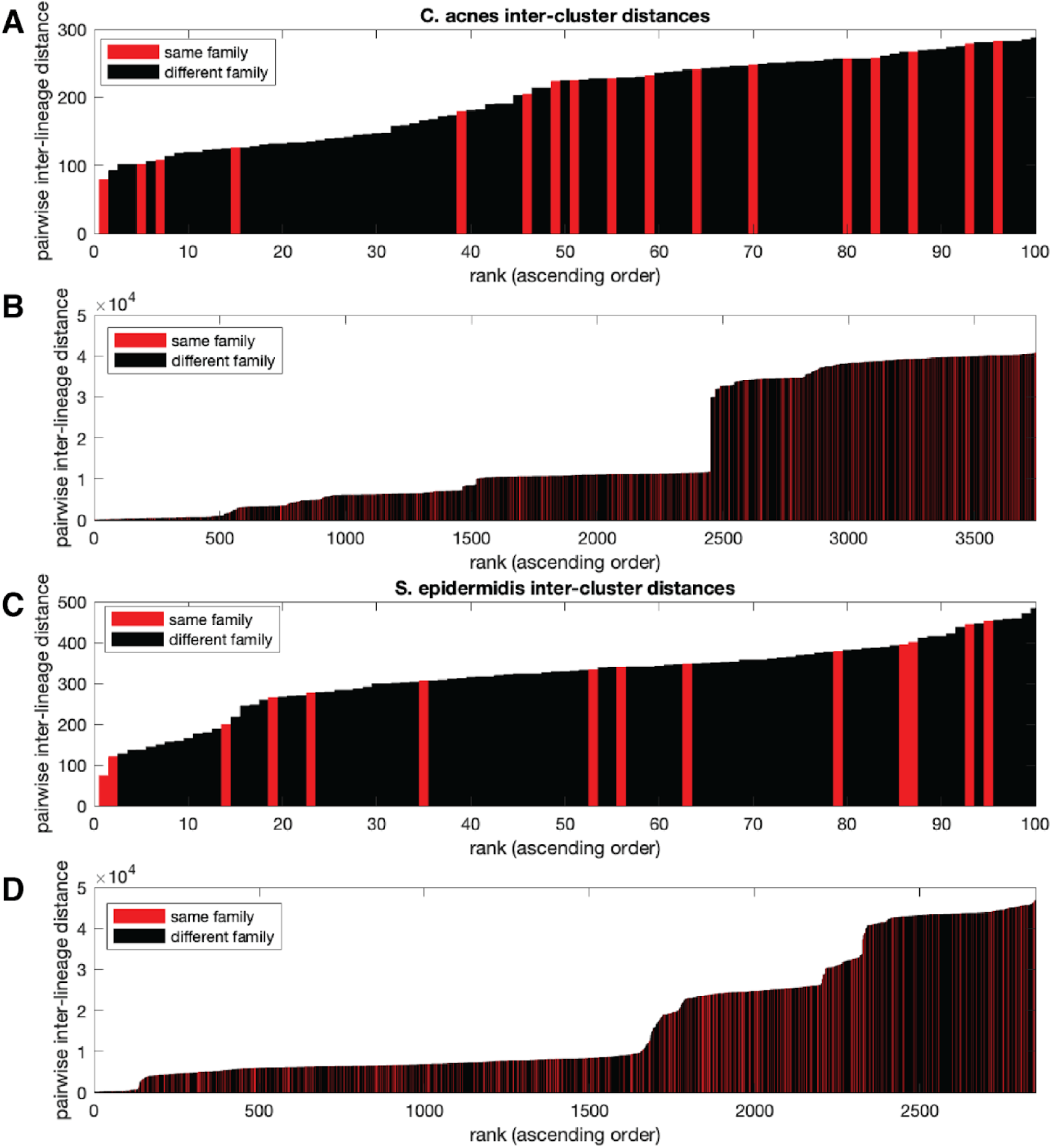
Lineages are not over-clustered. **(A)** For each pair of *C. acnes* lineages, their max pairwise distances were computed from reference genomes P_acnesC1 and SepidermidisATCC12228, respectively, and ranked in ascending order. Close together pairs are not more often from the same family, indicating that lineages are not over-clustered (first 100 are shown). **(B)** Same data as in (A) but expanded to include all pairs. **(C-D)** As in (A-B), but for *S. epidermidis*. For the two pairs of lineages in (C) and the one lineage in (A) which come from the same family (red) and are ranked the lowest, each pair of lineages come from different subjects, indicating that on-person diversity is not under-estimated.

**Figure S23.**
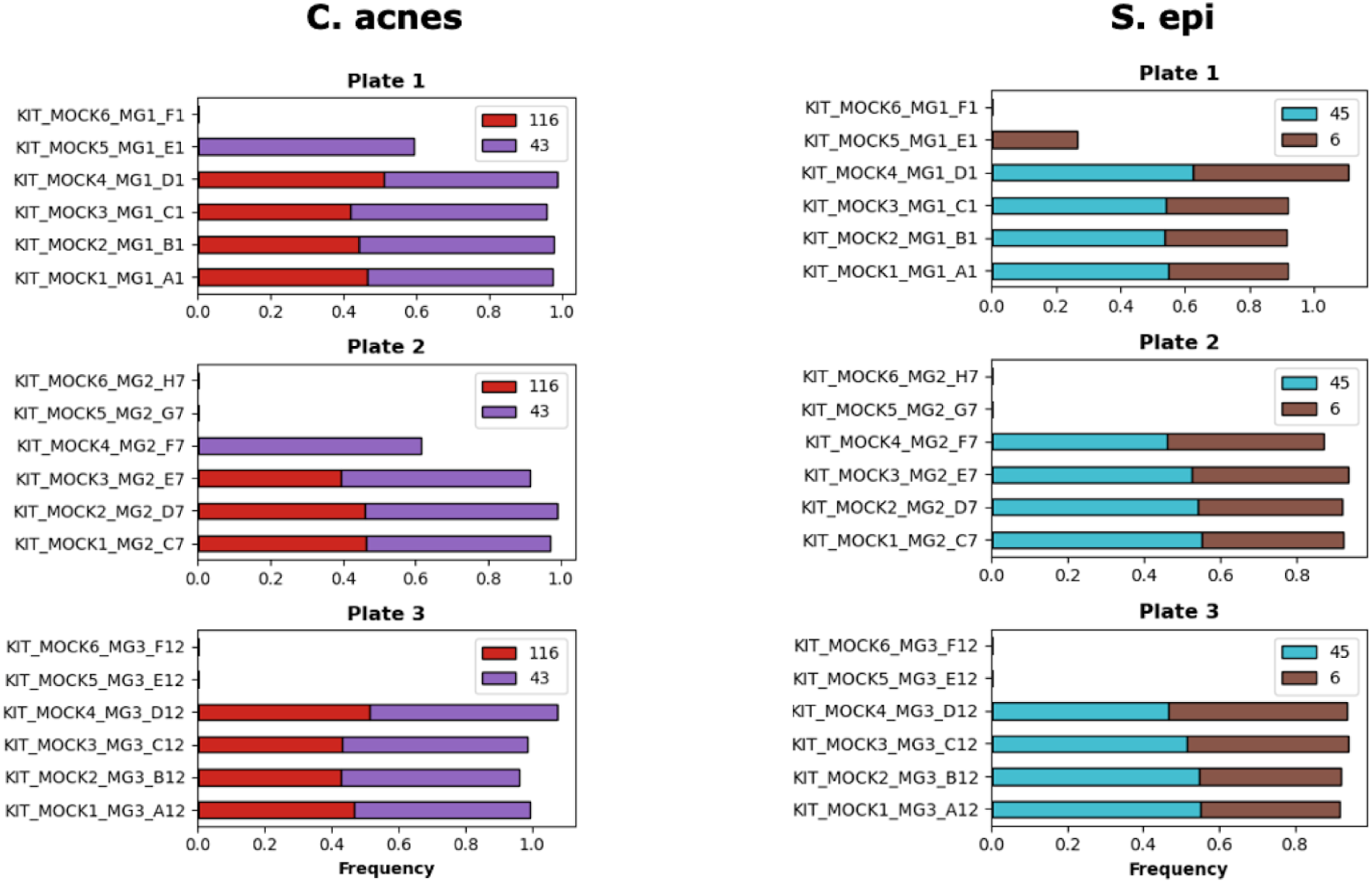
Mock communities. For both species, two lineages were mixed and used as part of a mock community on the plates MG1, MG2, and MG3. These high-abundance biomass samples cross contaminated some adjacent wells, so we excluded their abundances from calculation for all samples.

**Figure S24.**
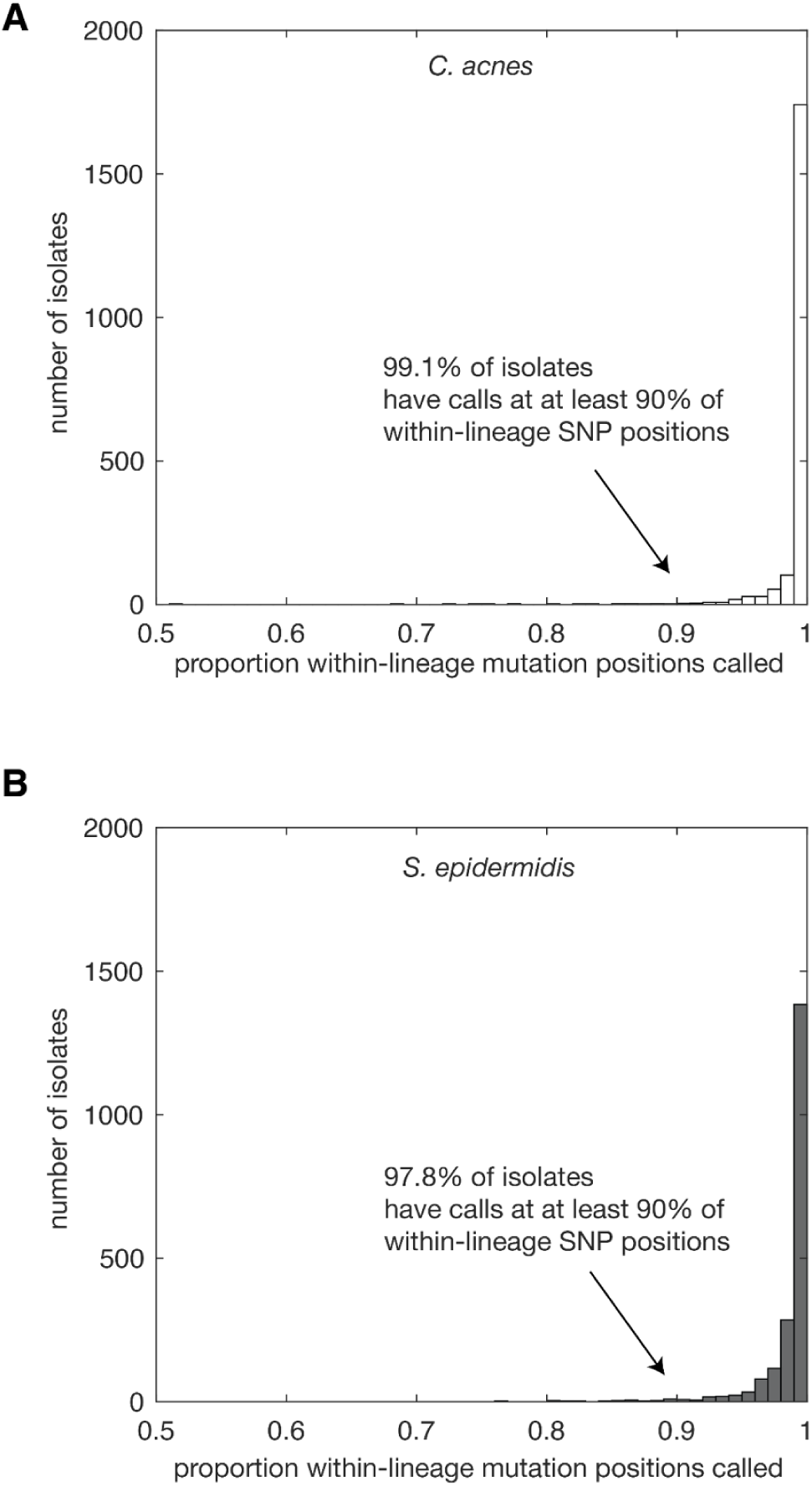
Proportion of non-ambiguous within-lineage SNP positions for each isolate. In clustered **(A)** *C. acnes* and **(B)** *S. epidermidis* isolates, the vast majority of isolates have nucleotide calls at each within-lineage SNP position. While ambiguous calls could come from cross contamination of closely related isolates not removed by our other filters, ambiguous calls are also expected from gene content found only in a subset of strains due to recent gain or loss (which is higher in *S. epidermidis*).

**Supplementary Tables 1-6**

Supplementary Table 1: subject_timepoint_metadata.xlsx

Supplementary Table 2: isolate_core_genome_filtering_CDHIT_homologues.xlsx

Supplementary Table 3: metagenomics_data.xlsx

Supplementary Table 4: lineage_MRCAs.xlsx

Supplementary Table 5: isolate_sample_names_lineage_membership.xlsx

Supplementary Table 6: SNPs.xlsx

## Notes

### Competing Interest Statement

The authors have declared no competing interest.

### Summary of Updates

We have revised this manuscript to clarify our methods throughout, improve the introduction and discussion, and add 6 additional Supplementary Figures.

## References

1. Costello, E.K., Lauber, C.L., Hamady, M., Fierer, N., Gordon, J.I., and Knight, R. (2009). Bacterial community variation in human body habitats across space and time. Science 326, 1694–1697.

2. Olm, M.R., Brown, C.T., Brooks, B., Firek, B., Baker, R., Burstein, D., Soenjoyo, K., Thomas, B.C., Morowitz, M., and Banfield, J.F. (2017). Identical bacterial populations colonize premature infant gut, skin, and oral microbiomes and exhibit different in situ growth rates. Genome Res. 27, 601–612.

3. Harris-Tryon, T.A., and Grice, E.A. (2022). Microbiota and maintenance of skin barrier function. Science 376, 940–945.

4. Oh, J., Byrd, A.L., Deming, C., Conlan, S., NISC Comparative Sequencing Program, Kong, H.H., and Segre, J.A. (2014). Biogeography and individuality shape function in the human skin metagenome. Nature 514, 59–64.

5. Oh, J., Byrd, A.L., Park, M., NISC Comparative Sequencing Program, Kong, H.H., and Segre, J.A. (2016). Temporal Stability of the Human Skin Microbiome. Cell 165, 854–866.

6. Human Microbiome Project Consortium (2012). Structure, function and diversity of the healthy human microbiome. Nature 486, 207–214.

7. Van Rossum, T., Ferretti, P., Maistrenko, O.M., and Bork, P. (2020). Diversity within species: interpreting strains in microbiomes. Nat. Rev. Microbiol. 18, 491–506.

8. Lieberman, T.D. (2022). Detecting bacterial adaptation within individual microbiomes. Philos. Trans. R. Soc. Lond. B Biol. Sci. 377, 20210243.

9. Paetzold, B., Willis, J.R., Pereira de Lima, J., Knödlseder, N., Brüggemann, H., Quist, S.R., Gabaldón, T., and Güell, M. (2019). Skin microbiome modulation induced by probiotic solutions. Microbiome 7, 95.

10. Koskella, B., Hall, L.J., and Metcalf, C.J.E. (2017). The microbiome beyond the horizon of ecological and evolutionary theory. Nat. Ecol. Evol. 1, 1606–1615.

11. Chen, D.W., and Garud, N.R. (2022). Rapid evolution and strain turnover in the infant gut microbiome. Genome Res. 32, 1124–1136.

12. Schmidt, T.S.B., Raes, J., and Bork, P. (2018). The Human Gut Microbiome: From Association to Modulation. Cell 172, 1198–1215.

13. Costello, E.K., Stagaman, K., Dethlefsen, L., Bohannan, B.J.M., and Relman, D.A. (2012). The application of ecological theory toward an understanding of the human microbiome. Science 336, 1255–1262.

14. Garud, N.R., and Pollard, K.S. (2020). Population genetics in the human microbiome. Trends Genet. 36, 53–67.

15. Shen, Z., Robert, L., Stolpman, M., Che, Y., Allen, K.J., Saffery, R., Walsh, A., Young, A., Eckert, J., Deming, C., et al. (2023). A genome catalog of the early-life human skin microbiome. Genome Biol. 24, 252.

16. Conwill, A., Kuan, A.C., Damerla, R., Poret, A.J., Baker, J.S., Tripp, A.D., Alm, E.J., and Lieberman, T.D. (2022). Anatomy promotes neutral coexistence of strains in the human skin microbiome. Cell Host Microbe 30, 171–182.e7.

17. Key, F.M., Khadka, V.D., Romo-González, C., Blake, K.J., Deng, L., Lynn, T.C., Lee, J.C., Chiu, I.M., García-Romero, M.T., and Lieberman, T.D. (2023). On-person adaptive evolution of Staphylococcus aureus during treatment for atopic dermatitis. Cell Host Microbe 31, 593–603.e7.

18. Hildebrand, F., Gossmann, T.I., Frioux, C., Özkurt, E., Myers, P.N., Ferretti, P., Kuhn, M., Bahram, M., Nielsen, H.B., and Bork, P. (2021). Dispersal strategies shape persistence and evolution of human gut bacteria. Cell Host Microbe 29, 1167–1176.e9.

19. Xue, K.S., Walton, S.J., Goldman, D.A., Morrison, M.L., Verster, A.J., Parrott, A.B., Yu, F.B., Neff, N.F., Rosenberg, N.A., Ross, B.D., et al. (2023). Prolonged delays in human microbiota transmission after a controlled antibiotic perturbation. bioRxiv. 10.1101/2023.09.26.559480.

20. Valles-Colomer, M., Blanco-Míguez, A., Manghi, P., Asnicar, F., Dubois, L., Golzato, D., Armanini, F., Cumbo, F., Huang, K.D., Manara, S., et al. (2023). The person-to-person transmission landscape of the gut and oral microbiomes. Nature 614, 125–135.

21. Lou, Y.C., Olm, M.R., Diamond, S., Crits-Christoph, A., Firek, B.A., Baker, R., Morowitz, M.J., and Banfield, J.F. (2021). Infant gut strain persistence is associated with maternal origin, phylogeny, and traits including surface adhesion and iron acquisition. Cell Rep Med 2, 100393.

22. Yassour, M., Jason, E., Hogstrom, L.J., Arthur, T.D., Tripathi, S., Siljander, H., Selvenius, J., Oikarinen, S., Hyöty, H., Virtanen, S.M., et al. (2018). Strain-Level Analysis of Mother-to-Child Bacterial Transmission during the First Few Months of Life. Cell Host Microbe 24, 146–154.e4.

23. Nayfach, S., Rodriguez-Mueller, B., Garud, N., and Pollard, K.S. (2016). An integrated metagenomics pipeline for strain profiling reveals novel patterns of bacterial transmission and biogeography. Genome Res. 26, 1612–1625.

24. Pochi, P.E., Strauss, J.S., and Downing, D.T. (1979). Age-related changes in sebaceous gland activity. J. Invest. Dermatol. 73, 108–111.

25. Oh, J., Conlan, S., Polley, E.C., Segre, J.A., and Kong, H.H. (2012). Shifts in human skin and nares microbiota of healthy children and adults. Genome Med. 4, 77.

26. Park, J., Schwardt, N.H., Jo, J.-H., Zhang, Z., Pillai, V., Phang, S., Brady, S.M., Portillo, J.A., MacGibeny, M.A., Liang, H., et al. (2022). Shifts in the Skin Bacterial and Fungal Communities of Healthy Children Transitioning through Puberty. J. Invest. Dermatol. 142, 212–219.

27. Schneider, A.M., Nolan, Z.T., Banerjee, K., Paine, A.R., Cong, Z., Gettle, S.L., Longenecker, A.L., Zhan, X., Agak, G.W., and Nelson, A.M. (2023). Evolution of the facial skin microbiome during puberty in normal and acne skin. J. Eur. Acad. Dermatol. Venereol. 37, 166–175.

28. Leyden, J.J., McGinley, K.J., Mills, O.H., and Kligman, A.M. (1975). Age-related changes in the resident bacterial flora of the human face. J. Invest. Dermatol. 65, 379–381.

29. Acosta, E.M., Little, K.A., Bratton, B.P., Lopez, J.G., Mao, X., Payne, A.S., Donia, M., Devenport, D., and Gitai, Z. (2023). Bacterial DNA on the skin surface overrepresents the viable skin microbiome. Elife 12. 10.7554/eLife.87192.

30. Zhou, W., Spoto, M., Hardy, R., Guan, C., Fleming, E., Larson, P.J., Brown, J.S., and Oh, J. (2020). Host-Specific Evolutionary and Transmission Dynamics Shape the Functional Diversification of Staphylococcus epidermidis in Human Skin. Cell 180, 454–470.e18.

31. Hubbell, S.P. (2001). The Unified Neutral Theory of Biodiversity and Biogeography (Princeton University Press).

32. Debray, R., Herbert, R.A., Jaffe, A.L., Crits-Christoph, A., Power, M.E., and Koskella, B. (2022). Priority effects in microbiome assembly. Nat. Rev. Microbiol. 20, 109–121.

33. Nübel, U., Nachtnebel, M., Falkenhorst, G., Benzler, J., Hecht, J., Kube, M., Bröcker, F., Moelling, K., Bührer, C., Gastmeier, P., et al. (2013). MRSA transmission on a neonatal intensive care unit: epidemiological and genome-based phylogenetic analyses. PLoS One 8, e54898.

34. Young, B.C., Golubchik, T., Batty, E.M., Fung, R., Larner-Svensson, H., Votintseva, A.A., Miller, R.R., Godwin, H., Knox, K., Everitt, R.G., et al. (2012). Evolutionary dynamics of *Staphylococcus aureus* during progression from carriage to disease. Proc. Natl. Acad. Sci. U. S. A. 109, 4550–4555.

35. Golubchik, T., Batty, E.M., Miller, R.R., Farr, H., Young, B.C., Larner-Svensson, H., Fung, R., Godwin, H., Knox, K., Votintseva, A., et al. (2013). Within-host evolution of Staphylococcus aureus during asymptomatic carriage. PLoS One 8, e61319.

36. Karoglan, A., Paetzold, B., Pereira de Lima, J., Brüggemann, H., Tüting, T., Schanze, D., Güell, M., and Gollnick, H. (2019). Safety and efficacy of topically applied selected Cutibacterium acnes strains over five weeks in patients with acne vulgaris: An open-label, pilot study. Acta Derm. Venereol. 99, 1253–1257.

37. Dubois, L., Valles-Colomer, M., Ponsero, A., Helve, O., Andersson, S., Kolho, K.-L., Asnicar, F., Korpela, K., Salonen, A., Segata, N., et al. (2024). Paternal and induced gut microbiota seeding complement mother-to-infant transmission. Cell Host Microbe 32, 1011–1024.e4.

38. Brito, I.L., Gurry, T., Zhao, S., Huang, K., Young, S.K., Shea, T.P., Naisilisili, W., Jenkins, A.P., Jupiter, S.D., Gevers, D., et al. (2019). Transmission of human-associated microbiota along family and social networks. Nat. Microbiol. 4, 964–971.

39. Kageyama, S., Furuta, M., Takeshita, T., Ma, J., Asakawa, M., and Yamashita, Y. (2022). High-Level acquisition of maternal oral bacteria in formula-fed infant oral Microbiota. MBio 13, e0345221.

40. Kim, H.J., Boedicker, J.Q., Choi, J.W., and Ismagilov, R.F. (2008). Defined spatial structure stabilizes a synthetic multispecies bacterial community. Proc. Natl. Acad. Sci. U. S. A. 105, 18188–18193.

41. Tilman, D. (1994). Competition and biodiversity in spatially structured habitats. Ecology 75, 2–16.

42. Pisa, H., Hermisson, J., and Polechová, J. (2019). The influence of fluctuating population densities on evolutionary dynamics. Evolution 73, 1341–1355.

43. Ahle, C.M., Stødkilde, K., Poehlein, A., Bömeke, M., Streit, W.R., Wenck, H., Reuter, J.H., Hüpeden, J., and Brüggemann, H. (2022). Interference and co-existence of staphylococci and Cutibacterium acnes within the healthy human skin microbiome. Commun Biol 5, 923.

44. Claesen, J., Spagnolo, J.B., Ramos, S.F., Kurita, K.L., Byrd, A.L., Aksenov, A.A., Melnik, A.V., Wong, W.R., Wang, S., Hernandez, R.D., et al. (2020). A Cutibacterium acnes antibiotic modulates human skin microbiota composition in hair follicles. Sci. Transl. Med. 12. 10.1126/scitranslmed.aay5445.

45. Bomar, L., Brugger, S.D., Yost, B.H., Davies, S.S., and Lemon, K.P. (2016). Corynebacterium accolens releases antipneumococcal free fatty acids from human nostril and skin surface triacylglycerols. MBio 7, e01725–15.

46. Christensen, G.J.M., Scholz, C.F.P., Enghild, J., Rohde, H., Kilian, M., Thürmer, A., Brzuszkiewicz, E., Lomholt, H.B., and Brüggemann, H. (2016). Antagonism between Staphylococcus epidermidis and Propionibacterium acnes and its genomic basis. BMC Genomics 17, 152.

47. Gless, B.H., Bojer, M.S., Peng, P., Baldry, M., Ingmer, H., and Olsen, C.A. (2019). Identification of autoinducing thiodepsipeptides from staphylococci enabled by native chemical ligation. Nat. Chem. 11, 463–469.

48. Mancuso, C.P., Baker, J.S., Qu, E., Tripp, A.D., Balogun, I.O., and Lieberman, T.D. (2024). Intraspecies warfare restricts strain coexistence in human skin microbiomes. bioRxiv. 10.1101/2024.05.07.592803.

49. Lima, R.D., Dos Reis, G.A., da Silva Reviello, J., Glatthardt, T., da Silva Coimbra, L., Lima, C.O.G.X., Antunes, L.C.M., and Ferreira, R.B.R. (2021). Antibiofilm activity of Cutibacterium acnes cell-free conditioned media against Staphylococcus spp. Braz. J. Microbiol. 52, 2373–2383.

50. Guan, C., Larson, P.J., Fleming, E., Tikhonov, A.P., Mootien, S., Grossman, T.H., Golino, C., and Oh, J. (2022). Engineering a “detect and destroy” skin probiotic to combat methicillin-resistant Staphylococcus aureus. PLoS One 17, e0276795.

51. Bek-Thomsen, M., Lomholt, H.B., and Kilian, M. (2008). Acne is Not Associated with Yet-Uncultured Bacteria. J. Clin. Microbiol. 46, 3355–3360.

52. Fitz-Gibbon, S., Tomida, S., Chiu, B.-H., Nguyen, L., Du, C., Liu, M., Elashoff, D., Erfe, M.C., Loncaric, A., Kim, J., et al. (2013). Propionibacterium acnes strain populations in the human skin microbiome associated with acne. J. Invest. Dermatol. 133, 2152–2160.

53. Tett, A., Pasolli, E., Farina, S., Truong, D.T., Asnicar, F., Zolfo, M., Beghini, F., Armanini, F., Jousson, O., De Sanctis, V., et al. (2017). Unexplored diversity and strain-level structure of the skin microbiome associated with psoriasis. NPJ Biofilms Microbiomes 3, 14.

54. Chang, H.-W., Yan, D., Singh, R., Liu, J., Lu, X., Ucmak, D., Lee, K., Afifi, L., Fadrosh, D., Leech, J., et al. (2018). Alteration of the cutaneous microbiome in psoriasis and potential role in Th17 polarization. Microbiome 6, 154.

55. Kong, H.H., Oh, J., Deming, C., Conlan, S., Grice, E.A., Beatson, M.A., Nomicos, E., Polley, E.C., Komarow, H.D., NISC Comparative Sequence Program, et al. (2012). Temporal shifts in the skin microbiome associated with disease flares and treatment in children with atopic dermatitis. Genome Res. 22, 850–859.

56. Lomholt, H.B., and Kilian, M. (2010). Population genetic analysis of Propionibacterium acnes identifies a subpopulation and epidemic clones associated with acne. PLoS One 5, e12277.

57. Brüggemann, H., Salar-Vidal, L., Gollnick, H.P.M., and Lood, R. (2021). A Janus-Faced Bacterium: Host-Beneficial and -Detrimental Roles of Cutibacterium acnes. Front. Microbiol. 12, 673845.

58. Rozas, M., Hart de Ruijter, A., Fabrega, M.J., Zorgani, A., Guell, M., Paetzold, B., and Brillet, F. (2021). From dysbiosis to healthy skin: Major contributions of Cutibacterium acnes to skin homeostasis. Microorganisms 9, 628.

59. Gaio, D., Anantanawat, K., To, J., Liu, M., Monahan, L., and Darling, A.E. (2022). Hackflex: low-cost, high-throughput, Illumina Nextera Flex library construction. Microb Genom 8. 10.1099/mgen.0.000744.

60. BMap (2022). SourceForge. https://sourceforge.net/projects/bbmap/.

61. Wood, D.E., Lu, J., and Langmead, B. (2019). Improved metagenomic analysis with Kraken 2. Genome Biol. 20, 257.

62. Lu, J., Breitwieser, F.P., Thielen, P., and Salzberg, S.L. (2017). Bracken: estimating species abundance in metagenomics data. PeerJ Comput. Sci. 3, e104.

63. Bankevich, A., Nurk, S., Antipov, D., Gurevich, A.A., Dvorkin, M., Kulikov, A.S., Lesin, V.M., Nikolenko, S.I., Pham, S., Prjibelski, A.D., et al. (2012). SPAdes: a new genome assembly algorithm and its applications to single-cell sequencing. J. Comput. Biol. 19, 455–477.

64. Seemann, T. (2014). Prokka: rapid prokaryotic genome annotation. Bioinformatics 30, 2068–2069.

65. Li, W., and Godzik, A. (2006). Cd-hit: a fast program for clustering and comparing large sets of protein or nucleotide sequences. Bioinformatics 22, 1658–1659.

66. Dunn†, J.C. (1974). Well-separated clusters and optimal fuzzy partitions. J. Cybern. 4, 95–104.

67. Kleinberg, J. (2002). An impossibility theorem for clustering. In Proceedings of the 15th International Conference on Neural Information Processing Systems NIPS’02. (MIT Press), pp. 463–470.

68. Watts, D.J., and Strogatz, S.H. (1998). Collective dynamics of “small-world” networks. Nature 393, 440–442.

69. Ester, M., Kriegel, H.-P., Sander, J., and Xu, X. (1996). A density-based algorithm for discovering clusters in large spatial databases with noise. In Proceedings of the Second International Conference on Knowledge Discovery and Data Mining KDD’96. (AAAI Press), pp. 226–231.

70. phlame (Github).

71. Scholz, C.F.P., Jensen, A., Lomholt, H.B., Brüggemann, H., and Kilian, M. (2014). A novel high-resolution single locus sequence typing scheme for mixed populations of Propionibacterium acnes in vivo. PLoS One 9, e104199.

72. Schwengers, O., Jelonek, L., Dieckmann, M.A., Beyvers, S., Blom, J., and Goesmann, A. (2021). Bakta: rapid and standardized annotation of bacterial genomes via alignment-free sequence identification. Microb Genom 7. 10.1099/mgen.0.000685.

73. Page, A.J., Cummins, C.A., Hunt, M., Wong, V.K., Reuter, S., Holden, M.T.G., Fookes, M., Falush, D., Keane, J.A., and Parkhill, J. (2015). Roary: rapid large-scale prokaryote pan genome analysis. Bioinformatics 31, 3691–3693.

74. Lieberman, T.D., Wilson, D., Misra, R., Xiong, L.L., Moodley, P., Cohen, T., and Kishony, R. (2016). Genomic diversity in autopsy samples reveals within-host dissemination of HIV-associated Mycobacterium tuberculosis. Nat. Med. 22, 1470–1474.

